# KRAS-Mediated CCDC6 Degradation Drives xCT Upregulation and Ferroptosis Evasion

**DOI:** 10.64898/2026.05.05.721389

**Authors:** Daniela Criscuolo, Rosaria Catalano, Carmela Baviello, Claudia Fioravanti, Elena Vigliar, Francesco Morra, Maria Marotta, Junsei Mimura, Antonino Iaccarino, Francesco Pepe, Dorina Belotti, Giancarlo Troncone, Francesco Merolla, Rosa Marina Melillo, Angela Celetti

**Author notes:** Correspondence should be sent to: Angela Celetti, MD, PhD, CNR – Institute of Endotypes in Oncology, Metabolism and Immunology “G Salvatore” (IEOMI) Via Pansini, 5, Naples, Italy. Rosa Marina Melillo and Angela Celetti contributed equally as senior authors.

## Abstract

Oncogenic KRAS mutations drive tumorigenesis by promoting pro-survival signaling and metabolic reprogramming, including the maintenance of redox balance to evade oxidative stress. A key mechanism involves the upregulation of the xCT cystine/glutamate antiporter, which sustains glutathione (GSH) synthesis and protects cells from oxidative damage and ferroptosis. While it is known that the ETS1-ATF4 complex mediates transcriptional upregulation of xCT, the upstream regulators linking KRAS signaling to this axis remain to be fully defined.

Here, we demonstrate that oncogenic KRAS signaling induces the GSK3β-mediated proteasomal degradation of the tumor suppressor CCDC6. We show that CCDC6 acts as a negative regulator of the xCT-promoting transcription factor ATF4 by directly interacting with it and preventing its recruitment to the xCT promoter. Consequently, KRAS-driven CCDC6 degradation disinhibits ATF4, leading to increased xCT expression, elevated intracellular GSH, and enhanced resistance to ferroptosis.

Crucially, pharmacological inhibition of CCDC6 turnover using proteasome, GSK3β, or specific KRAS mutant inhibitors (Sotorasib, Adagrasib, HRS4642) restored CCDC6 protein levels and robustly sensitized KRAS-mutated cells to ferroptosis-inducing agents like Sulfasalazine. Furthermore, validation in preclinical models and human colorectal cancer samples revealed that CCDC6 protein levels are predominantly downregulated in KRAS-mutant cases

This work uncovers a novel KRAS/CCDC6/xCT signaling axis that mediates ferroptosis resistance in KRAS-mutated cancers. Moreover, it identifies CCDC6 turnover as a critical vulnerability and a promising therapeutic target to enhance the efficacy of ferroptosis-inducing agents.

## INTRODUCTION

RAS oncogenes are among the most frequently mutated genes in human cancers, particularly in lung adenocarcinoma (LUAD), colorectal carcinoma (CRC), and pancreatic ductal adenocarcinoma (PDAC) [1, 2]. The RAS gene family includes the KRAS, NRAS, and HRAS members, whose genes are located on different chromosomes

Among the three isoforms, KRAS mutations are the most prevalent isoform in human cancers. They occur in nearly 90% of Pancreatic Ductal Adenocarcinoma (PDAC) cases -primarily at codon 12- as well as 20–50% of Lung Adenocarcinoma (LUAD), where they are linked to poor prognosis. Additionally, KRAS mutations are found in over 40% of Colorectal Cancer (CRC) cases, typically involving codons 12, 13, and 61. In contrast, other RAS isoforms are less frequently mutated: NRAS mutations are mainly detected in melanoma and thyroid cancer (approximately 5%). HRAS mutations are the rarest in human tumors, appearing primarily in bladder and cervical cancers at a rate of roughly 2% [3].

These mutations constitutively lock the RAS protein in its "ON" (GTP-bound) state, leading to continuous activation of downstream pro-proliferative pathways like RAF-MEK-ERK and anti-apoptotic pathways such as PI3K-AKT. This sustained signaling collectively drives neoplastic transformation and tumor growth [4, 5]. Significantly, RAS mutations markedly limit the efficacy of anti-EGFR monoclonal antibodies (e.g., cetuximab, panitumumab) in cancers like CRC, as these treatments are primarily effective only in patients with wild-type RAS. This presents a considerable challenge in cancer treatment [6, 7, 8].

Beyond their direct proliferative effects, RAS oncogenes also exert their transforming power by influencing the cell’s redox balance, thereby favoring tumor progression [9]. A key discovery is that the KRAS oncogene helps cancer cells cope with oxidative stress by boosting intracellular levels of glutathione (GSH), one of the most powerful antioxidants. This is achieved through increased production of xCT/SLC7A11, a cystine-glutamate antiporter crucial for importing cystine, a key building block for GSH synthesis [9].

The KRAS oncogene upregulates xCT by activating the Ras-Raf-Mek-Erk pathway, which in turn leads to the synergistic activation of transcription factors ETS-1 and ATF4, ultimately "switching on" the xCT gene [10]. Essentially, KRAS doesn’t just enable cells to grow uncontrollably; it also equips them with a robust defense mechanism against oxidative stress. This increased xCT expression is not merely a reactive measure, but an inherent strategy employed by the KRAS oncogene to actively support and sustain cellular transformation and tumorigenesis. The xCT channel’s role in promoting tumorigenesis through its antioxidant activity has been observed across various cancer types, highlighting its significance in cancer development [9, 10].

xCT is also central to ferroptosis, an iron-dependent form of regulated cell death characterized by lipid peroxides accumulation, distinct from other cell death mechanisms. Ferroptosis is gaining interest as a tumor suppression mechanism, with some cancer therapies even inducing it through xCT modulation [9]. Both apoptosis and ferroptosis act as tumor suppressors by eliminating cells under oxidative stress. In tumor cells, xCT inactivation or cystine removal induces ferroptosis, while xCT overexpression promotes GSH synthesis and confers ferroptosis resistance. Furthermore, the xCT channel plays a crucial role in early carcinogenesis by being negatively regulated by tumor suppressors like p53 and BAP1 [10].

Our previous work demonstrated that also the loss of the CCDC6 tumor suppressor enhances xCT transcription, leading to oxidative stress tolerance and ferroptosis resistance in cell cultures in vitro [11, 12]. CCDC6 (Coiled Coil Domain Containing 6) is a ubiquitous protein with both nuclear and cytosolic localization, endowed with pro-apoptotic activity and involved in DNA damage repair mechanisms. It is also known to negatively regulate CREB family members [13]. Given that the regulation of the xCT promoter significantly depends on the CREB family member ATF4, which acts synergistically with ETS1, we investigated whether the loss or functional alteration of CCDC6, particularly in the context of oncogenic RAS signaling, may contribute to uncontrolled ATF4 activity. This, in turn, could determine augmented xCT transcription, increased GSH levels, and evasion from regulated cell death pathway of ferroptosis. These data are corroborated by the observation that low or absent CCDC6 protein levels significantly correlate with KRAS mutation in CRC samples.

Here, we propose a novel mechanism with which KRAS-mutated cells increase their tolerance to oxidative stress and prevent the activation of ferroptosis: the loss of CCDC6 activity due to KRAS-mediated protein degradation.

Taken together, our results confirm a direct link between RAS-driven CCDC6 turnover, xCT upregulation, and ferroptosis evasion. In conclusion, we unveil a novel mechanism of tumor suppression and also open new therapeutic avenues for targeting this critical pathway in RAS-mutated cancers. Moreover, our data suggest that the loss of CCDC6 by alternative, still uncharacterized, molecular mechanisms, may represent a marker of ferroptosis resistant tumors.

## RESULTS

### Human cancer cells carrying KRAS mutations exhibit low CCDC6 protein levels, increased xCT expression and ferroptosis resistance

In human non-small cell lung cancer (NSCLC) cell lines carrying KRAS mutations [H460Q61H and HOP62G12C], we observed lower CCDC6 protein levels compared to KRAS-wild type NSCLC cells [H1975WT], despite no corresponding changes in transcript levels (Figure 1A, B). Notably, constitutively active RAS mutants in different tumors increase GSK3β kinase levels [14], and we have shown that GSK3β, by phosphorylating the CCDC6 degron, leads to CCDC6 recognition by the FBXW7 E3 ubiquitin ligase and proteasome-dependent degradation [15]. Thus, our observations indicated an increased protein turnover of CCDC6 in the KRAS mutated cells, driven by post-transcriptional mechanisms.

**Figure 1:**
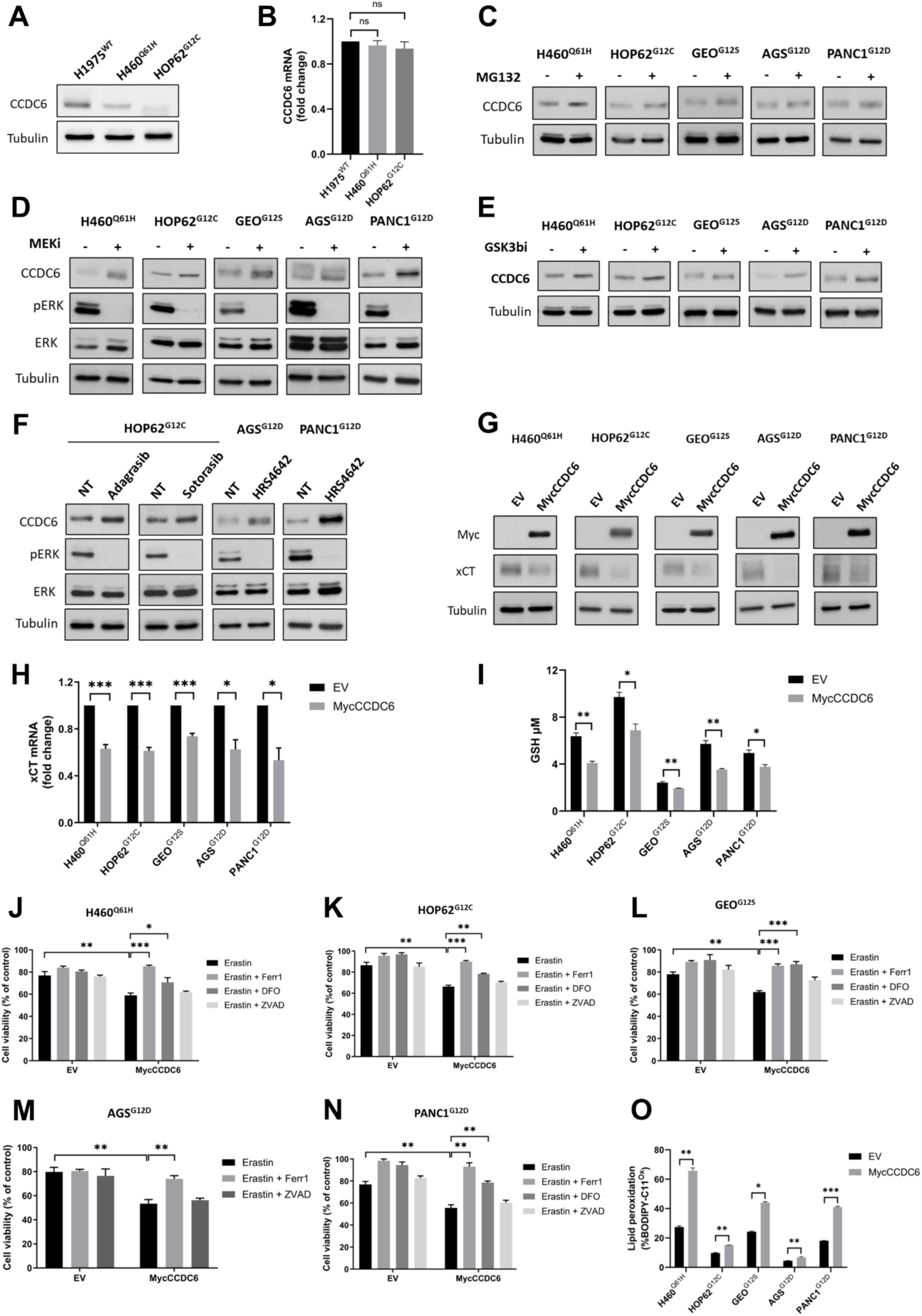
KRAS mutations drive CCDC6 degradation, xCT up-regulation and ferroptosis resistance. (A) Western blot analysis of CCDC6 expression in H1975, H460 and HOP62 cells. Anti-tubulin is shown as loading control. (B) Relative CCDC6 expression assessed by qPCR in H1975, H460 and HOP62 cells. (C-E) Western blot analysis of CCDC6 expression in H460, HOP62, GEO, AGS and PANC1 cells treated with MG132 [25μM] for 1h (C), MEK inhibitor Selumetinib [10μM] for 2h (D), or GSK3ϕ3 inhibitor SB216763 [10μM] for 2h (E), as indicated. Anti-pERK and anti-ERK immunoblots demonstrate the efficacy of the pharmacological treatments. Anti-tubulin immunoblots are shown as loading control. (F) Western blot analysis of CCDC6 expression in HOP62 cells treated with the KRASG12C inhibitors Adagrasib [6μM] for 3h or Sotorasib [6μM] for 1h, and in AGS and PANC1 cells treated with the KRASG12D inhibitor HRS4642 [3nM] for 30min and [6μM] for 1h, respectively. Anti-pERK and anti-ERK immunoblots demonstrate the efficacy of the pharmacological treatments. Anti-tubulin immunoblots are shown as loading control. (G) Western blot analysis of xCT expression in H460, HOP62, GEO, AGS and PANC1 cells upon transient transfection of MycCCDC6 or empty vector (EV), as a control. Anti-myc and ant-tubulin immunoblots are shown as transfection and loading control, respectively (H) Relative xCT expression assessed by qPCR in the same cells population as in (G). (I) Intracellular GSH levels in H460, HOP62, GEO, AGS and PANC1 cells upon transient transfection of MycCCDC6 or empty vector (EV), as a control. (J-N) Cell viability was assessed in H460 (J), HOP62 (K), GEO (L), AGS (M) and PANC1 (N) cells transfected with MycCCDC6 or empty vector (EV) and treated with Erastin (20μM H460, 2.5μM HOP62, 4μM GEO, 20μM AGS, 4μM PANC1) alone or combined with 10μM Z-VAD-fmk (ZVAD), 10μM ferrostatin-1 (Ferr1) or 100μM deferoxamine (DFO) for 24h. (O) Bar Graphs show the percentage of oxidized BODIPY-C11 positive cells in indicated cells upon transient transfection of MycCCDC6 or empty vector (EV), as control. Where shown, data are reported as mean ± SEM of 3 independent repeats. Statistical significance was verified by 2-tailed Student’s t-test (* p <0.05; ** p <0.01, *** p <0.001)

We then asked if the increased CCDC6 turnover could be driven by different KRAS mutations, and assessed which KRAS downstream pathways was eventually required. To this aim, we measured CCDC6 protein levels in response to proteasome inhibitor (MG132), MEK1/2 inhibitor (Selumetinib), or GSK3β inhibitor (SB216763) in a panel of KRAS-mutated human cancer cell lines from NSCLC [H460Q61H and HOP62G12C], colon cancer [GEOG12S], gastric adenocarcinoma [AGSG12D] and pancreatic cancer [PANC-1G12D]. The restoration of CCDC6 levels upon treatment with each inhibitor suggested that KRAS signaling activation influenced the increased CCDC6 turnover (Figure 1C, D, E)

Moreover, the silencing of the GSK3b kinase determined an appreciable increase of the CCDC6 levels (Figure S1A). Noteworthy, specific KRAS inhibitors targeting the KRASG12C (Sotorasib, Adagrasib) and KRASG12D (HRS4642) variants also rescued CCDC6 protein levels in the HOP62^G12C^, in AGS^G12D^ and in PANC-1^G12D^ cells (Figure 1F), confirming that the oncogenic activity of mutated KRAS leads to an increased turnover of CCDC6.

The upregulation of cystine/glutamate antiporter xCT has been implicated in supporting KRAS oncogenic activity by maintaining intracellular redox balance [10]. Recently, we demonstrated that the loss of CCDC6 function is associated with increased xCT expression and ferroptosis resistance [11].

Consistent with this observation, re-expressing myc-tagged CCDC6 in KRAS-mutated cancer cells resulted in a decrease in xCT protein and mRNA levels (Figure 1G, H), accompanied by a reduction in xCT activity, as evidenced by a decrease in intracellular glutathione levels (Figure 1I) and in cystine uptake (Figure S1B). These data suggest that RAS-induced CCDC6 proteasome-dependent degradation may contribute to an antioxidant program that protects KRAS mutated cancer cells from ferroptosis via xCT upregulation.

To test this hypothesis, we treated the KRAS-mutated cells with the ferroptosis inducer Erastin, in the presence or absence of ferroptosis or apoptosis inhibitors and evaluated cell viability. While Erastin treatment exerted limited effects on cell viability, transient re-expression of CCDC6 significantly potentiated Erastin effects reducing the survival of these cells, and this effect was significantly reversed by Ferrostatin and Deferoxamine, and to a lesser extent by Z-VAD (Figure 1J-N).

Accordingly, restoring CCDC6 expression increased lipid peroxidation, a key marker of ferroptosis, as detected by flow cytometry using the C11-BODIPY fluorescent probe (Figure 1O).

### Ectopic expression of different KRAS oncogenic variants in KRAS wild-type cells Reproduces the ferroptosis-resistant phenotype of KRAS-Mutated Cancer Cells

To validate and further confirm that various RAS mutations affect CCDC6 levels, we transiently overexpressed different KRAS variants in the KRAS wild type 293T cells.

Consistent with our primary findings in human cancer cells harboring endogenous KRAS mutations, transient expression of different KRAS mutants reduced CCDC6 protein levels (Figure 2A) without altering transcript levels (Figure 2B).

**Figure 2:**
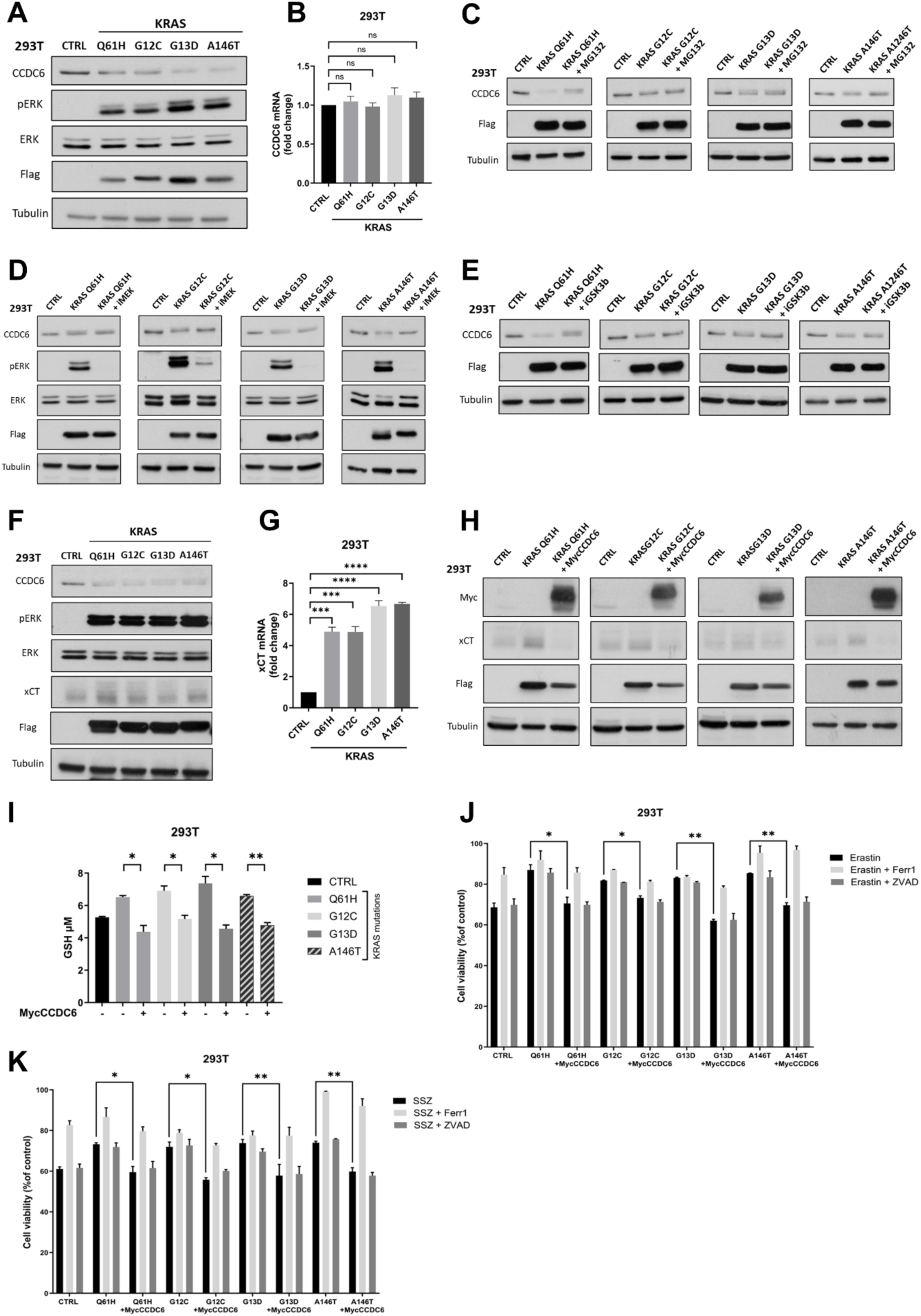
Ectopic expression of KRAS variants triggers CCDC6 degradation and ferroptosis resistance in 293T cells. (A) Western blot analysis of CCDC6 expression in 293T cells transfected with mutated KRAS isoforms (Q61H, G12C, G13D, A146T) or empty vector (CTRL), as control. Anti-pERK, anti-ERK, and anti-flag immunoblots were performed to confirm the activity and expression of Flag-KRAS mutants, respectively. Anti-tubulin is shown as loading control. (B) Relative CCDC6 expression assessed by qPCR in the same cell populations as in (A). (C-E) Western blot analysis of CCDC6 in 293T cells expressing mutated KRAS isoforms (Q61H, G12C, G13D, A146T) or empty vector (CTRL) after treatment with MG132 [20μM] for 30min (C), MEK inhibitor Selumetinib [10μM] for 3h (D) or GSK3ϕ3 inhibitor SB216763 [10μM] for 3h (E), as indicated. Anti-pERK and anti-ERK immunoblots demonstrate the efficacy of the pharmacological treatment. Anti-flag immunoblots confirm Flag-KRAS mutant expression. Anti-tubulin is shown as loading control. (F) Western blot analysis of xCT expression in the same cell populations as in (A). Anti-pERK, anti-ERK, and anti-flag immunoblots were performed to confirm the activity and expression of Flag-KRAS mutants. Anti-tubulin is shown as loading control. (G) xCT relative expression assessed by qPCR in the same cell populations as in (A). (H) Western blot analysis of xCT expression in 293T cells transfected with mutated KRAS isoforms (Q61H, G12C, G13D, A146T) or empty vector (CTRL), in presence or absence of MycCCDC6. Anti-myc and anti-flag immunoblots were performed to confirm the expression of MycCCDC6 and Flag-KRAS mutant plasmids, respectively. Anti-tubulin is shown as loading control. (I) Intracellular GSH levels in the 293T cells transfected with mutated KRAS isoforms (Q61H, G12C, G13D, A146T) or empty vector (CTRL), in presence or absence of MycCCDC6 (J) Cell viability was assessed in indicated cells upon treatment with Erastin [2μM] alone or combined with 10μM Z-VAD-fmk (ZVAD) or 10μM ferrostatin-1 (Ferr1), for 24h. (K) Cell viability was assessed in indicated cells upon treatment with Sulfasalazine (SSZ) [500μM] alone or combined with 10μM Z-VAD-fmk (ZVAD) or 10μM ferrostatin-1 (Ferr1), for 24h. Where shown, data are reported as mean ± SEM of 3 independent repeats. Statistical significance was verified by 2-tailed Student’s t-test (* p <0.05; ** p <0.01, *** p <0.001 and **** p <0.0001).

The CCDC6 degradation was reversed by inhibiting the proteasome (MG132) or interfering with the MEK1/2 and GSK3β signaling pathways (Figure 2C, D, E), mirroring our observations in the KRAS-mutated cancer cell lines. Like our findings in neoplastic cells [15], silencing of the FBXW7-ubiquitin ligase also increased CCDC6 levels (Figure S2).

The results from the KRAS wildtype 293T cells, when overexpressing the KRAS mutated variants, strongly reproduce the key evidence that low CCDC6 protein levels are associated with high levels of the cystine/glutamate antiporter xCT, evaluated both as transcript and as protein by Real Time PCR and Western Blot, respectively (Figure 2F, G).

Moreover, re-expressing CCDC6 in these cells reduces xCT channel expression and GSH intracellular levels (Figure 2H, I), further supporting the regulatory role of CCDC6 on xCT expression. Crucially, the restoration of CCDC6 levels in KRAS mutant-overexpressing cells re-sensitized the cells to ferroptotic agent Erastin and Sulfasalazine (Figure 2J, K, C), confirming the essential contribution of CCDC6 to RAS-induced ferroptosis resistance, as seen in our cancer models.

### CCDC6 Negatively Regulates ATF4-Mediated xCT Expression

CCDC6 is a known interactor and negative regulator of several CREB family members [11, 16–18]. ATF4 is a critical mediator of the integrated stress response (ISR) and is activated by various stress signals [19]. Genes regulated by ATF4 are primarily involved in amino acid synthesis, metastasis, angiogenesis, and drug resistance. Elevated ATF4 expression is observed in several neoplastic diseases. Notably, in tumorigenesis processes dependent on ATF4 activity, a transcriptional increase of the xCT channel is reported, which is associated with resistance to ferroptosis and cytotoxic stress [20]. Furthermore, oncogenic forms of KRAS have been shown to protect fibroblasts from oxidative stress by increasing intracellular GSH levels [9, 10]. At the molecular level, this glutathione increase directly depends on the transcriptional upregulation of the cystine/glutamate antiporter gene, xCT. This regulation, dependent on the RAS oncogenic pathway, is mediated by the ETS-1 transcription factor, which synergizes with ATF4 to directly transactivate the xCT promoter [10].

Through co-immunoprecipitation experiments, we confirmed the interaction between CCDC6 and ATF4 (Figure 3A). We then investigated whether CCDC6 could modulate the ATF4 transcriptional activity on xCT promoter in response to oncogenic RAS signaling.

**Figure 3:**
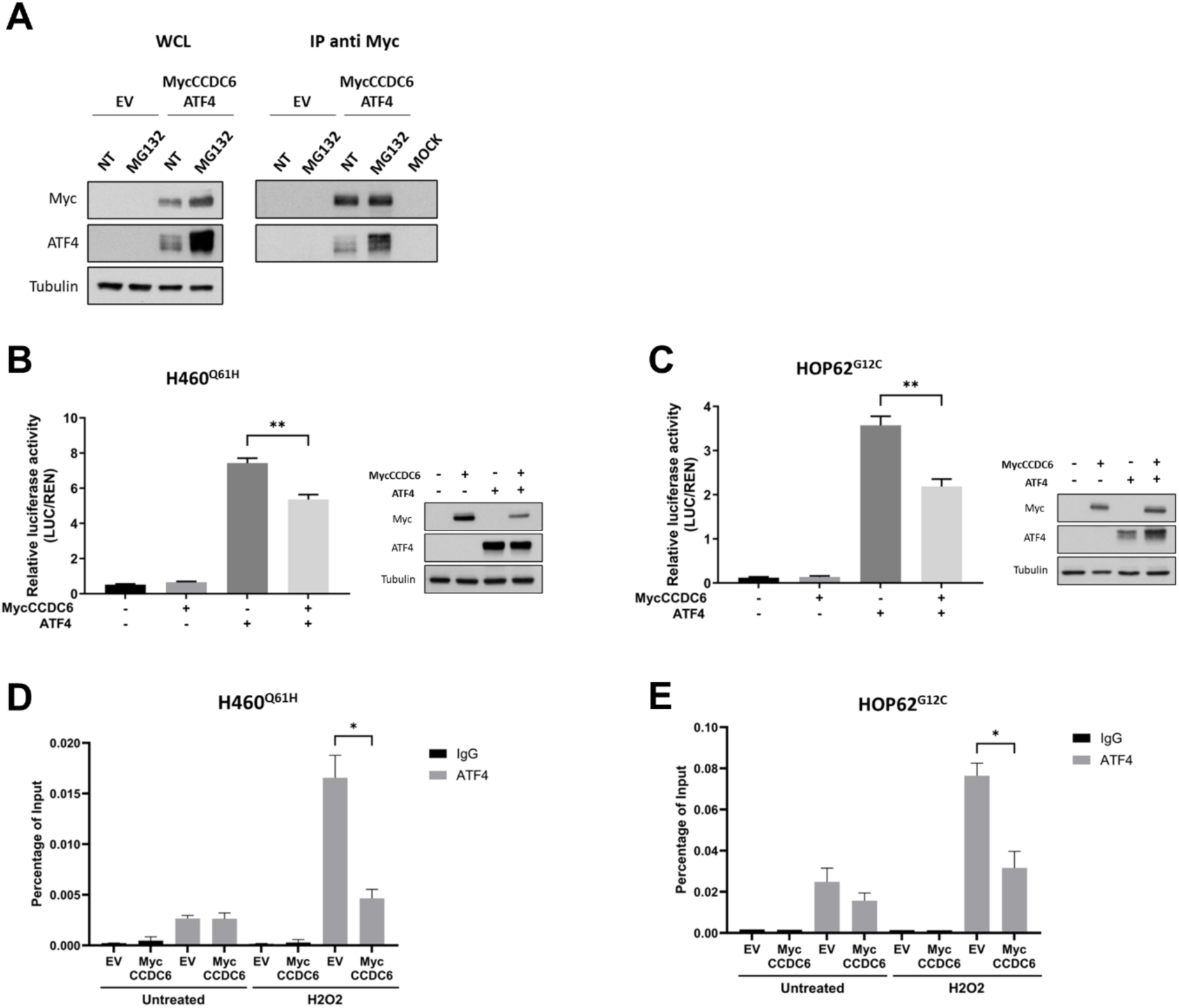
CCDC6 interacts with ATF4 and negatively regulates its transcriptional activity on the xCT promoter. (A) 293T cells were co-transfected with MycCCDC6 and ATF4 plasmids, or empty vector (EV) for 48h and then treated with the proteasome inhibitor MG132 [20 µM] for 4h. Protein extracts were subjected to immunoprecipitation (IP) with anti-myc antibody, or with immunoglobulin G (mock) as a control, followed by immunoblotting with anti-myc and anti-ATF4 antibodies. Total whole cell lysates (WCL), shown on the left, were immunoblotted with the indicated antibodies. (B-C) Luciferase reporter assay and Western blot analysis in H460 (B) and HOP62 (C) cell lines. Cells were co-transfected with a firefly luciferase construct containing the xCT promoter and a Renilla luciferase expression construct, in the presence or absence of MycCCDC6 and/or ATF4 plasmids. Firefly and Renilla luciferase activities were determined using the Dual-Luciferase reporter system (Promega). For each sample, the Firefly/Renilla ratio (LUC/REN) is shown as Relative luciferase activity. Transfection efficiency of MycCCDC6 and ATF4 plasmids was evaluated by Western blot analysis using anti-myc and anti-ATF4 antibodies. Anti-tubulin was used as a loading control. (D-E) ChiP assay was carried out on H460 (D) and HOP62 (E) cells transfected with MycCCDC6 or empty vector (EV), as control, and treated with of H2O2 to induce oxidative stress. qTR-PCR was used to determine the percentage enrichment of ATF4 binding on xCT promoter over input. Immunoglobulin G (IgG) was used as a negative control. Where shown, data are reported as mean ± SEM of 3 independent repeats. Statistical significance was verified by 2-tailed Student’s t-test (* p <0.05; ** p <0.01).

To this end, we performed a luciferase assay in KRAS mutant cancer cell lines [H460^Q61H^ and HOP62^G12C^] which exhibit increased CCDC6 turnover due to RAS oncogenic activity.

We transiently re-expressed CCDC6, both with and without a vector overexpressing ATF4, in the presence of a xCT-promoter luciferase reporter plasmid.

Interestingly, overexpression of CCDC6 significantly reduced the luciferase activity observed in the presence of ATF4 (Figure 3B, C), suggesting that CCDC6 negatively regulates the transcriptional activity of ATF4 on the xCT promoter. To investigate whether CCDC6 directly influences the binding of ATF4 to the xCT promoter, we performed chromatin immunoprecipitation (ChIP) assays in H460^Q61H^ and HOP62^G12C^ cell lines under basal conditions and following treatment with H₂O₂, in the presence or in the absence of CCDC6 enforced expression. Consistent with previous reports [10], oxidative stress strongly enhanced ATF4 recruitment to the xCT (Figure 3D, E). However, ATF4 recruitment to the xCT promoter was significantly impaired in CCDC6-overexpressing KRAS mutant cells (mycCCDC6) compared to control cells (EV) (Figure 3D, E), indicating that CCDC6 prevents the physical association of ATF4 with the xCT promoter.

Collectively, these results indicate that increased CCDC6 turnover, resulting from oncogenic RAS activity, allows the ATF4 recruitment to the xCT promoter, thereby promoting xCT expression and increasing resistance to oxidative stress.

### Inhibition of CCDC6 protein degradation sensitizes KRAS mutated cancer cells to ferroptosis inducers

Finally, we aimed to evaluate whether pharmacological inhibition of CCDC6’s ubiquitin-dependent degradation could increase the sensitivity of KRAS mutant cancer cells [H460^Q61H^ and HOP62^G12C^, AGS^G12D^, PANC1^G12D^] to ferroptosis-inducing agents. To this end, we treated cells with Bortezomib, the first proteasome inhibitor used clinically [21] and with the GSK3b inhibitor SB 216763 [22, 23], which were able to increase CCDC6 protein levels in KRAS-mutated cancer cells (Figure 1C - E; 2C- E). These agents were used alone or in combination with Sulfasalazine (SSZ), an anti-inflammatory agent also known for its ability to inhibit xCT and consequently induce ferroptosis by depleting intracellular glutathione [24].

Treatment with Sulfasalazine (SSZ) as single agent showed a limited effect on cell viability, while a significant reduction in cell viability was observed when Sulfasalazine (SSZ) was combined with Bortezomib or GSK3β inhibitor (Figure 4A - E). To prove that CCDC6 protein stability is involved in modulating ferroptosis sensitivity, we used P5091, an inhibitor of USP7, the deubiquitinase of CCDC6, which increases CCDC6 degradation. As shown in figures 4A – E, P5091 treatment reverted the effect of Sulfasalazine/GSK3β inhibitor combination.

**Figure 4:**
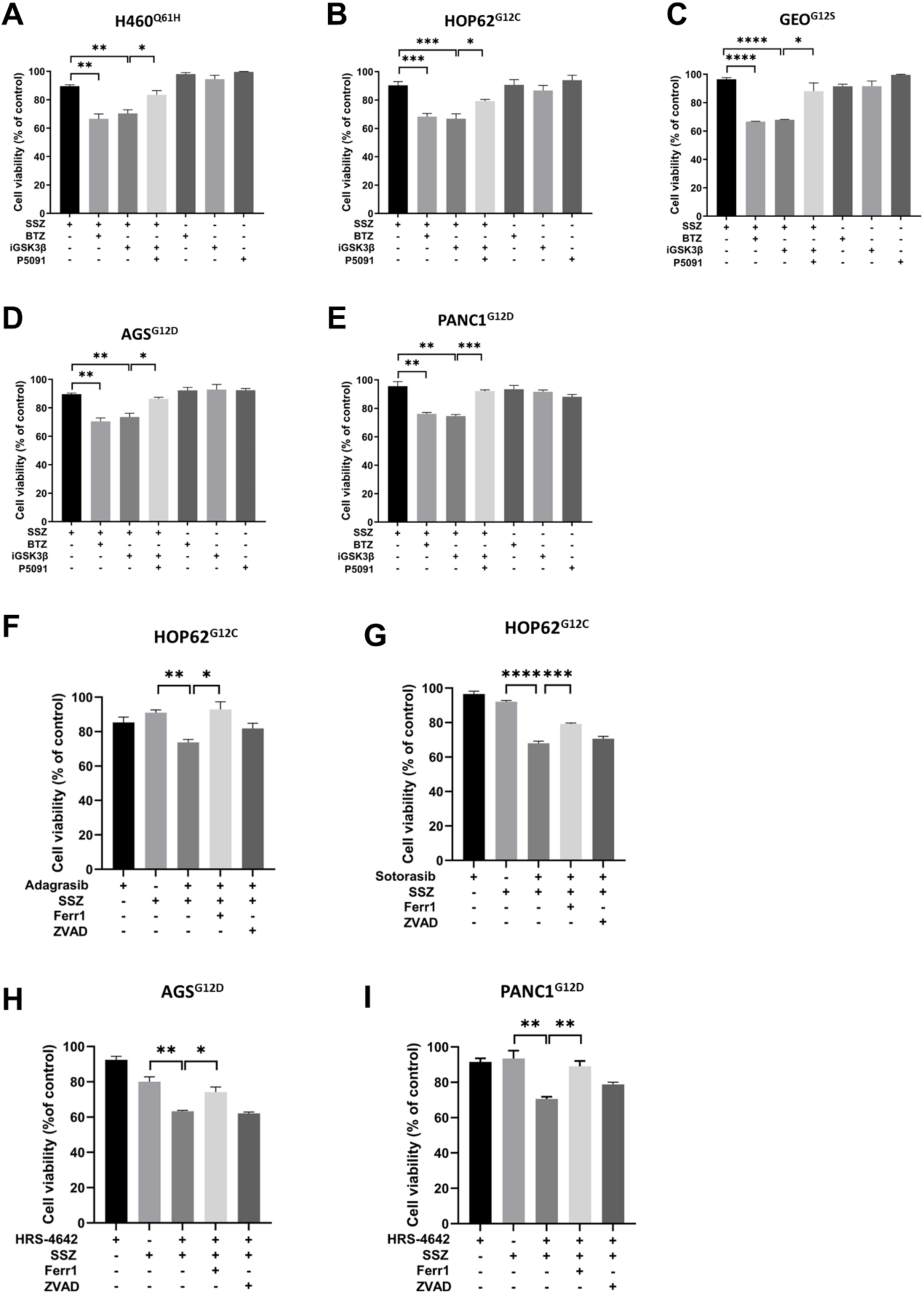
Inhibition of CCDC6 protein degradation enhances sensitivity to ferroptosis inducers. (A-E) Cell viability was assessed in H460 (A), HOP62 (B), GEO (C), AGS (D) and PANC1 (E) cells treated with the ferroptosis inducers Sulfasalazine (SSZ), the proteasome inhibitor Bortezomib (BTZ), the GSK3β inhibitor SB216763 (iGSK3β), or the USP7 deubiquitinase inhibitor (P5091), either alone or in combination. The drug concentrations were as follows: SSZ at 800µM (H460), 200µM (HOP62, PANC1, GEO), 1 mM (AGS); BTZ at 5 nM (H460, AGS, PANC1), 10 nM (HOP62), 4nM (GEO); iGSK3β at 10 µM (H460), 40 µM (HOP62), and 20 µM (AGS, PANC1, GEO); P5091 at 2.5 µM (H460), 1µM (HOP62, AGS, PANC1, GEO) (F-G) HOP62 cell viability was measured after treatment with Sulfasalazine (SSZ) [400µM], used alone or combined with the KRASG12C inhibitors Adagrasib [2µM] (F) or Sotorasib [40µM] (G). The ferroptosis inhibitor ferrostatin-1 (Ferr1) [10 µM] or the pan-caspase inhibitor Z-VAD-fmk (ZVAD) [10 µM] were added to the SSZ-containing combinations. (H-I) Cell viability was assessed in AGS (H) and PANC1 (I) treated with Sulfasalazine (SSZ), [1mM] in AGS and [200µM] in PANC1, used alone or combined with the KRASG12D inhibitor HRS4642, [6nM] in AGS and [5µM] PANC1. The ferroptosis inhibitor ferrostatin-1 (Ferr1) [10 µM] or the pan-caspase inhibitor Z-VAD-fmk (ZVAD) [10 µM] were added to the SSZ-containing combinations. Where shown, data are reported as mean ± SEM of 3 independent repeats. Statistical significance was verified by 2-tailed Student’s t-test (* p <0.05; ** p <0.01, *** p <0.001 and **** p <0.0001).

The specific KRASG12C inhibitors, Sotorasib and Adagrasib, are currently under investigation in several clinical trials for advanced or metastatic NSCLC [25]. Recently, a novel KRASG12D inhibitor, HRS4642, demonstrating potent and selective anti-tumor activity across various models, has entered a Phase III clinical trial (NCT07438106) for patients with locally advanced pancreatic cancer harboring the KRASG12D mutation [26].

As detailed in Figure 1F, the use of the specific KRAS inhibitors Sotorasib, Adagrasib, or HRS4642 determined the restoration of CCDC6 protein levels in HOP62^G12C^, AGS^G12D^, PANC1^G12D^ cells, indicating that the increased CCDC6 turnover was dependent on oncogenic KRAS signaling.

To evaluate whether targeting specific KRAS mutations could sensitize cancer cells to ferroptosis, we performed cell viability assays using specific KRAS inhibitors (Sotorasib or Adagrasib for HOP62^G12C^ cells, and HRS4642 for AGS^G12D^ and PANC1^G12D^ cells) in combination with the ferroptosis-inducing agent Sulfasalazine (SSZ). Our results show that a significant response was observed when Sulfasalazine (SSZ) was used in combination with the KRAS-specific inhibitors, while single-agent treatments with either the specific KRAS inhibitors or Sulfasalazine (SSZ) had a limited effect on cell viability (Figure 4F-I). This reduction in viability was considerably restored by the ferroptosis inhibitor ferrostatin, and to a lesser extent by the apoptosis inhibitor Z-VAD-fmk (Figure 4F-I), indicating that the cell death induced by the combination of a specific KRAS inhibitor and Sulfasalazine (SSZ) is predominantly mediated by ferroptosis.

To quantify the observed synergy, we analyzed the dose-response curves for Sulfasalazine (SSZ) in the presence and absence of a fixed concentration of the specific KRAS inhibitors.

The results show that the combined treatment reduced the IC50 of Sulfasalazine (SSZ) in all tested cell lines (Figure S3A-D). The synergistic nature of the interaction was confirmed by the Combination Index (CI) values, which were less than 1.0 in all tested combination (Figure S3A-D). Additionally, Dose Reduction Index (DRI) values for Sulfasalazine (SSZ) were greater than 1.0 (Figure S3A-D), indicating that a lower dose of Sulfasalazine (SSZ) is needed to achieve the same effect when used in combination, further confirming the synergy.

Overall, our results indicate that targeting the KRAS-driven degradation of CCDC6, either indirectly with proteasome or GSK3β inhibitors, or directly, with specific KRAS inhibitors, represents a promising therapeutic approach to increase the vulnerability of KRAS-mutated cancer cells to ferroptotic cell death.

### CCDC6 protein levels correlate with KRAS mutations in PDAC preclinical models and in human colorectal carcinoma (CRC) primary samples

CCDC6 expression and modulation was investigated in PDAC preclinical models. Immunohistochemical (IHC) analysis of xenografts tissue samples, derived from orthotopic FC1199 (KRASG12D) cell transplants in the pancreas [27], showed low CCDC6 staining intensity compared to adjacent normal pancreatic tissue, as shown by the tumor/normal DAB OD ratio (Figure 5A).

**Figure 5:**
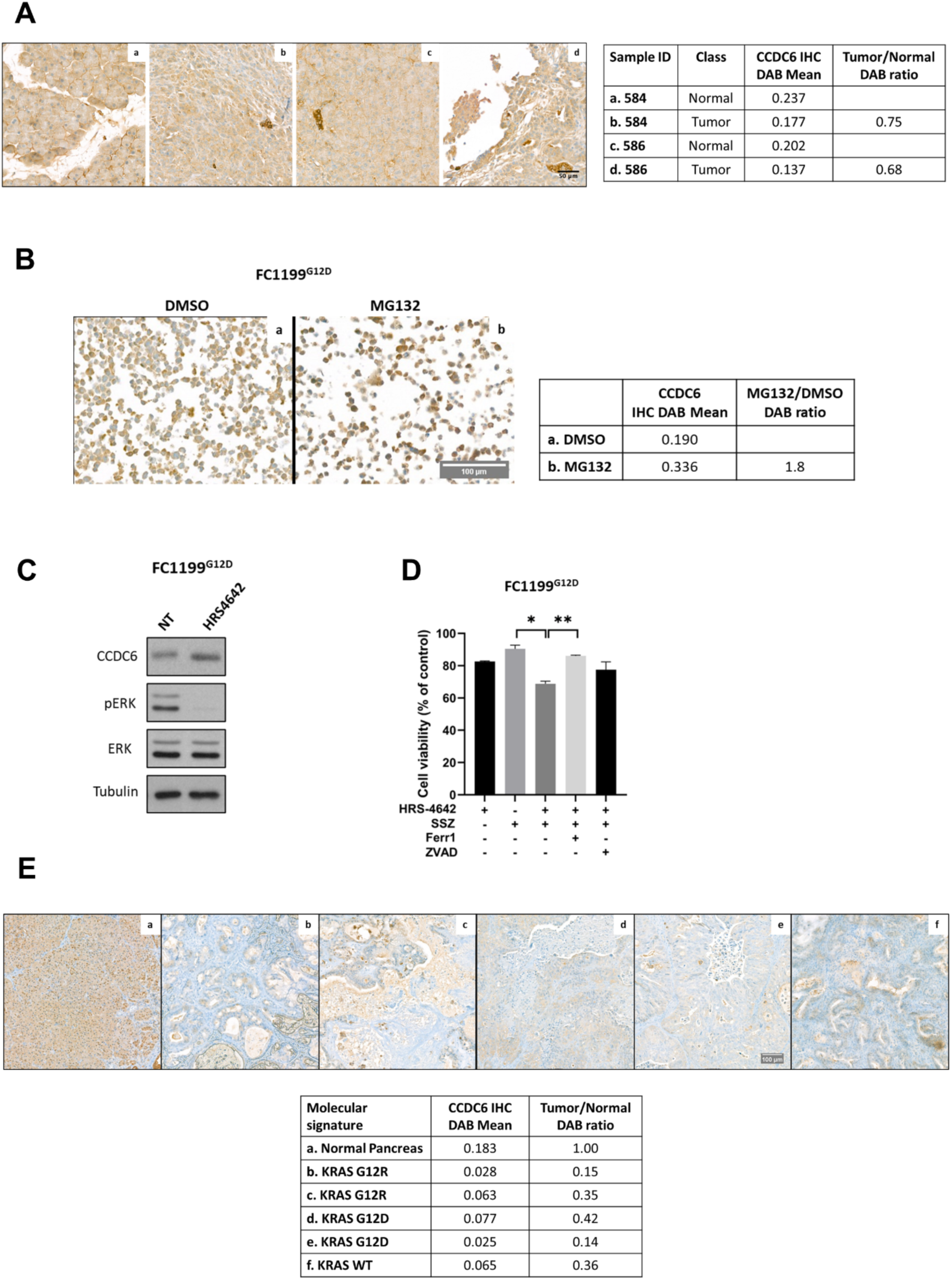
CCDC6 protein levels are regulated by KRAS signaling in PDAC preclinical models. (A) Immunohistochemical (IHC) analysis of CCDC6 expression in xenografts tissue sample derived from orthotopic FC1199 cell transplants in the pancreas. Representative images showing CCDC6 protein expression levels in pancreatic normal and tumor sections from two independent samples (#584 and #586). Protein levels were evaluated in tumor sections and compared with adjacent normal pancreatic tissue. Scale bar = 50 µm. The table (right) displays the quantitative analysis of the IHC signal expressed as DAB mean intensity and Tumor/Normal DAB ratio. (B) Immunocytochemical (IHC) analysis of CCDC6 protein expression levels in murine FC1199 cells. Representative images showing CCDC6 protein levels in FC1199 cells treated with vehicle DMSO (a) or the proteasome inhibitor MG132 (b) [10µM] for 8h. Scale bar = 10 µm. The table (right) displays the quantitative analysis of the IHC signal expressed as DAB mean intensity and MG132/DMSO DAB ratio. (C) Western blot analysis of CCDC6 expression in murine FC1199 pancreatic cancer cells treated with the KRASG12D inhibitor HRS4642 [5µM] for 3h. Anti-pERK and anti-ERK immunoblots demonstrate the efficacy of the pharmacological treatment. Anti-tubulin immunoblot is shown as loading control. (D) Cell viability was assessed in FC1199 cells treated with sulfasalazine (SSZ) [400µM] used alone or combined with the KRASG12D inhibitor HRS4642 [1µM] for 1.25h. The ferroptosis inhibitor ferrostatin-1 (Ferr1) [10 µM] or the pan-caspase inhibitor Z-VAD-fmk (ZVAD) [10 µM] were added to the SSZ-containing combinations. (E) Immunohistochemical (IHC) analysis of CCDC6 expression in Patient-Derived Xenografts (PDX) of PDAC. Representative images (top) showing CCDC6 protein levels in normal pancreatic tissue (a) and in PDX samples carrying KRAS mutations KRAS G12R (b, c) or KRAS G12D (d, e), and KRAS wild-type (WT) (f). Scale bar = 100 µm. The table (bottom) displays the quantitative analysis of the IHC signal expressed as DAB mean intensity and Tumor/Normal DAB ratio. Where shown, data are reported as mean ± SEM of 3 independent repeats. Statistical significance was verified by 2-tailed Student’s t-test (* p <0.05; ** p <0.01).

Accordingly, immunocytochemical analysis of cell block preparations from FC1199 cells revealed a notable increase in CCDC6 protein levels following treatment with the proteasome inhibitor MG132, as evidenced by the MG132/DMSO DAB OD ratio (Figure 5B). Furthermore, in these same cells, treatment with the specific KRASG12D inhibitor, HRS4642, increased CCDC6 protein levels (Figure 5C) and significantly sensitized the cells to Sulfasalazine (SSZ) (Figure 5D), as observed by Western blot analysis and cell viability assays.

Consistently, tissue samples from patient-derived xenografts (PDX) of PDAC cases carrying KRASG12D or KRASG12R mutations (N = 4) [28] exhibited lower CCDC6 expression relative to normal pancreatic ductal tissue, as quantified by tumor/normal DAB OD ratio (Figure 5E, a, b, c, d, e). Notably, one KRAS wild-type PDX sample also exhibited low CCDC6 expression, yet it showed positive anti-phospho-MAPK staining, supporting the activation of downstream RAS signaling pathways (Figure 5E, f; Figure S4, b), In a parallel cohort of 101 primary colorectal carcinomas, CCDC6 protein expression was evaluated by two independent pathologists using a semi-quantitative scoring system that integrated staining intensity (1+, 2+, 3+) and the percentage of positive tumor cells within neoplastic areas. Tumors were stratified according to RAS mutational status into MUT (n = 60) and WT (n = 41) groups (Figure 6A). Aside from three samples in which no staining for CCDC6 was observed, protein expression differences were captured using a composite semi-quantitative H-score, calculated from pathologist-assigned percentages of 1+, 2+, and 3+ tumor cells [score = (1×%1+ + 2×%2+ + 3×%3+)/100] (Figure 6B). Representative images are shown in Figure S5. Using this integrated metric, the KRAS/NRAS mutated tumors (MUT) exhibited lower CCDC6 expression compared to KRAS wild type tumors (WT) (mean 1.50 vs 1.87: median 1.48 vs 2.00). This difference reached statistical significance (two-sided Mann–Whitney U test, p = 0.034), indicating a shift toward weaker staining intensity in the mutant cohort (Table 1).

**Figure 6:**
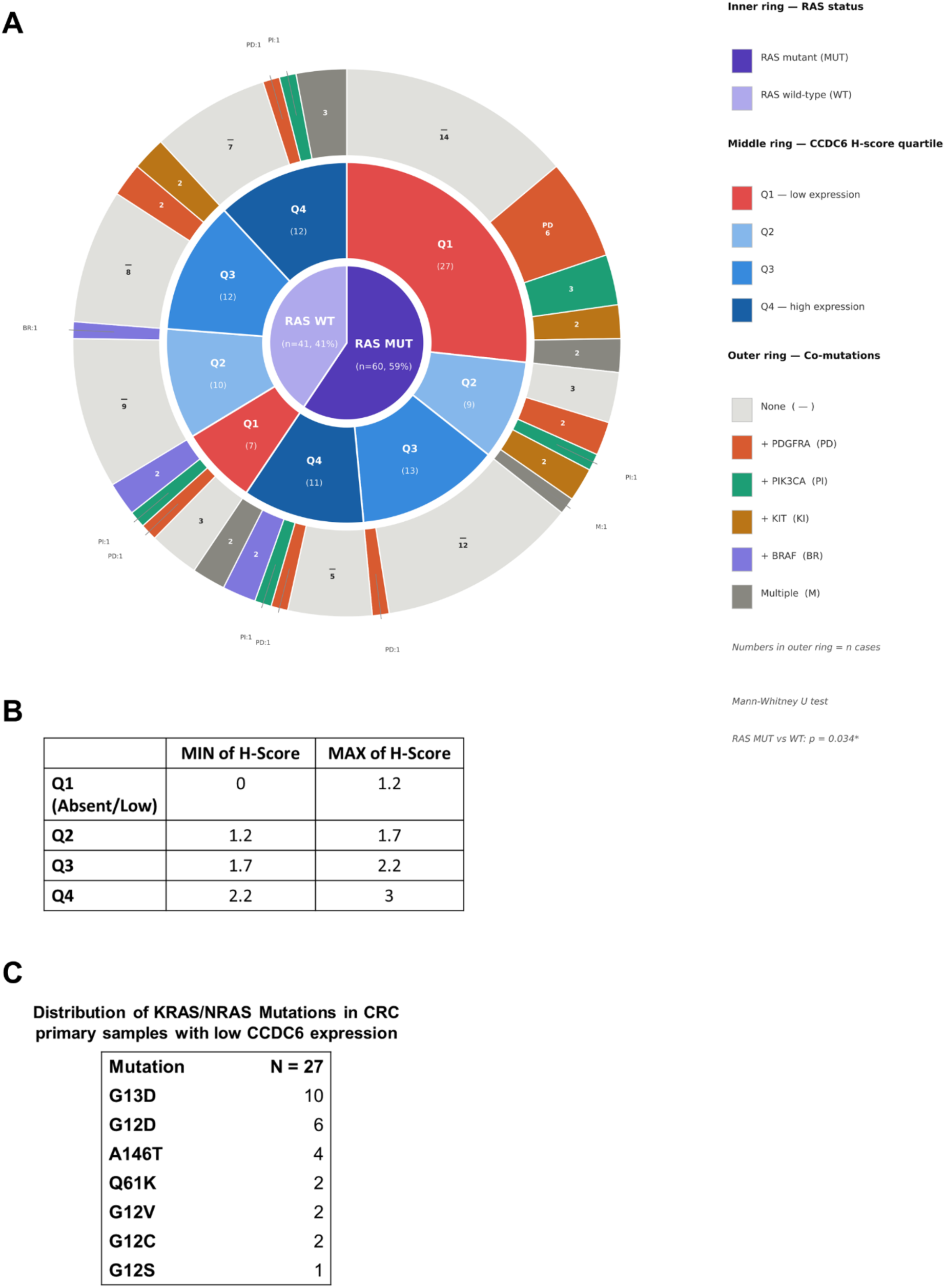
Distribution of CCDC6 expression according to RAS mutational status in Colorectal Cancer (CRC) primary samples. (A) Mutational landscape and CCDC6 protein expression in 101 colorectal cancer cases. The three-ring diagram summarises the co-occurrence of RAS (KRAS/NRAS) mutational status, CCDC6 immunohistochemical H-score quartiles, and concomitant gene alterations. Inner ring: RAS mutational status (MUT, n=60, 59%; WT, n=41, 41%). Middle ring: distribution of CCDC6 H-score quartiles (Q1-Q4, defined on the entire cohort) within each RAS group. Q1 = absent/lowest expression; 04 = highest expression. Outer ring: co-mutational profile (PDGFRA, PIK3CA, KIT, BRAF, or multiple alterations) within each quartile subgroup; grey segments indicate cases with no additional mutation among the tested genes. RAS MUT cases displayed significantly lower CCDC6 H-score than RAS WT cases (Mann-Whitney U test, p = 0.034). Abbreviations: MUT, mutant; WT, wild-type; H-score, IHC combinatory score. (B) The table defines the semi-quantitative scoring criteria used for patient stratification. The H-score range (0–3) was divided into quartiles: Q1 (0–1.2) represents Absent/Low expression, while Q2 (1.2–1.7), Q3 (1.7–2.2), and Q4 (2.2–3.0) represent increasing levels of CCDC6 protein expression. (C) The table displays the specific distribution of KRAS identified in the CRC samples with low CCDC6 expression.

**Table 1.**
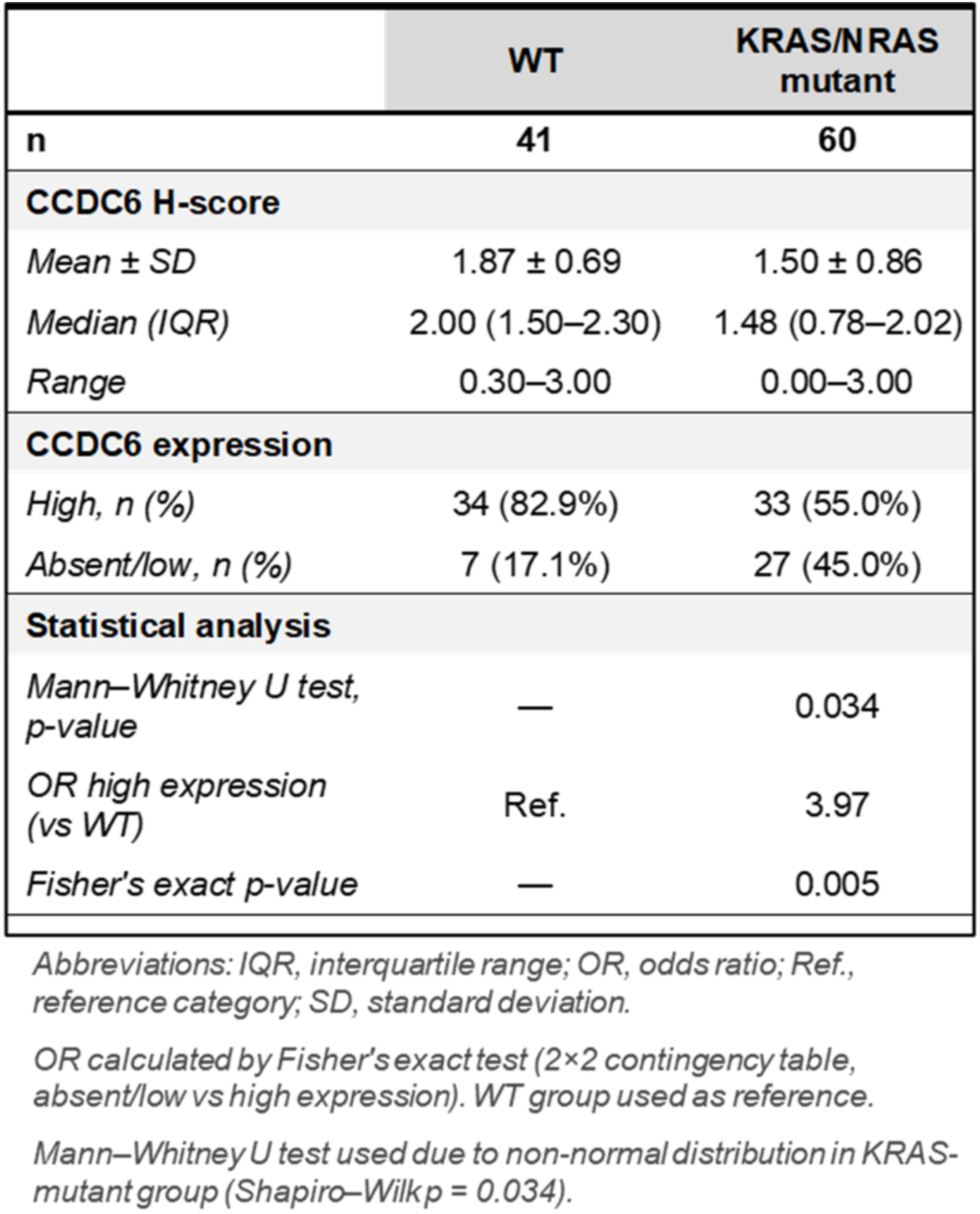
CCDC6 combinational score in RAS mutant vs WT samples. The table presents the distribution of CCDC6 protein expression, quantified by H-score (mean, SD, median, IQR, and range) and categorical expression (High vs. Absent/Low), stratified by RAS mutational status (WT vs. MUT). Differences between WT and MUT groups were analyzed using a two-sided Mann–Whitney U test (p = 0.034). The association between RAS status and CCDC6 expression levels was assessed by Fisher’s exact test (p = 0.005). The Odds Ratio (OR = 3.97) for high CCDC6 expression in the WT group versus the MUT group was 3.97, using the WT cohort as the reference category.

|  | WT | KRAS/NRAS mutant |
| --- | --- | --- |
| **n** | **41** | **60** |
| **CCDC6 H-score** |  |  |
| *Mean ± SD* | 1.87 ± 0.69 | 1.50 ± 0.86 |
| *Median (IQR)* | 2.00 (1.50–2.30) | 1.48 (0.78–2.02) |
| *Range* | 0.30–3.00 | 0.00–3.00 |
| **CCDC6 expression** |  |  |
| *High, n (%)* | 34 (82.9%) | 33 (55.0%) |
| *Absent/low, n (%)* | 7 (17.1%) | 27 (45.0%) |
| **Statistical analysis** |  |  |
| *Mann–Whitney U test, p-value* | — | 0.034 |
| *OR high expression (vs WT)* | Ref. | 3.97 |
| *Fisher's exact p-value* | — | 0.005 |

Abbreviations: IQR, interquartile range; OR, odds ratio; Ref., reference category; SD, standard deviation.
OR calculated by Fisher's exact test (2 × 2 contingency table, absent/low vs high expression). WT group used as reference.
Mann–Whitney U test used due to non-normal distribution in KRAS-mutant group (Shapiro–Wilk $p = 0.034$ ).

To further evaluate this effect, tumors were stratified according to quartiles of the composite score. Cases in the lowest quartile (Q1) were defined as absent/low-CCDC6 tumors. Absent/low expression was significantly enriched in the MUT group (27/60, 45%) compared with WT tumors (7/41, 17%), corresponding to an odds ratio (OR) of 3.97 (Fisher’s exact test, p = 0.005) (Figure 6 A, B, Table1). Interestingly, the distribution of KRAS/NRAS isoforms within the low-CCDC6 cohort reflects the frequency typically observed in colorectal tumors (Figure 6 C) [3].

Importantly, reduced CCDC6 expression in RAS mutated tumors was often observed as a heterogeneous intratumoral pattern. This phenotype was also observed in a minority of WT tumors, including cases harboring alterations in downstream signaling genes such as BRAF or PIK3CA, although these observations remain descriptive due to limited numbers (3 out of 41).

Collectively, our data indicate that CCDC6 downregulation in human CRC is characterized by reduced staining intensity, primarily driven by enhanced protein turnover, consistent with our cellular and preclinical findings.

## DISCUSSION

The clinical management of RAS-driven tumors remain a challenge due to their aggressive phenotype and therapeutic resistance [29]. Our study identifies a novel signaling axis linking KRAS activation to the proteasomal degradation of CCDC6, which in turn promotes an antioxidant program via ATF4-mediated xCT upregulation, conferring resistance to ferroptosis.

We show that KRAS-driven MAPK signaling triggers GSK3β activation, which is required for CCDC6 degradation. While CCDC6 impairment is documented across various cancers [30], the upstream triggers for its turnover remained elusive [13, 15]. Our work fills this gap, establishing a direct link between oncogenic RAS and the physical/functional loss of CCDC6.

Beyond its known role in proliferation, KRAS profoundly alters cellular redox balance. It is well established that KRAS bolsters defenses against oxidative stress by upregulating xCT/SLC7A11 to increase glutathione (GSH) synthesis [9, 10]. Our findings refine this model by revealing a previously unrecognized KRAS-CCDC6-ATF4 axis. We demonstrate that CCDC6 physically interacts with ATF4, preventing its binding to the xCT promoter. Consequently, KRAS-induced CCDC6 loss "unleashes" ATF4, driving xCT expression, GSH accumulation, and ferroptosis evasion. This causal link was validated by re-expressing CCDC6 in KRAS-mutant cells, which restored sensitivity to ferroptotic agents, a finding consistently reproduced across both endogenous mutants and transfected cellular models.

Interestingly, our analysis of human CRC cohorts and PDX models suggests that CCDC6 loss is not exclusive to KRAS mutations. We observed low CCDC6 expression in approximately 15% of KRAS wild-type tumours. While some of these cases harbor BRAF or PI3K alterations—suggesting a convergent MAPK/PI3K signaling node—others might be driven by cell-autonomous EGFR signaling [31, 32]. Indeed, CCDC6 negatively regulates AREG via CREB1 [16, 33]; thus, an EGFR-driven feedback loop might promote CCDC6 turnover even in the absence of RAS mutations. Further studies are needed to confirm if EGFR inhibitors like Gefitinib could rescue CCDC6 levels in these specific subsets.

The translational implications of our study are significant. We demonstrate that CCDC6 levels can be rescued by inhibiting its degradation (via Bortezomib or the GSK3β inhibitor SB216763) or by direct KRAS inhibition (G12C-Sotorasib, Adagrasib, or the novel G12D-specific HRS4642). Restoring CCDC6 levels re-sensitizes KRAS-mutated cells to Sulfasalazine (SSZ), an FDA-approved drug for rheumatic diseases [34]. Notably, the recent development of allele-specific inhibitors like HRS4642 offers a breakthrough for G12D-mutant PDAC and CRC, where G12C inhibitors have shown limited efficacy compared to NSCLC due to adaptive feedback loops.

Finally, the loss of CCDC6 provides a "double hit" to cancer cell survival. Beyond ferroptosis resistance, we confirm that CCDC6 loss impairs homologous recombination (HR) repair, as evidenced by reduced γH2AX/RAD51 foci and increased Olaparib sensitivity in gastric cancer models [Figure S6]. This suggests that CCDC6 could serve as a dual biomarker: predicting resistance to ferroptosis and sensitivity to PARP inhibitors.

In conclusion, our study uncovers a KRAS-driven CCDC6/ATF4/xCT signaling axis that governs ferroptosis resistance. By identifying CCDC6 turnover as a targetable vulnerability, we provide a rationale for combining KRAS or proteasome inhibitors with ferroptosis-inducing agents, potentially expanding the therapeutic toolkit for RAS-mutated malignancies.

## MATERIALS AND METHODS

### Cell lines, drugs and chemicals

Human Non-Small Cell Lung Cancer (NSCLC) cell lines (H1975, RRID:CVCL_1511; H460, RRID:CVCL_0459; HOP62; RRID:CVCL_1285), were cultured in RPMI 1640 (Gibco, Paisley, UK). Human colorectal (GEO, RRID:CVCL_1227), gastric (AGS; RRID:CVCL_0139), pancreatic (PANC1, RRID:CVCL_0480) cancer cell lines, embryonic kidney cells (293T, RRID:CVCL_0063) and murine pancreatic cancer cells lines (FC1199) were cultured in DMEM (Gibco, Paisley, UK). The FC1199 pancreatic cancer cell line, derived from tumors arisen in *LSL-KrasG12D/+;LSL-Trp53R172H/+;Pdx-1-Cre* mice in the C57BL/6 background, was provided by D.A. Tuveson (Cold Spring Harbor, NY, USA). All media were supplemented 10% fetal bovine serum (Gibco, Paisley, UK), 1% penicillin/streptomycin (Gibco, Paisley, UK). Erastin (E7781), Sulfasalazine (S0883), MG132 (474790), ferrostatin-1 (SML0583), deferoxamine (D9533), GSK3*β* inhibitor SB216763 (S3442), H2O2 (H1009), Diethyl maleate (DEM) (D97703) and Hoechst-33258 (94403) were provided by Sigma-Aldrich, Inc (St. Louis, CA, USA). Adagrasib (S8884), Sotorasib (S8830), HRS4642 (E4651), MEK1/2 inhibitor Selumetinib (AZD6244), Bortezomib (S1013), USP7 inhibitor P5091, and Olaparib (AZD2281) were from SelleckChem. Z-VAD-fmk (FMK001) was from MedChemExpress.

### Reagents and antibodies

For the biochemical and immunofluorescence analysis the following antibodies were utilised: anti-CCDC6 (Abnova, Cat# H00008030-M03, RRID:AB_529979); anti-tubulin (Sigma-Aldrich, Cat# T6557, RRID:AB_477584); anti-xCT/SLC7A11 (Cell signaling; Cat# 12691, RRID:AB_2687474); anti-Phospho-p44/42 MAPK (Erk1/2) (Cell signaling, Cat# 4370, RRID:AB_2315112); anti-p44/42 MAPK (Erk1/2) (Cell signaling, Cat# 4695, RRID:AB_390779); anti-Gsk3*β* (Cell signaling, Cat# 9315, RRID:AB_490890); anti-FLAG (DYKDDDDK Tag 9A3) (Cell signaling, Cat# 8146 RRID:AB_10950495); anti-ATF4 (Cell signaling, Cat# 11815, RRID:AB_2616025); anti-Myc (Santa Cruz, Cat# sc-40, RRID:AB_627268); anti-cyclin E (Santa Cruz, Cat# sc-247, RRID:AB_627357); anti-RAD51 (Abcam, Cat# ab133534, RRID:AB_2722613); anti-γH2AX (Merck Millipore, Cat# 05-636, RRID:AB_309864). Secondary antibodies conjugated to horseradish peroxidase were provided by Bio-Rad (anti-mouse Cat# 170-5047, RRID:AB_11125753; anti-rabbit Cat# 170-5046, RRID:AB_11125757). Fluorescently labeled secondary antibodies were provided by Abcam (Alexa Fluor 594, Cat# ab150116, RRID:AB_2650601; Alexa Fluor 488, Cat# ab150077, RRID:AB_2630356).

### Plasmids and transfection

Plasmids containing the mutated RAS isoforms (RRID:Addgene_1000000089) were cloned into pDEST27 (RRID:Addgene_118371) using Gateway LR Clonase II Enzyme Mix (Thermo Fisher Scientific, Cat# 11791020). The pcDNA4/myc-His A empty vector (Invitrogen, Cat# V1030-20, RRID:SCR_008450) and the pcDNA4ToA-mycCCDC6wt (wild-type) construct were utilized for transfections. pxCT-pro-WT-Luc (human xCT gene promoter-luciferase reporter was a gift from Ken Itoh (Irosaki University, Japan). Sh-FBXW7 was kindly provided by Wenye Wei [15]. pRK-ATF4 plasmid was obtained from Addgene (RRID:Addgene_26114). The plasmids were transfected with the FuGene HD (Promega) reagent.

### Western Blot and co-immunoprecipitation (Co-IP)

Total protein extracts were obtained using RIPA buffer (50 mM Tris–HCl pH 7.5, 150 mM NaCl, 1% Triton X-100, 0.5% Na Deoxycholate, 0.1% SDS) with a mix of protease inhibitors (Roche). Protein concentration was estimated by Bradford assay (Biorad). For Western blotting, protein samples were separated by 10% SDS-PAGE and transferred to a PVDF membrane. Membranes were blocked with 5% TBS-BSA and incubated with the primary antibodies. Immunoblotting experiments were carried out according to standard procedures and visualised using Bio Rad ChemiDoc MP Imaging System (RRID:SCR_019037). For co-immunoprecipitation, HEK293T cells were transfected with mycCCDC6 and ATF4 expression plasmids. After 24h, cells were treated with 20 µM MG132 for 6h and lysed in lysis buffer (Tris-HCl ph 7.5, 10mM, NaCl 150 mM, EDTA 0,1 mM, NP40 0,5%). Following centrifugation, 3.4mg of protein were incubated O.N with anti-Myc antibody at 4°C. Antigen–antibody complexes were bound to Protein G–Sepharose (Santa Cruz Biotechnology) for 1h at 4 °C with rocking. Non-specific interactions were removed by extensive washing and bound proteins were eluted by boiling in 2× SDS sample loading buffer and subjected to SDS-PAGE for immunoblot analysis.

### Real time PCR

PCR reactions were performed on RNA isolated from cell lines using Trizol reagent (Life Technologies) and reverse-transcribed using iScript RT supermix (Bio-Rad). Quantitative real time (qRT-PCR) was performed with iTaq Universal SYBR Green Supermix (Bio-Rd) using the following primers: CCDC6 forward 5’-ggagaaagaaacccttgctg-3’ and reverse 5’-tcttcatcagtttgttgacctga-3’; xCT/SLC7A11 forward 5’-gcgtgggcatgtctctgac-3’ and reverse 5’-gctggtaatggaccaaagacttc-3’; β-Actin forward 5’-tgcgtgacattaaggagaag-3’ and reverse 5’-gctcgtagctcttctcca-3’.

### Sensitivity Test and Design for Drug Combination

The CellTiter 96 AQueous One Solution Cell Proliferation Assay (Promega) was used to determine cell viability and drug IC50 values [11]. Drug combination analysis was conducted using CompuSyn software (RRID:SCR_022931) to calculate the Combination Index (CI) to define synergistic (<1), additive (=1), or antagonistic (>1) interactions, and the Dose Reduction Index (DRI) to quantify the dose reduction achievable in synergistic combinations

### GSH assay

GSH levels were detected using a GSH-Glo Glutathione Assay (Promega) following the manufacturer’s instructions. Briefly, 3000 cells per well were seeded in a 96-well plate 24h before analysis. The culture medium was removed and 100 μl of 1× GSH-Glo Reagent was added to each well, followed by incubation at room temperature for 30 min. Next, 100 μl of reconstituted Luciferin Detection Reagent was added. After 20 min, luminescence was measured using Synergy H1 microplate reader (BioTek, RRID:SCR_019748). Cellular GSH concentrations were calculated based on a GSH standard curve according to the manufacturer’s instructions.

### Lipid peroxidation assay

Lipid peroxidation levels were measured by BODIPY 581/591 C11 dye (D3861, Invitrogen). Briefly, cells were incubated in a 6-well plate containing 5 μM BODIPY. After incubation for 30 min at 37°C, cells were washed with PBS and trypsinized, then subjected to flow cytometry analysis using a Miltenyi MACSQuant Analyzer 10 - Flow Cytometer (Miltenyi Biotec, RRID:SCR_020268).

### Cystine uptake assay

The cystine uptake assay was performed using BioTracker Cystine-FITC Live Cell Dye (#SCT047, Sigma). Briefly, cells were seeded in a 6-well plate for 24h and treated with 1mM DEM for 3h. After incubation with 5 μM Cystin-FTC for 30 min at 37°C, cells were washed with PBS, trypsinized, and analyzed by flow cytometry using a Miltenyi MACSQuant Analyzer 10 (Miltenyi Biotec, RRID:SCR_020268).

### Luciferase assay

NSCLC cells were seeded in 12-well plates and co-transfected with pxCT-pro-WT-Luc and Renilla plasmids, in the presence or absence of pRK-ATF4 and MycCcdc6. After 24h, cells were harvested and luciferase activity was measured using the Dual-Luciferase Reporter Assay System (#E1910, Promega) following the manufacturer’s instructions. Firefly luciferase luminescence was measured after adding Luciferase Assay Reagent II. Subsequently, Renilla luciferase luminescence was read after adding Stop & Glo Reagent. All luminescence readings were performed on a Bio-Tek Synergy H1 Microplate Reader (BioTek, RRID:SCR_019748), and final activity was expressed as the ratio of Firefly to Renilla luminescence.

### Chromatin Immunoprecipitation (ChIP)

H460 and HOP62 cells were treated with hydrogen peroxide (tempistiche) and fixed with 1% formaldehyde for 10 min at 37°C, followed by reaction quenching with glycine. The extracts were lysed and sonicated to produce chromatin suspensions of DNA approximately 300 bp in length. Subsequently, the lysates were incubated with anti-ATF4 antibody overnight at 4°C. Normal rabbit IgG was used as a negative control. After antigen-antibody reaction, immunocomplexes were captured with Protein A–Sepharose (Santa Cruz Biotechnology), and coimmunoprecipitated DNA fragments were purified. The relative amounts of immunoprecipitated DNA fragments were evaluated by qPCR using the following primer pair: xCT/SLC7A11 forward 5’-ttgagcaacaagctcctcct-3’ and reverse 5’-caaaccagctcagcttcctc-3’. The results are presented as the ratio to input DNA.

### Immunofluorescence staining

Following genotoxic stress exposure (10μM etoposide for 4h), cells were fixed with 4% PFA, permeabilized with 0.05% Triton-PBS and blocked with 1% BSA-PBS. After staining with primary antibodies, cells were washed in PBS and incubated for 30 min at room temperature with fluorescent secondary antibody. Nuclei were visualized by staining with Hoechst. Cells with ≥ 5 distinct γH2AX or RAD51 foci were considered positive. The percentage of positive nuclei was calculated from a minimum of 250 analyzed nuclei per sample.

### Immunocytochemical and immunohistochemical analyses

Murine FC1199 pancreatic cancer cells harboring the KRASG12D mutation were cultured under standard conditions and subjected to pharmacological treatments with the proteasome inhibitor MG132 or vehicle control (DMSO). Following treatment, cells were processed for cell block preparation by formalin fixation and paraffin embedding.

Immunocytochemical (ICC) analysis was performed on cell block sections using an anti-CCDC6 antibody, followed by detection with a diaminobenzidine (DAB)-based chromogenic system.

For in vivo studies, FC1199 cells were orthotopically implanted into the pancreas of immunocompromised mice to generate xenograft tumors. Tumor and adjacent normal pancreatic tissues were collected, formalin-fixed, and paraffin-embedded. Immunohistochemical (IHC) staining for CCDC6 was performed using a DAB-based detection system. Patient-derived xenograft (PDX) samples from pancreatic ductal adenocarcinoma (PDAC) cases harboring KRASG12D or KRASG12R mutations (n = 4), as well as one KRAS wild-type case, were processed using the same protocol. Additional IHC staining for phospho-MAPK (pMAPK) was performed to assess downstream RAS pathway activation. Normal pancreatic ductal tissue was processed and stained, as control (Figure S4).

In a parallel study, 101 primary human colorectal carcinoma (CRC) samples were screened for RAS mutations and microsatellite status (MSS vs. MSI) using Next-Generation Sequencing (NGS). The 101 samples were microsatellite stable (MSS). 60 exhibited RAS mutations (55 KRAS and 5 NRAS), while the remaining 41 were RAS wild type. Additional mutations identified via NGS are reported in the Table S1. All the primary FFPE colorectal carcinoma (CRC) specimens were retrospectively analyzed. CCDC6 protein expression was assessed on tumor sections by two independent pathologists (EV, FM) blinded to mutational status. Immunohistochemical evaluation was performed using a semi-quantitative scoring system integrating both staining intensity and the proportion of positive tumor cells within neoplastic areas. Staining intensity was categorized as 1+ (weak), 2+ (moderate), or 3+ (strong), and the percentage of tumor cells at each intensity level was recorded.

A composite semi-quantitative H-score was calculated for each case using the formula: H-score = (1 × %1+ + 2 × %2+ + 3 × %3+)/100, yielding a continuous variable ranging from 0 to 3. Cases with complete absence of staining were assigned a score of 0. Tumors were stratified according to RAS mutational status into MUT (n = 60) and WT (n = 41) groups.

For categorical analyses, tumors were further classified based on quartiles of the H-score distribution. Cases falling within the lowest quartile (Q1), together with tumors showing complete absence of staining, were defined as “absent/low CCDC6 expression.” Subgroup analyses were performed restricting the MUT cohort to KRAS-mutant cases.

Digital image analysis was conducted using QuPath (version 0.6.0) [35]. Color deconvolution was applied to isolate the DAB signal, and optical density (OD) measurements were extracted. For ICC experiments, CCDC6 expression was expressed as the ratio between treated and control conditions (MG132/DMSO). For tissue samples, expression levels were quantified as tumor-to-normal DAB OD ratios. All analyses were performed using standardized thresholds and batch processing to ensure consistency.

### Statistical analysis

Statistical analysis was performed using GraphPad Prism version 10 software (RRID: SCR_002798). All data are presented as mean ± SEM. Student’s t-test was used for comparative analysis between data groups. The following asterisk rating system for P values was used: *, P < 0.05;**, P < 0.01; ***, P < 0.001; ****, P < 0.0001.

Regarding immunohistochemical (IHC) analysis, statistical evaluations were conducted using non-parametric methods due to non-normal distribution of H-scores. Differences between groups were assessed using the two-sided Mann–Whitney U test. Associations between categorical variables were evaluated using Fisher’s exact test, and odds ratios (ORs) with corresponding p-values were calculated to estimate effect size. A p-value < 0.05 was considered statistically significant.

