## Supplementary_Figures S1-S7 for "KRAS-Mediated CCDC6 Degradation Drives xCT Upregulation and Ferroptosis Evasion"

### Figure S1

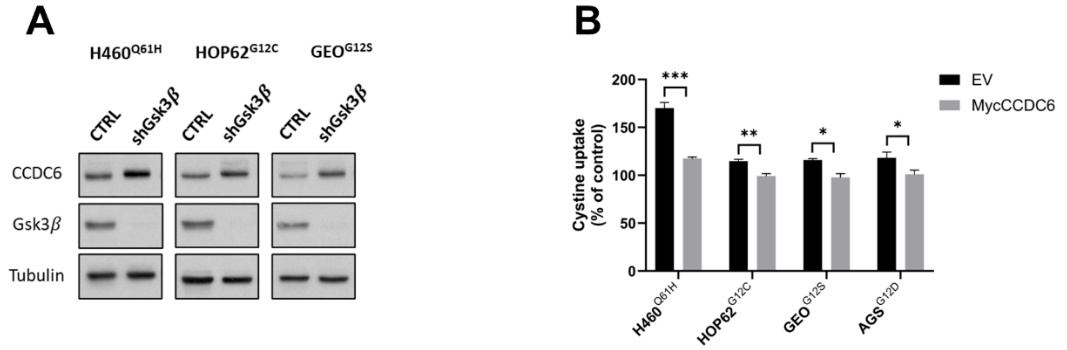


**Figure S1: Regulation of CCDC6 stability by Gsk3$\beta$ and impact of CCDC6 re-expression on cystine uptake.**
(A) Western blot analysis of CCDC6 expression in H460, HOP62 and GEO cells upon transient transfection with shGsk3$\beta$. Anti-Gsk3$\beta$ and anti-tubulin immunoblots are shown as transfection and loading control, respectively.
(B) Cystine uptake was assessed in H460, HOP62, GEO and AGS cells transfected with MycCCDC6 or empty vector (EV) and treated with Diethyl Malonate (DEM) [1mM] for 3h by a BioTracker Cystine-FITC Live Cell Dye followed by flow cytometry analysis. Data are reported as mean ± SEM of 3 independent repeats. Statistical significance was verified by 2-tailed Student's t-test (* $p < 0.05$; ** $p < 0.01$, *** $p < 0.001$).

### Figure S2

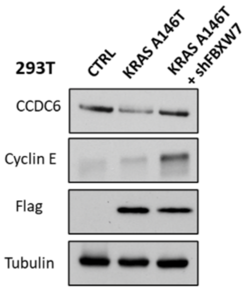


**Figure S2: FBXW7 involvement in KRAS-mediated CCDC6 degradation.**
Western blot analysis of CCDC6 expression in 293T cells transfected with KRAS isoform A146T or empty vector (CTRL), in presence or absence of shFBXW7. Expression levels of cyclin E are shown as a representative substrate of FBXW7. Anti-flag immunoblots was performed to confirm the expression of Flag-KRAS mutant plasmid. Anti-tubulin is shown as loading control.

### Figure S3

A

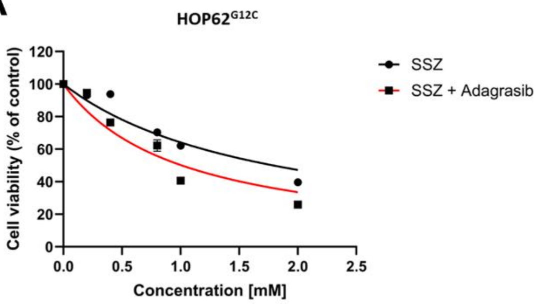


| Drug sensitivity (IC50) HOP62 Adagrasib [2µM] | - | + | Combination index (CI) | Dose Reduction Index (DRI) |
|---|---|---|---|---|
| SSZ [mM] | 1.79 +/- 0.15 | 1.01 +/- 0.09 | 0.78 | 2.04 |

B

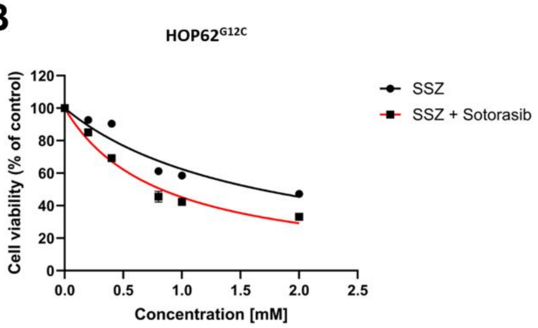


| Drug sensitivity (IC50) HOP62 Sotorasib [40µM] | - | + | Combination index (CI) | Dose Reduction Index (DRI) |
|---|---|---|---|---|
| SSZ [mM] | 1.67 +/- 0.11 | 0.82 +/- 0.04 | 0.68 | 2.08 |

C

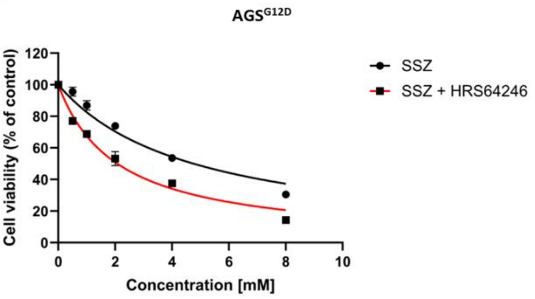


| Drug sensitivity (IC50) AGS HRS-4642 [6nM] | - | + | Combination index (CI) | Dose Reduction Index (DRI) |
|---|---|---|---|---|
| SSZ [mM] | 4.76 +/- 0.32 | 2.08 +/- 0.12 | 0.58 | 1.57 |

D

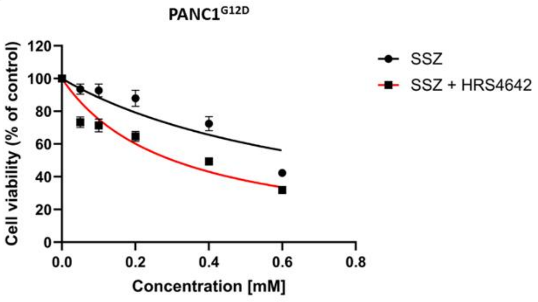


| Drug sensitivity (IC50) PANC1 HRS-4642 [5µM] | - | + | Combination index (CI) | Dose Reduction Index (DRI) |
|---|---|---|---|---|
| SSZ [mM] | 0.76 +/- 0.10 | 0.3 +/- 0.03 | 0.72 | 1.89 |

**Figure S3: KRAS inhibitors and Sulfasalazine effect on cell viability.**
(A-D) The synergistic effect of combining Sulfasalazine (SSZ) with KRAS inhibitors was evaluated using dose-response curves. IC50 values for Sulfasalazine (SSZ) were determined in the absence or presence of a fixed concentration of KRASG12C inhibitors (Adagrasib or Sotorasib) in HOP62 cells (A, B) and the KRASG12D inhibitor (HRS4642) in AGS (C) and PANC1 (D) cells for 24h. The synergy was confirmed by calculating the Combination Index (CI) and the Dose Reduction Index (DRI) from the same curves.

### Figure S4

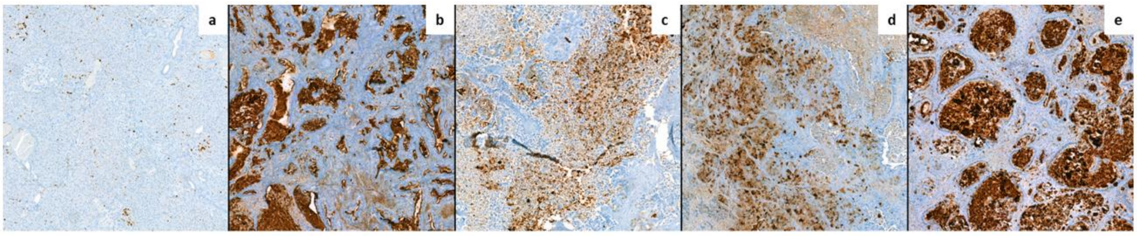


**Figure S4: Immunohistochemical (IHC) analysis of pMAPK expression in Patient-Derived Xenografts (PDX) of PDAC**.

Representative images showing pMAPK protein levels in normal pancreatic ductal tissue (a) and in PDX samples with different KRAS status: KRAS wild type (b), KRAS G12D (c), KRAS G12D (d), KRAS G12R (e). Scale bar = 100 µm.

### Figure S5

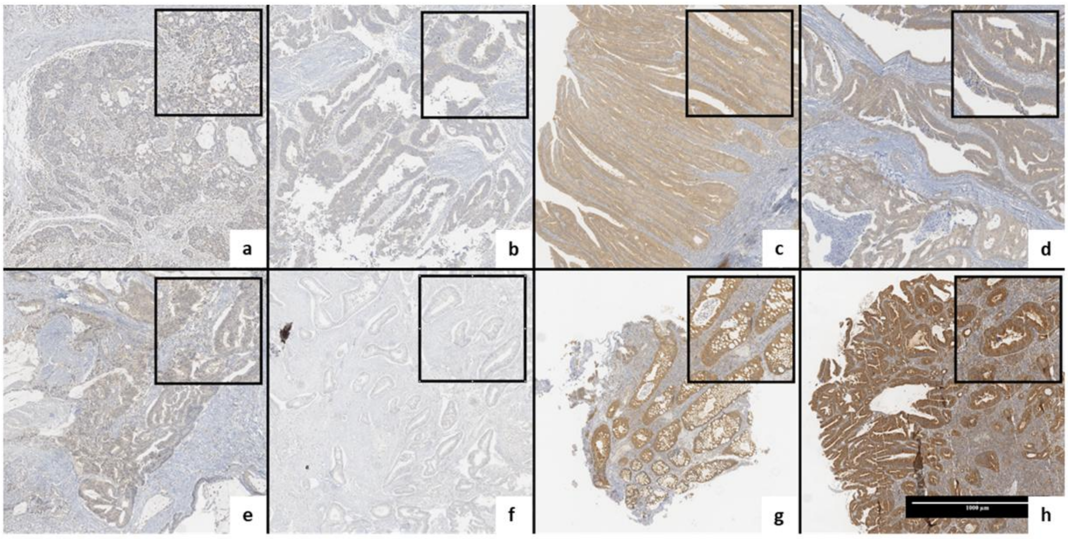


**Figure S5: Immunohistochemical (IHC) analysis of CCDC6 expression in primary colorectal carcinoma (CRC) samples.**

Representative images showing CCDC6 protein levels in KRAS wild-type (WT) cases with Absent/Low (a, b) or High (c, d) CCDC6 expression, and in KRAS mutant (MUT) cases with Absent/Low (e, f) or High (g, h) CCDC6 expression. Scale bar = 1000 µm.

### Figure S6

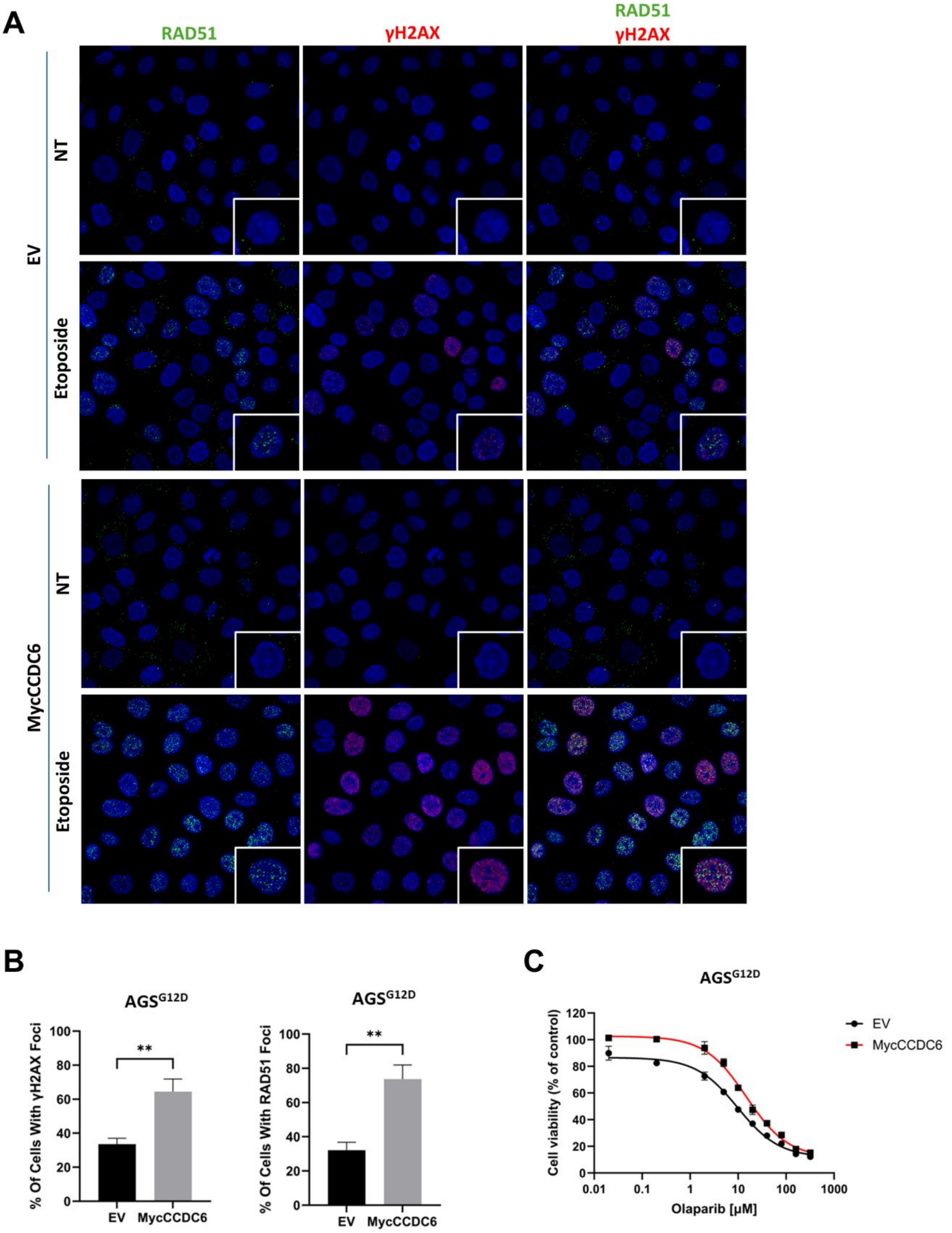


| Drug sensitivity (IC50) | |
|---|---|
| AGS | Olaparib [μM] |
| EV | 10.15 +/- 1.22 |
| MycCCDC6 | 22.95 +/- 1.56 |

**Figure S6: Re-expression of CCDC6 restores HR repair efficiency and induces PARP inhibitor resistance in AGS$^{G12D}$ cells.**

(A) Representative immunofluorescence images of AGS cells transiently transfected with MycCCDC6 or empty vector (EV) as a control. Cells were either left untreated (NT) or treated with Etoposide [10 μM] for 4h (to induce DNA damage) and stained for γH2AX (red), RAD51 (green), and DAPI (blue) to visualize DNA damage foci formation and the nucleus, respectively.
(B) Bar graphs represent the percentage of AGS cells with more than 5 foci. Data are reported as mean ± SEM of 3 independent repeats. Statistical significance was verified by 2-tailed Student's t-test (** $p < 0.01$).
(C) Dose-response curves were generated to assess the effect of CCDC6 re-expression on AGS cell viability in the presence of increasing concentrations of the PARP inhibitor Olaparib for 96h. MycCCDC6 re-expression was compared to an empty vector (EV) control. The IC50 values were calculated for each condition. The table (bottom) summarizes the calculated IC50 values ± SD.

### Figure S7

Original Western Blot Relative to Figure 1A

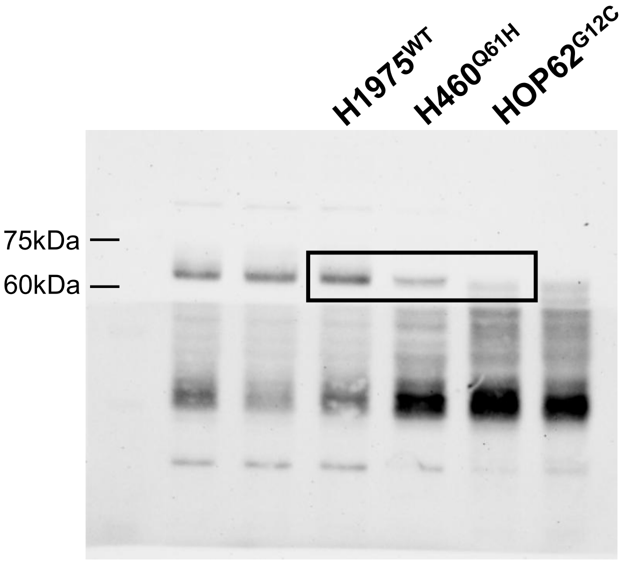


**CCDC6**

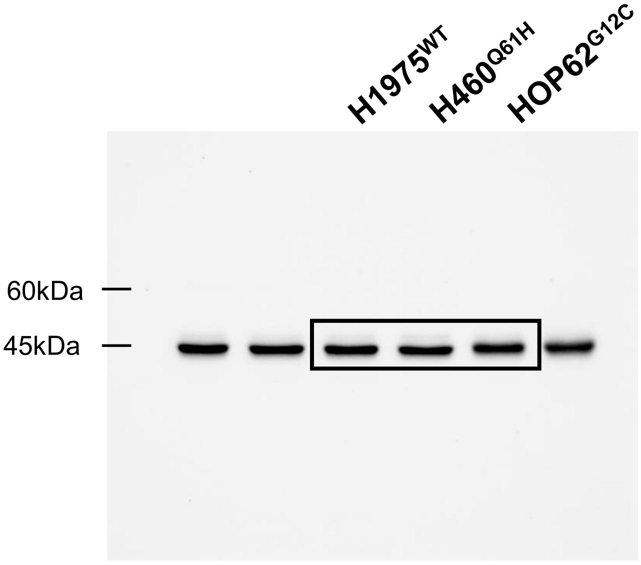


**Tubulin**

### Original Western Blot Relative to Figure 1C

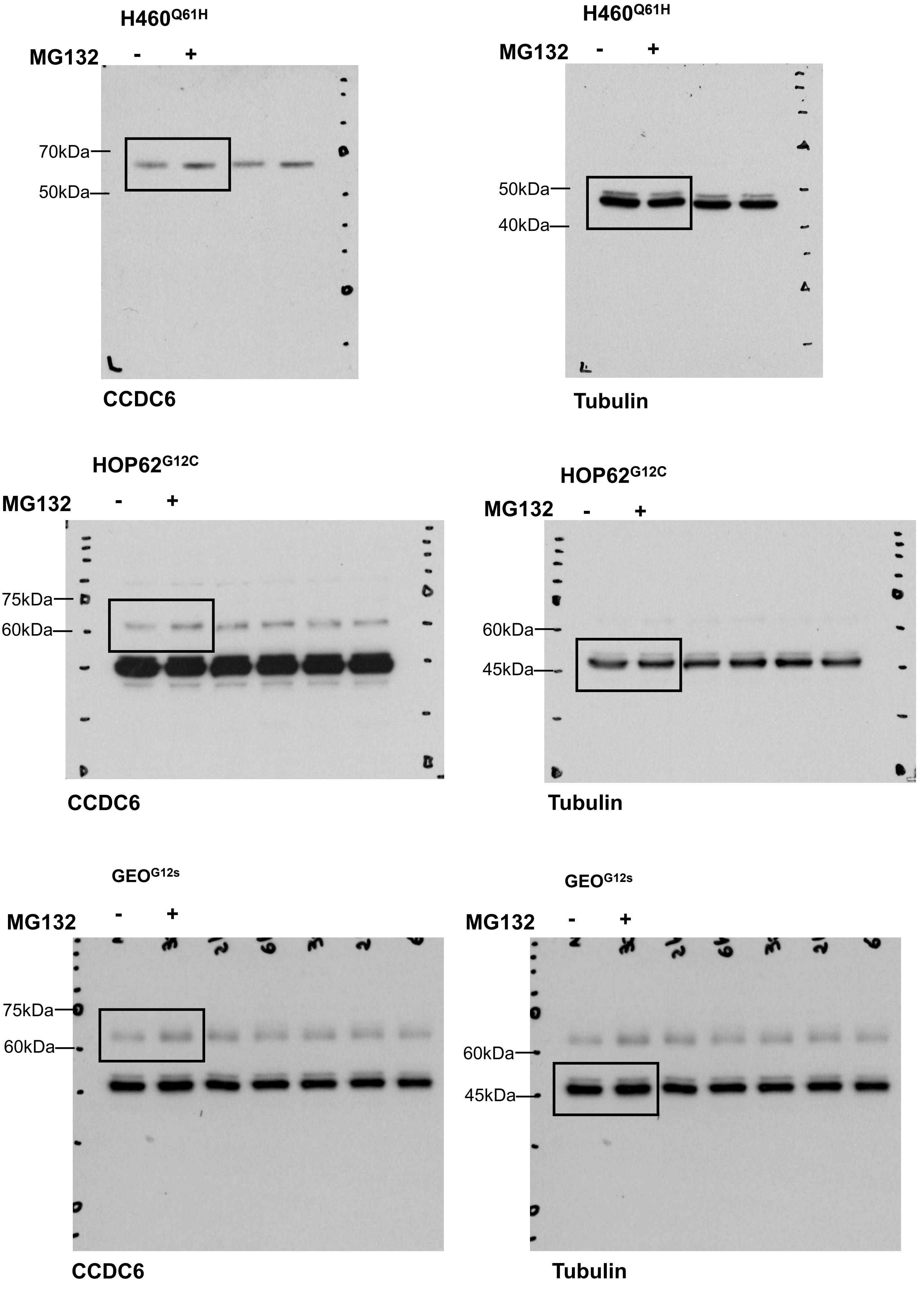

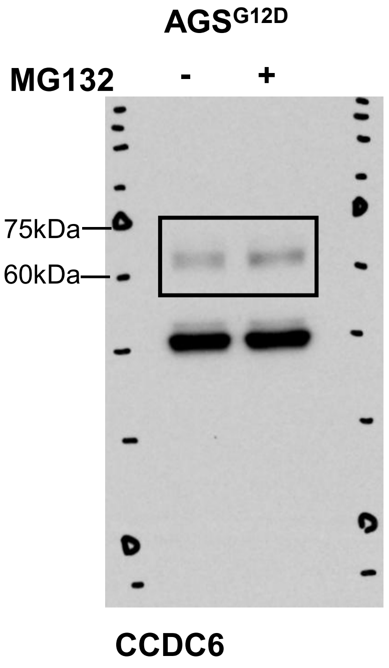
AGS$^{G12D}$
MG132 - +
75kDa
60kDa
CCDC6

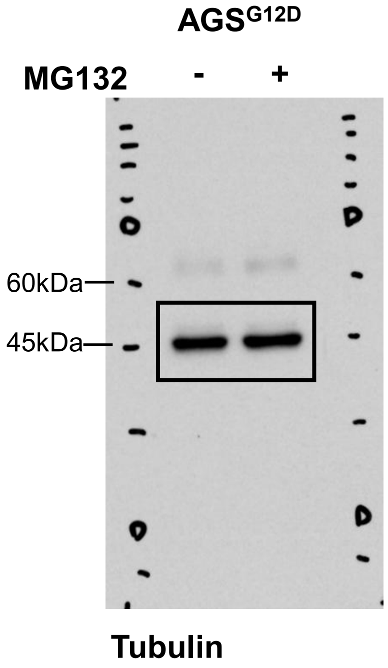
AGS$^{G12D}$
MG132 - +
60kDa
45kDa
Tubulin

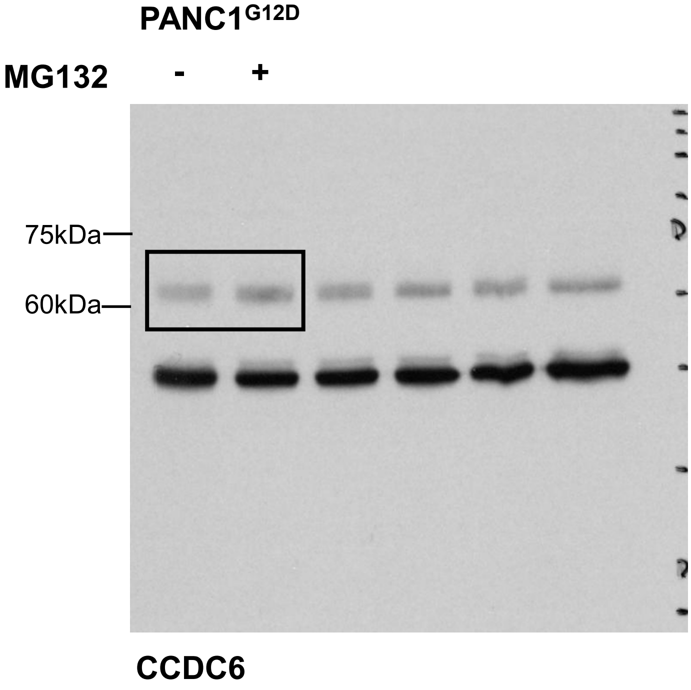
PANC1$^{G12D}$
MG132 - +
75kDa
60kDa
CCDC6

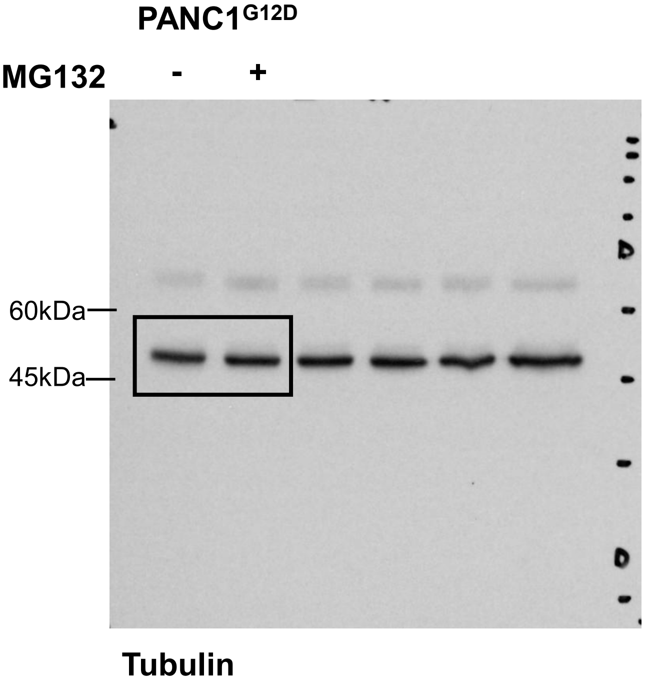
PANC1$^{G12D}$
MG132 - +
60kDa
45kDa
Tubulin

Original Western Blot Relative to Figure 1D

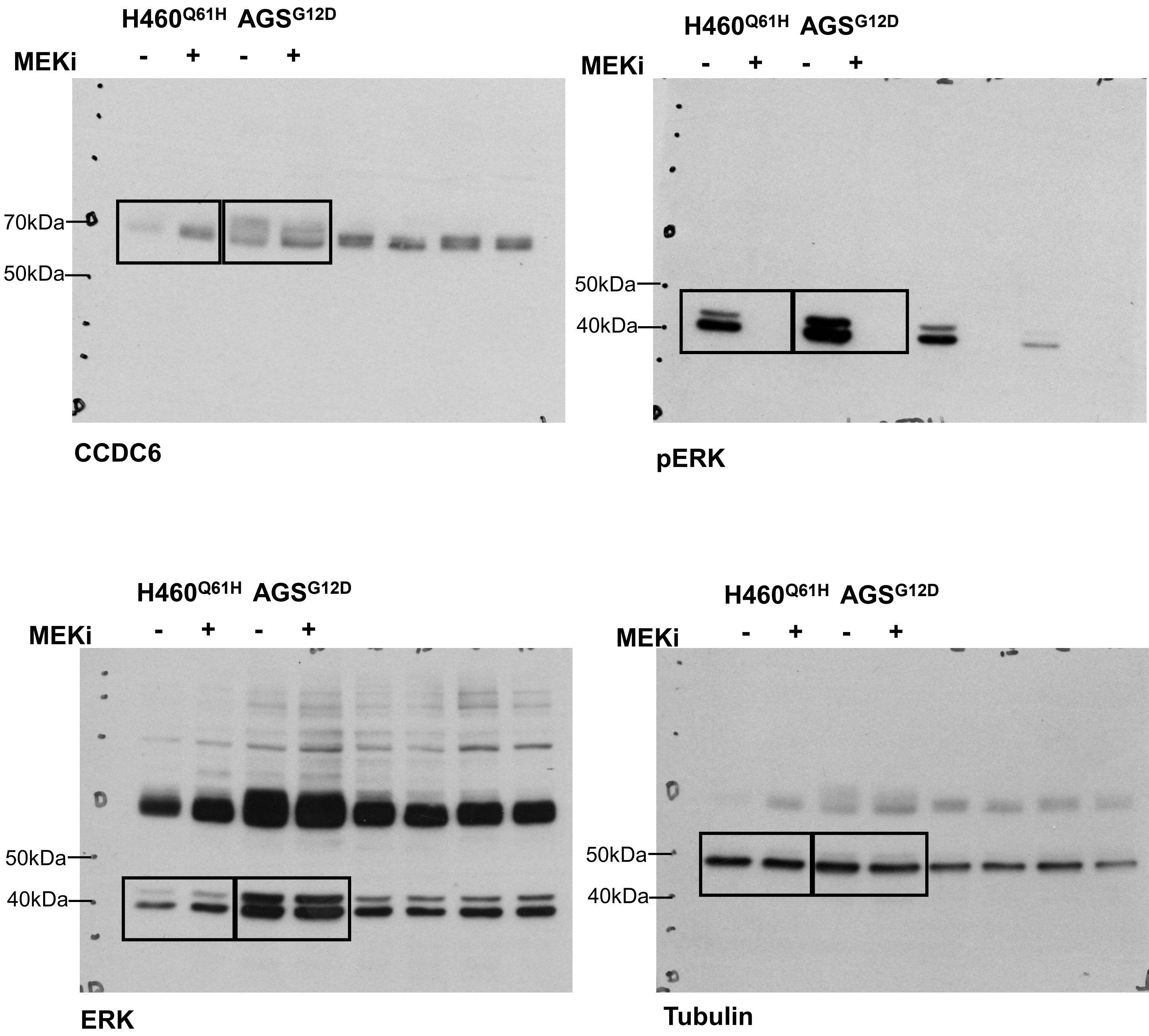

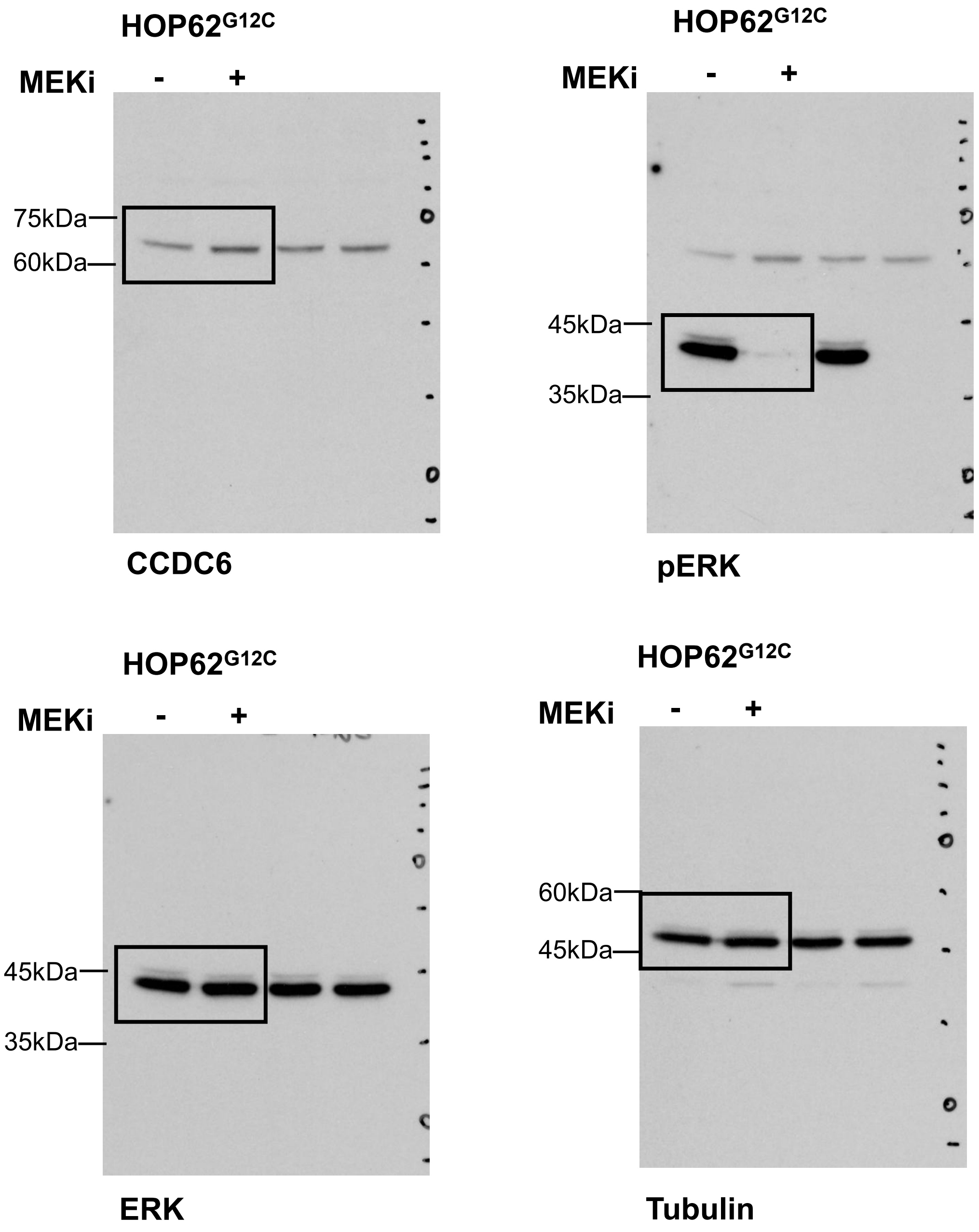

HOP62$^{G12C}$
MEKi - +
75kDa
60kDa
CCDC6
HOP62$^{G12C}$
MEKi - +
45kDa
35kDa
pERK
HOP62$^{G12C}$
MEKi - +
45kDa
35kDa
ERK
HOP62$^{G12C}$
MEKi - +
60kDa
45kDa
Tubulin

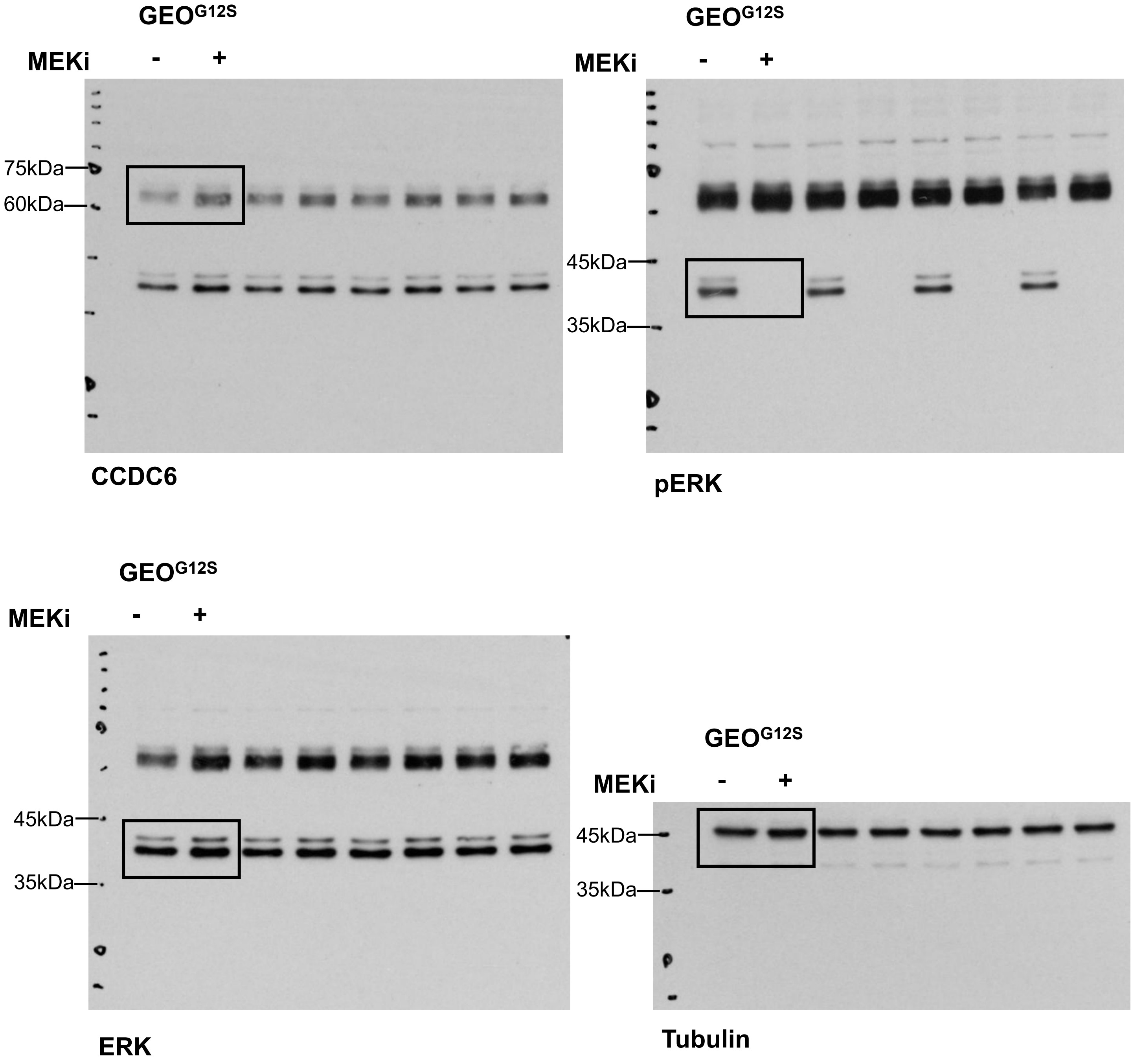

GEO G12S
MEKi - +
75kDa
60kDa
CCDC6
GEO G12S
MEKi - +
45kDa
35kDa
pERK
GEO G12S
MEKi - +
45kDa
35kDa
ERK
GEO G12S
MEKi - +
45kDa
35kDa
Tubulin

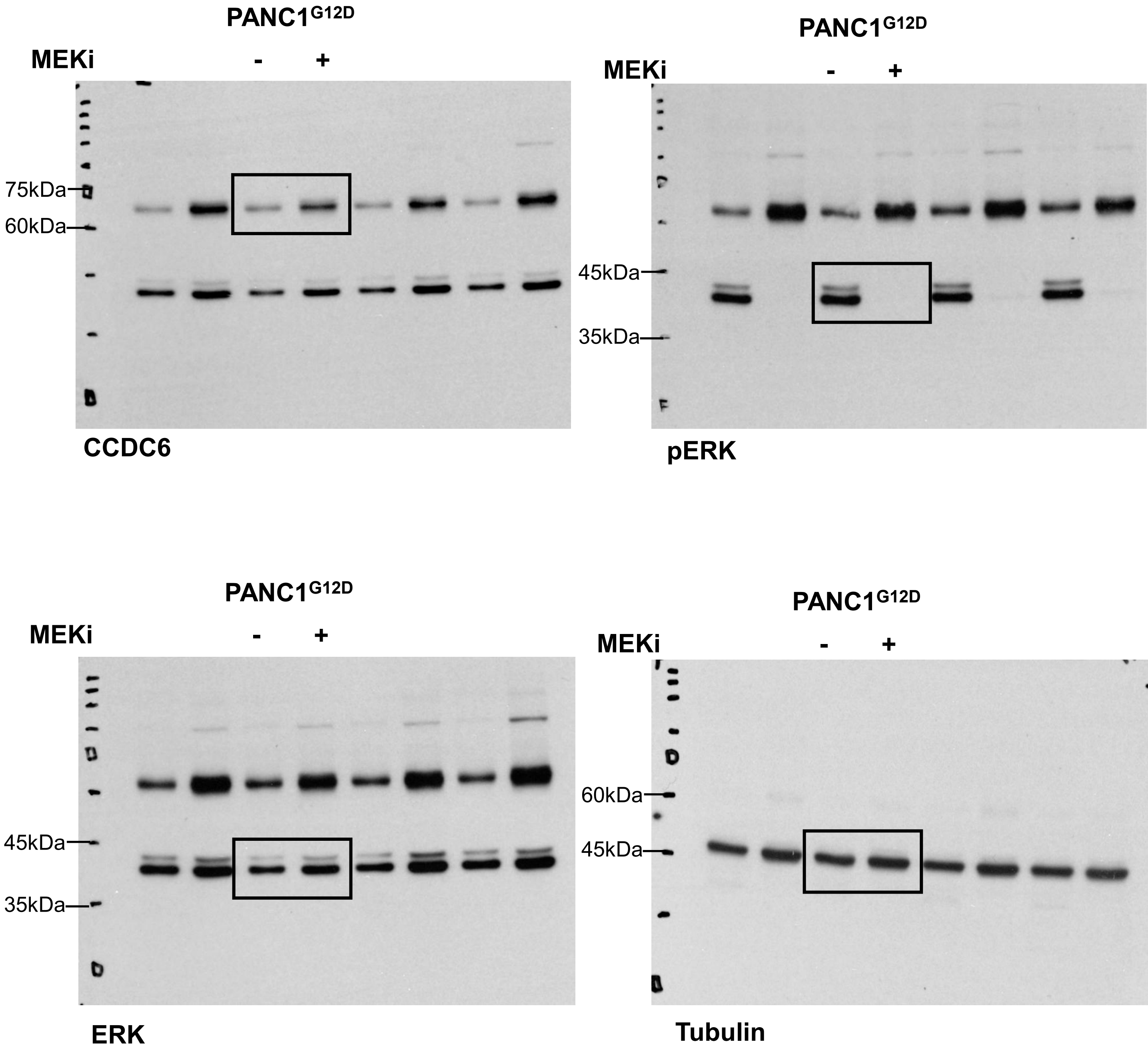

PANC1G12D
MEKi - +
75kDa
60kDa
CCDC6
PANC1G12D
MEKi - +
45kDa
35kDa
pERK
PANC1G12D
MEKi - +
45kDa
35kDa
ERK
PANC1G12D
MEKi - +
60kDa
45kDa
Tubulin

Original Western Blot Relative to Figure 1E

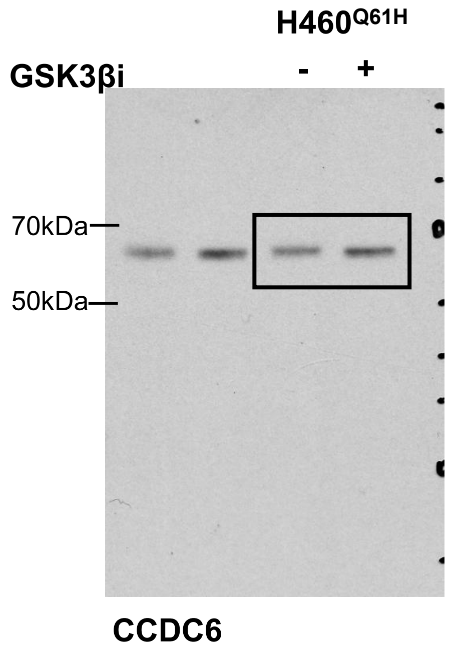


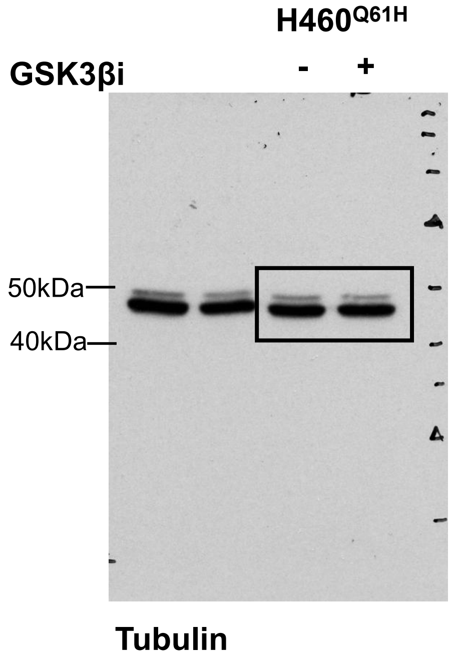


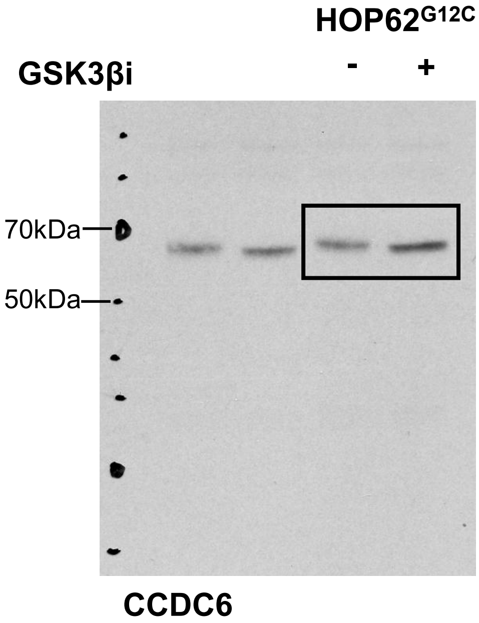


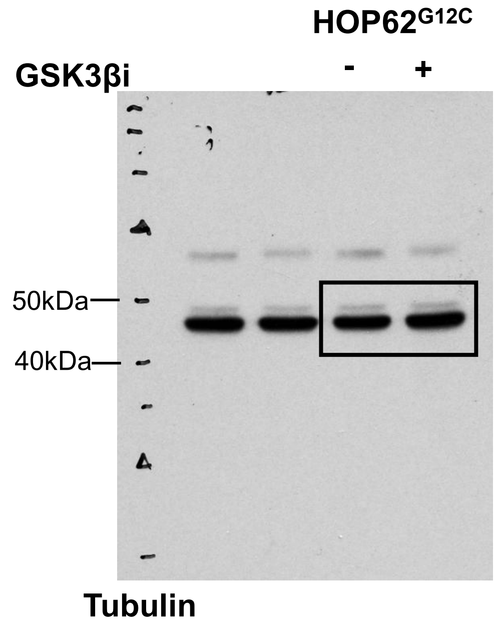


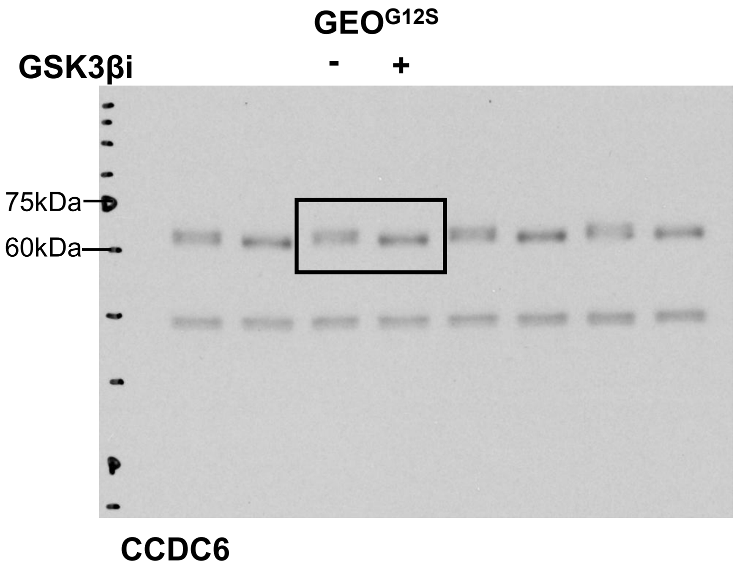


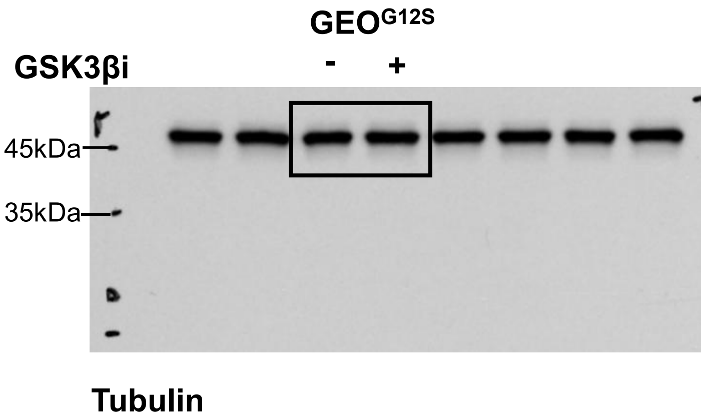

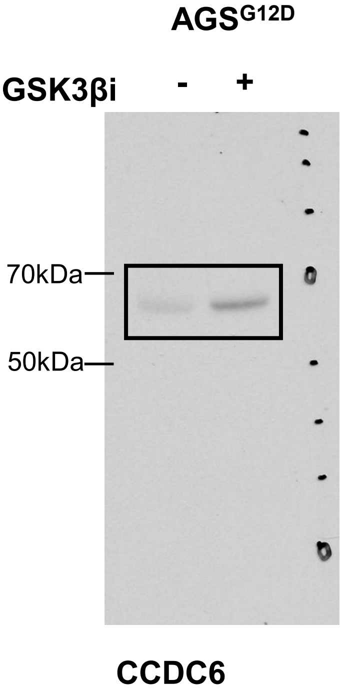
AGS$^{G12D}$
GSK3βi - +
70kDa
50kDa
CCDC6

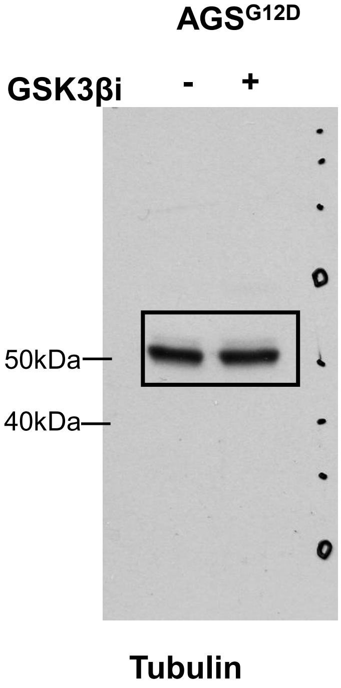
AGS$^{G12D}$
GSK3βi - +
50kDa
40kDa
Tubulin

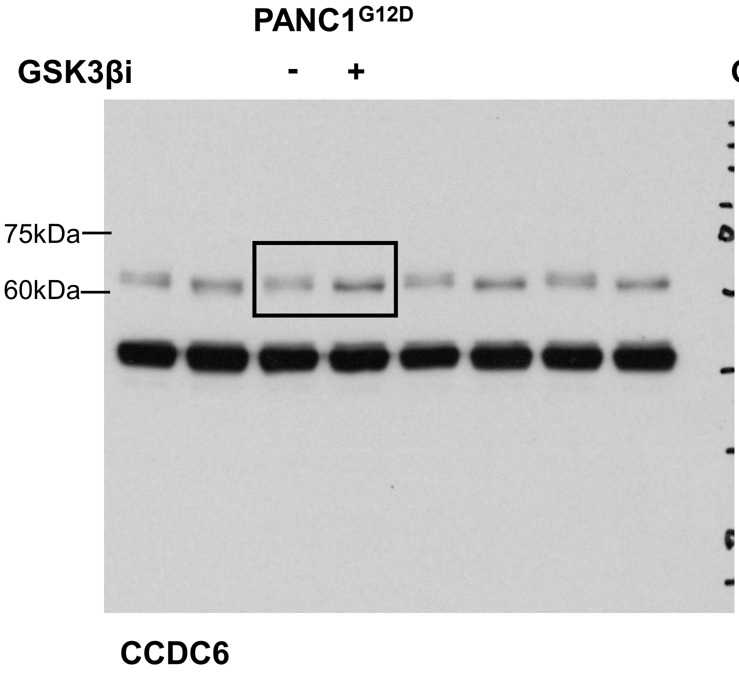
PANC1$^{G12D}$
GSK3βi - +
75kDa
60kDa
CCDC6

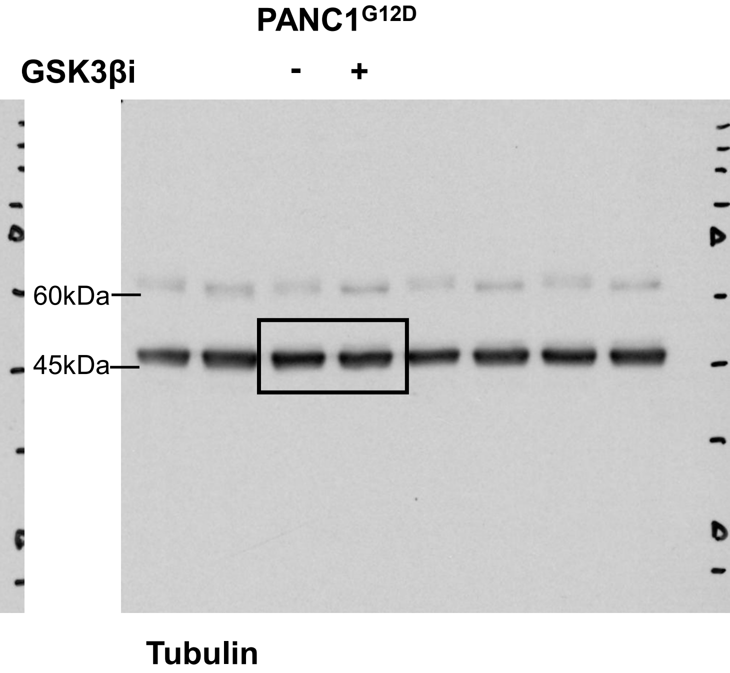
PANC1$^{G12D}$
GSK3βi - +
60kDa
45kDa
Tubulin

Original Western Blot Relative to Figure 1F

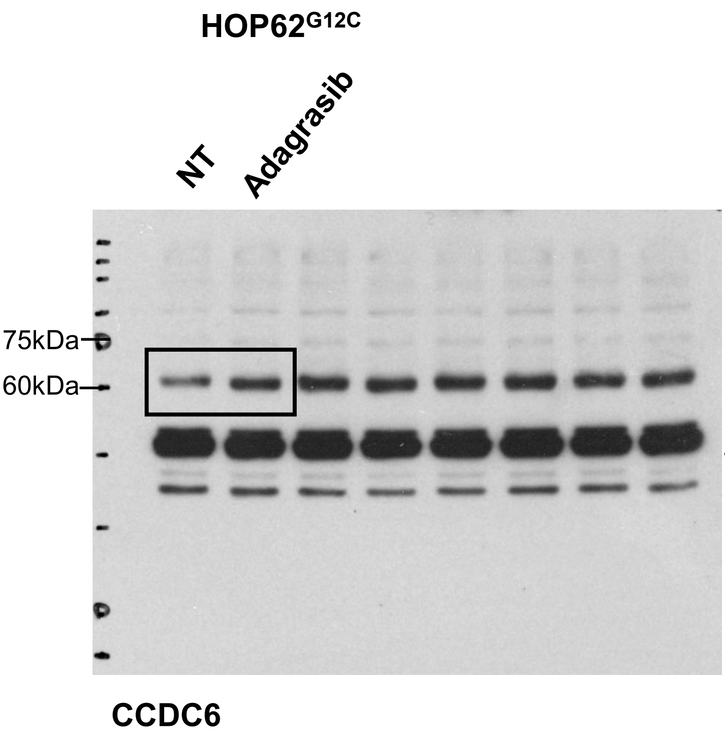


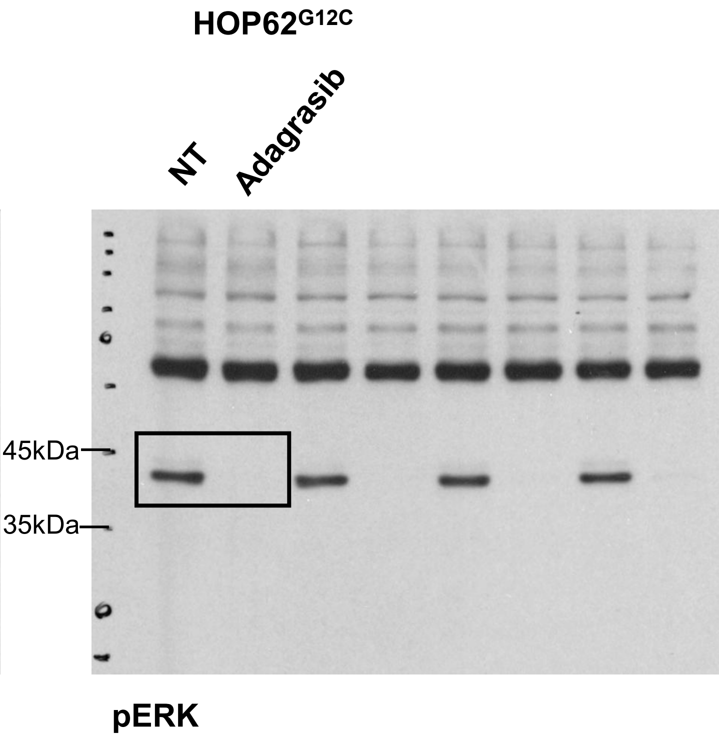


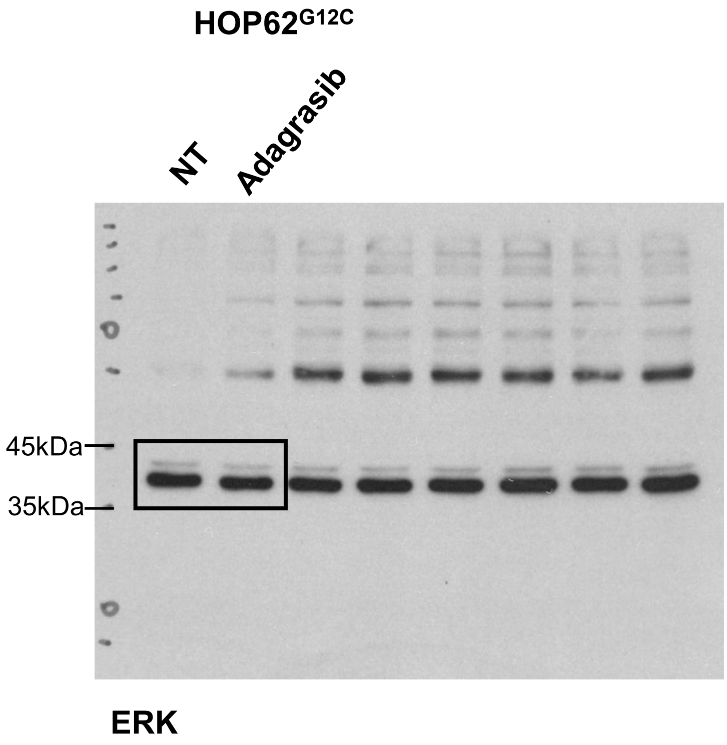


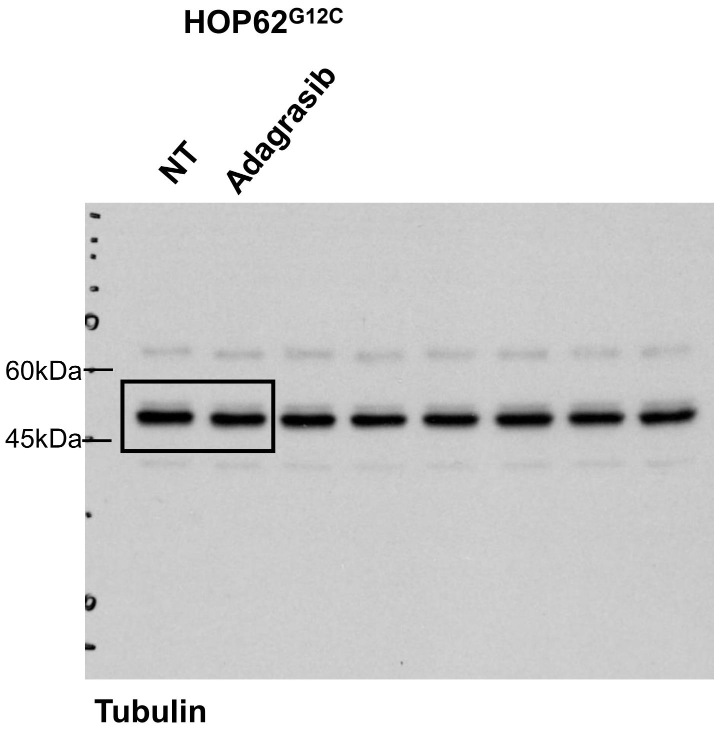

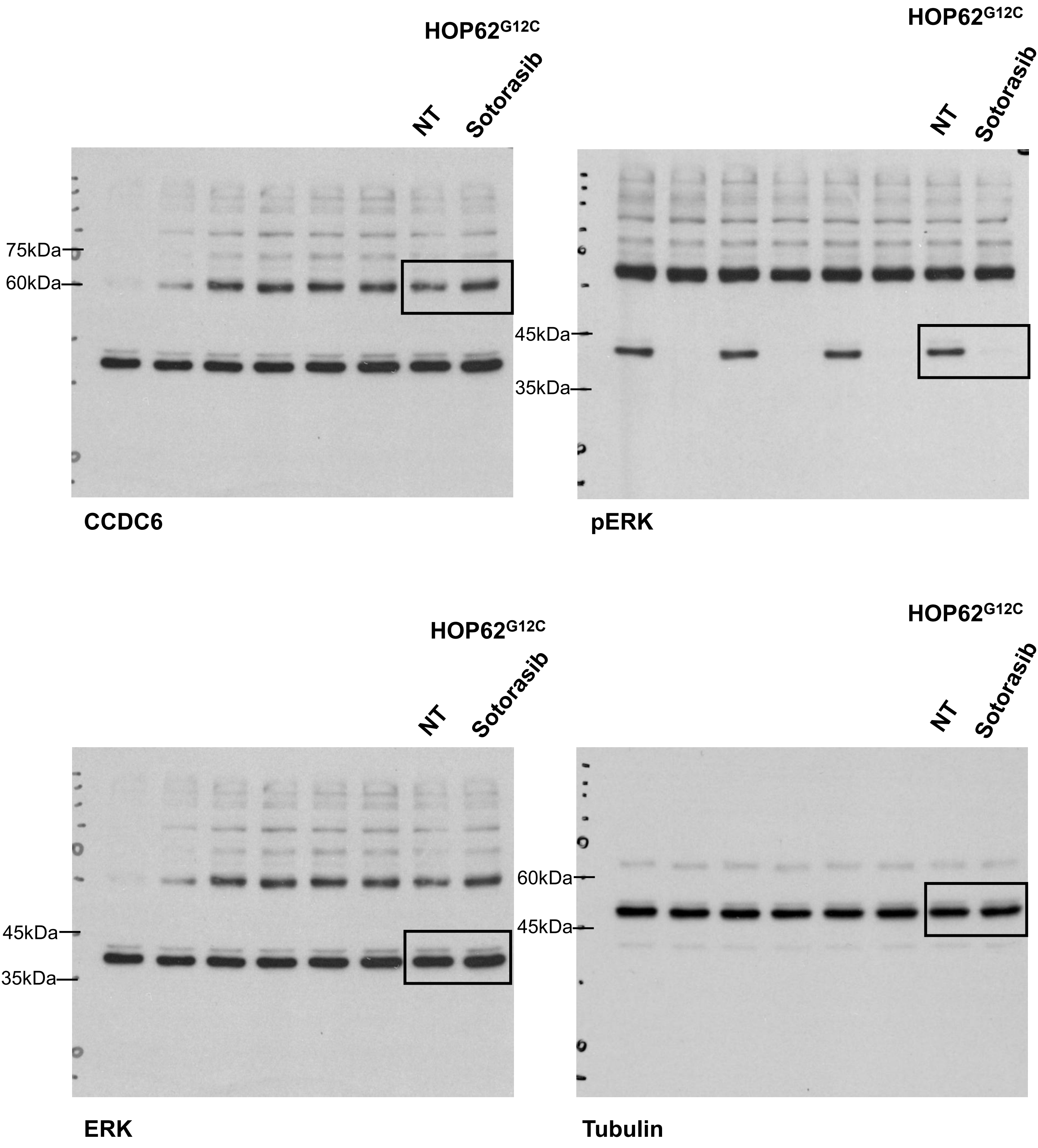
HOP62G12C
NT
Sotorasib
75kDa
60kDa
CCDC6
HOP62G12C
NT
Sotorasib
45kDa
35kDa
pERK
HOP62G12C
NT
Sotorasib
45kDa
35kDa
ERK
HOP62G12C
NT
Sotorasib
60kDa
45kDa
Tubulin

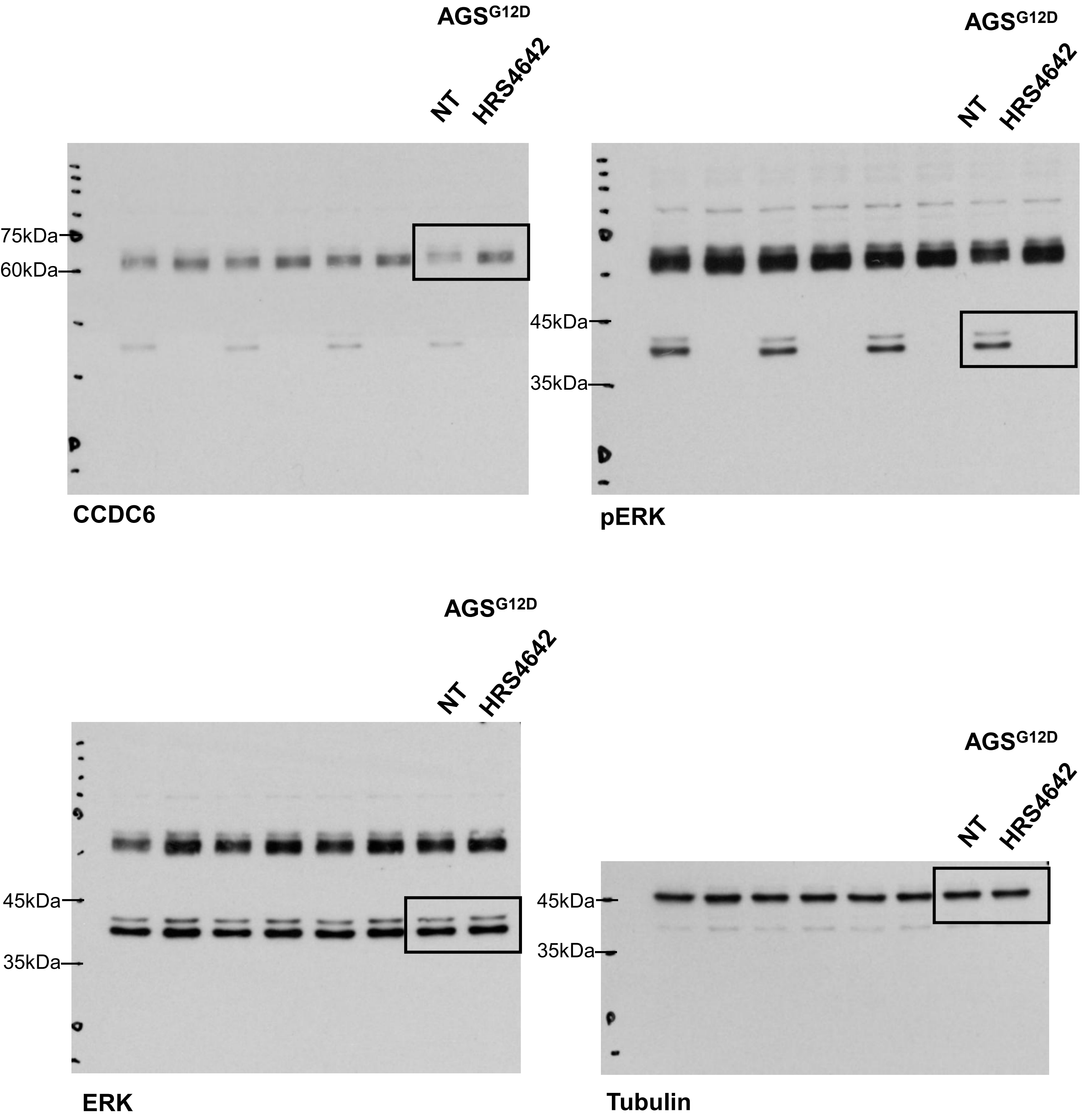
AGSG12D
NT
HRS4642
75kDa
60kDa
CCDC6
AGSG12D
NT
HRS4642
45kDa
35kDa
pERK
AGSG12D
NT
HRS4642
45kDa
35kDa
ERK
AGSG12D
NT
HRS4642
45kDa
35kDa
Tubulin

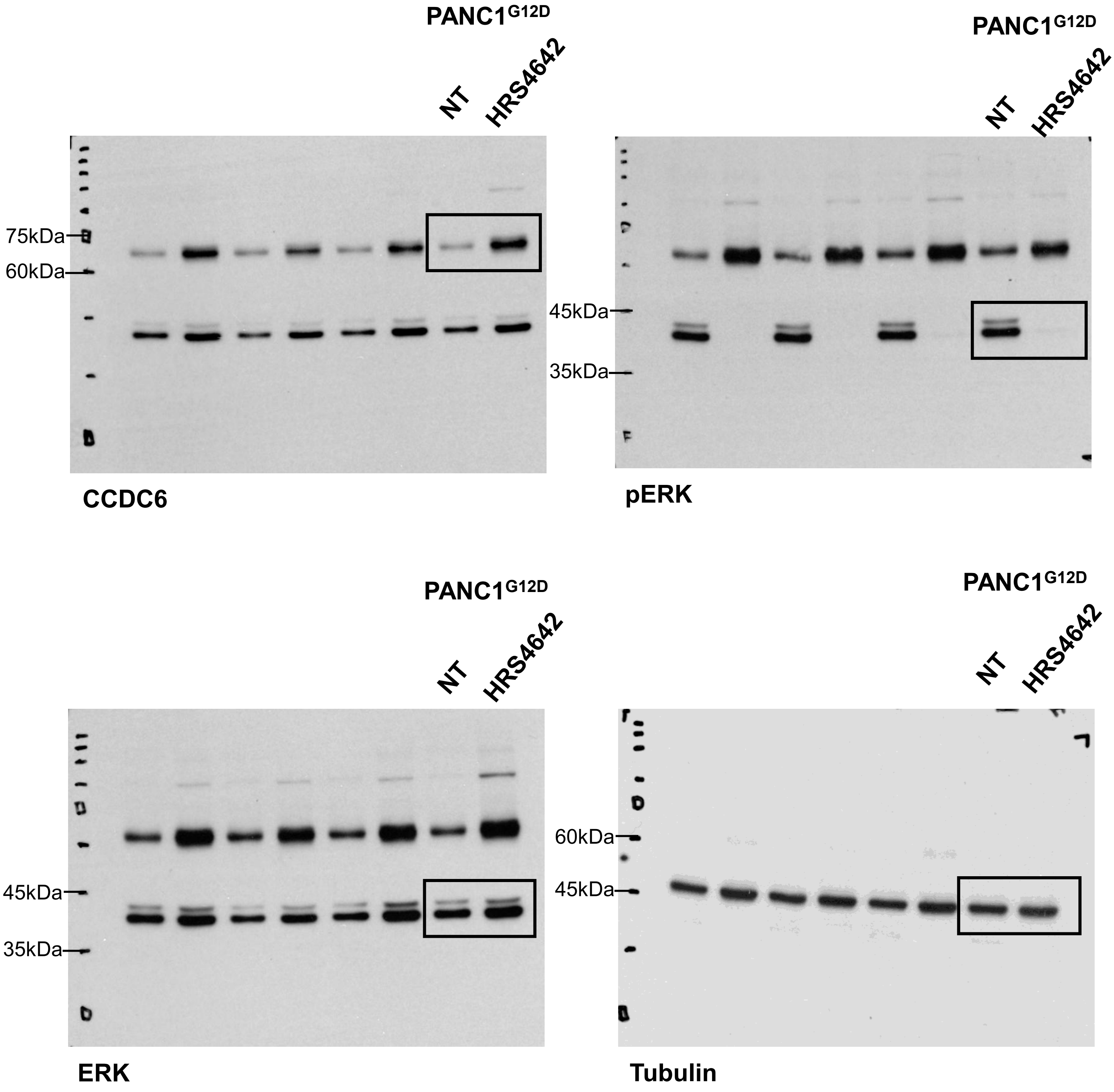

PANC1G12D
NT
HRS4642
75kDa
60kDa
CCDC6
PANC1G12D
NT
HRS4642
45kDa
35kDa
pERK
PANC1G12D
NT
HRS4642
45kDa
35kDa
ERK
PANC1G12D
NT
HRS4642
60kDa
45kDa
Tubulin

Original Western Blot Relative to Figure 1G

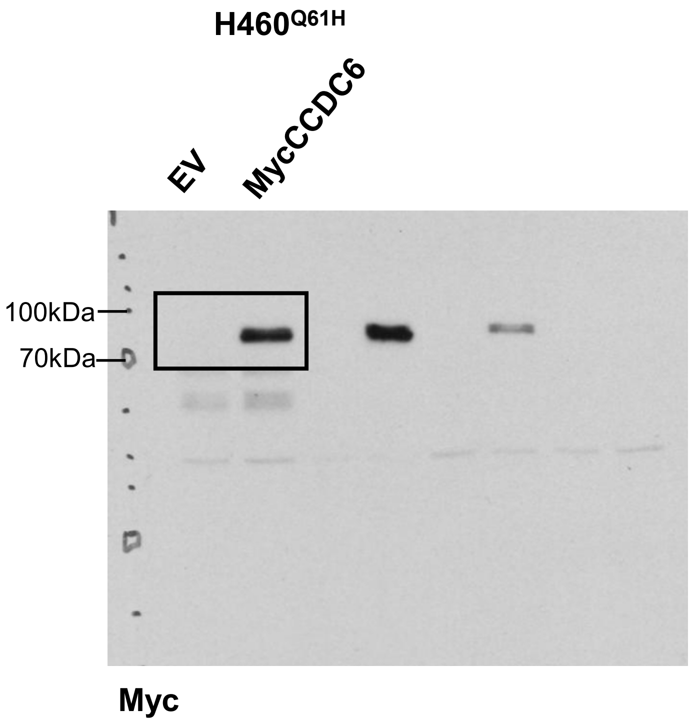


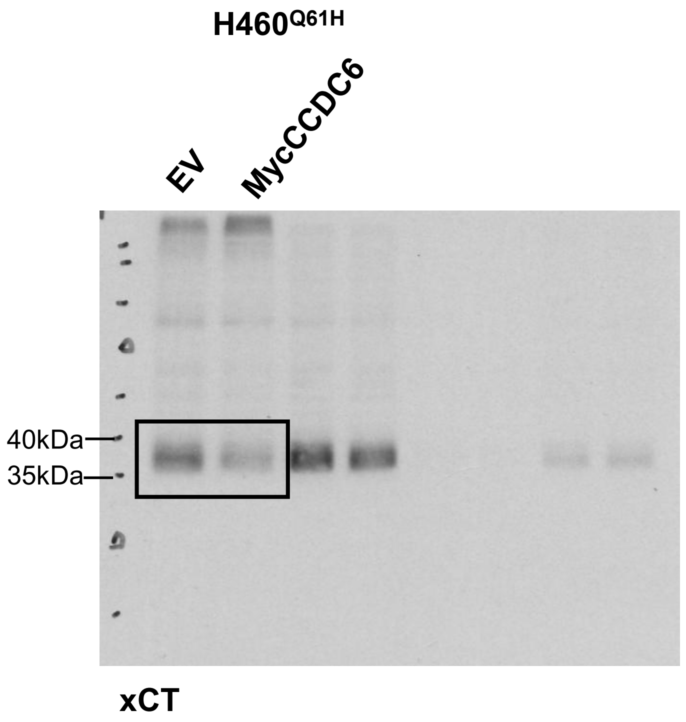


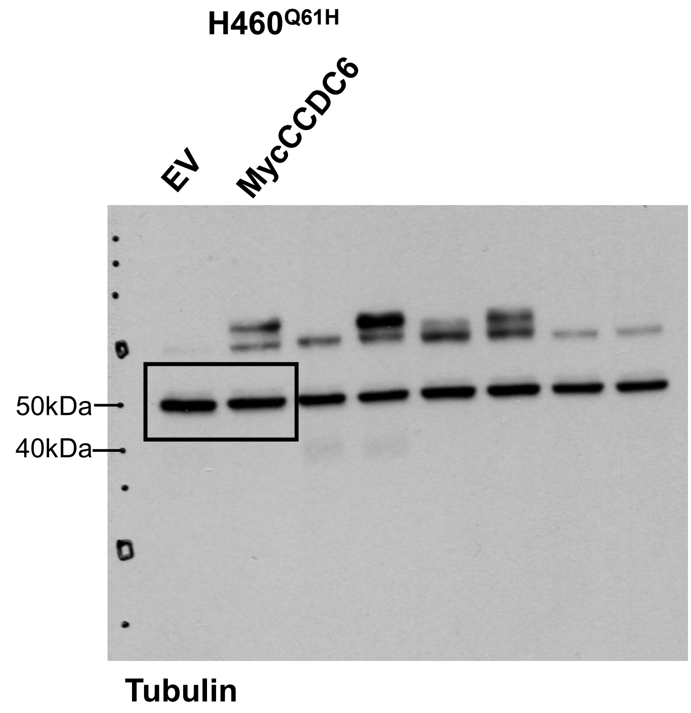

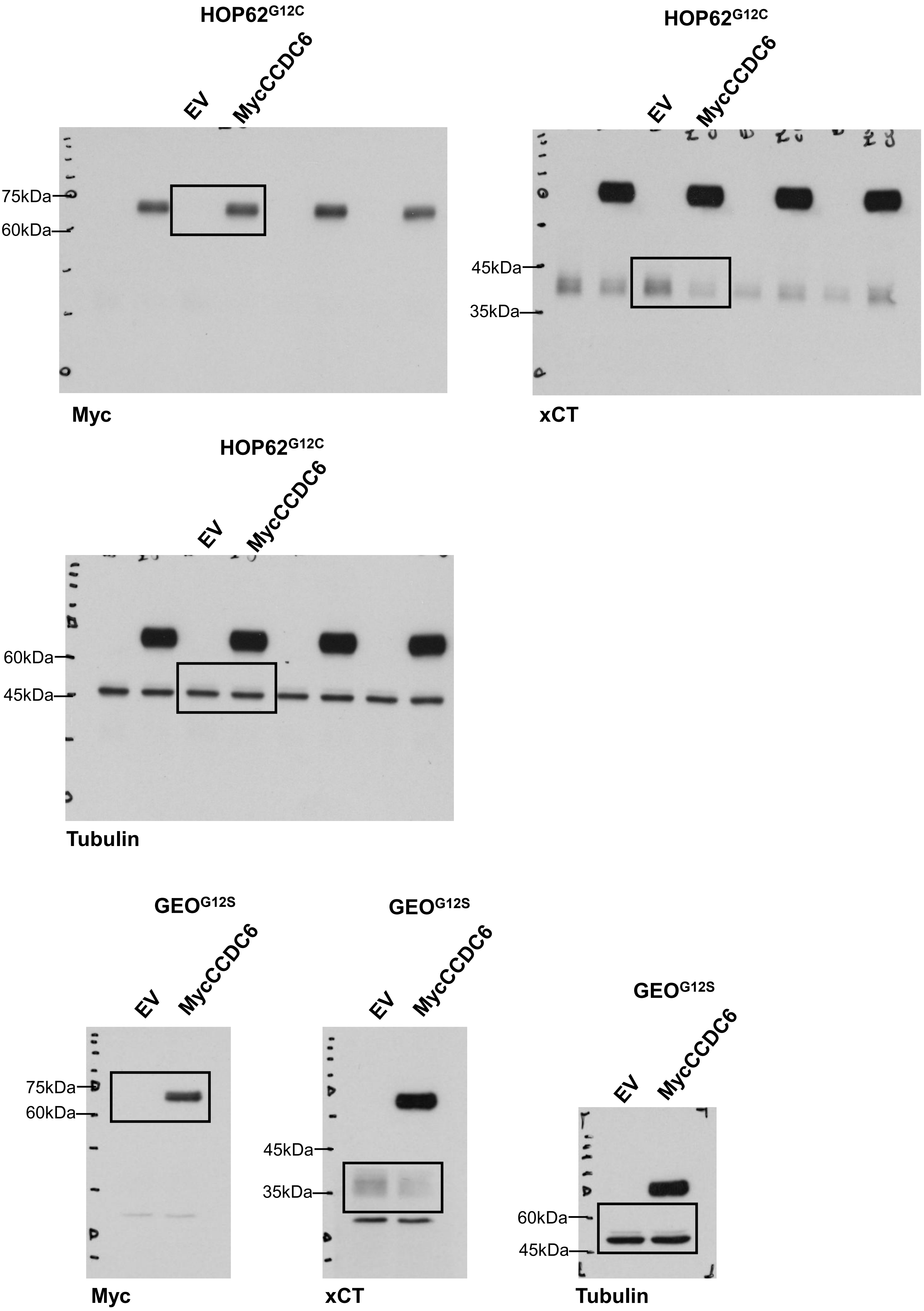

HOP62G12C
EV
MycCCDC6
75kDa
60kDa
Myc
HOP62G12C
EV
MycCCDC6
45kDa
35kDa
xCT
HOP62G12C
EV
MycCCDC6
60kDa
45kDa
Tubulin
GEOG12S
EV
MycCCDC6
75kDa
60kDa
Myc
GEOG12S
EV
MycCCDC6
45kDa
35kDa
xCT
GEOG12S
EV
MycCCDC6
60kDa
45kDa
Tubulin

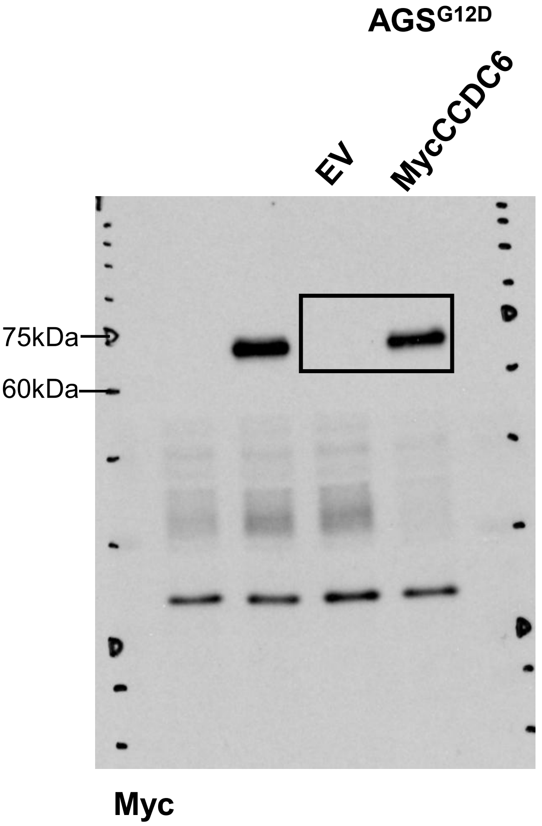
AGS$^{G12D}$
EV
MycCCDC6
75kDa
60kDa
Myc

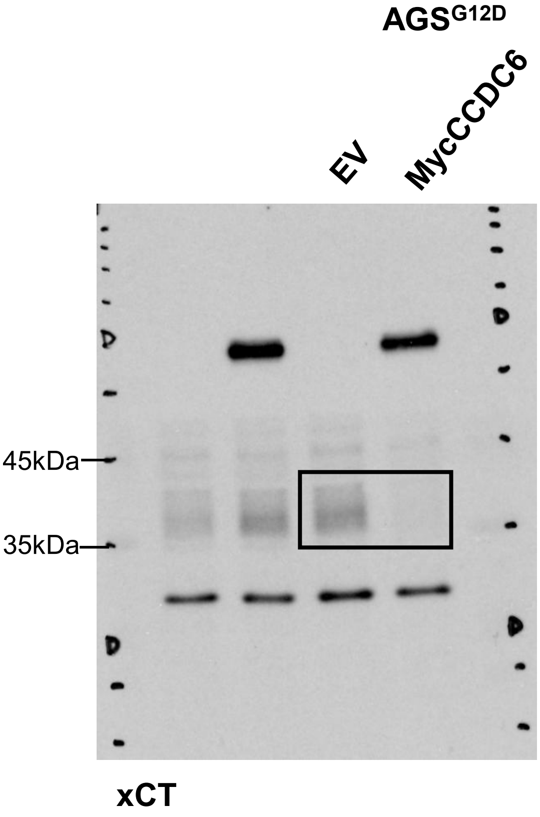
AGS$^{G12D}$
EV
MycCCDC6
45kDa
35kDa
xCT

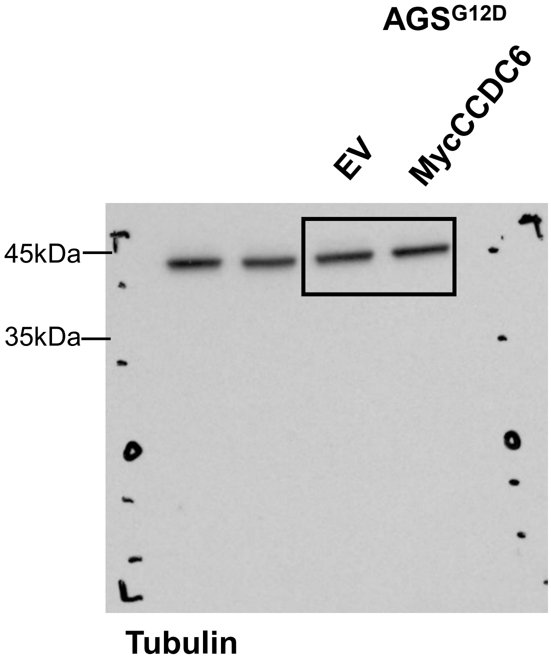
AGS$^{G12D}$
EV
MycCCDC6
45kDa
35kDa
Tubulin

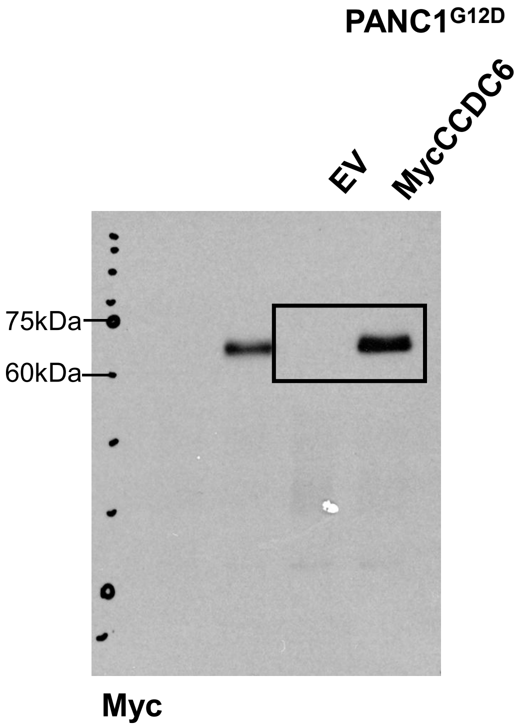
PANC1G12D
EV
MycCCDC6
75kDa
60kDa
Myc

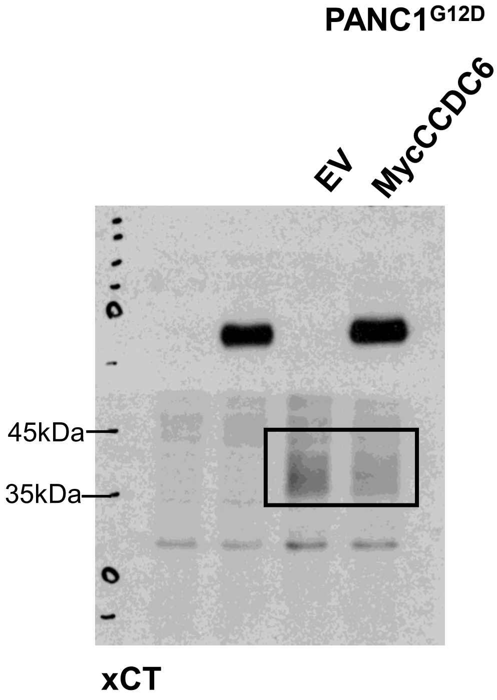
PANC1G12D
EV
MycCCDC6
45kDa
35kDa
xCT

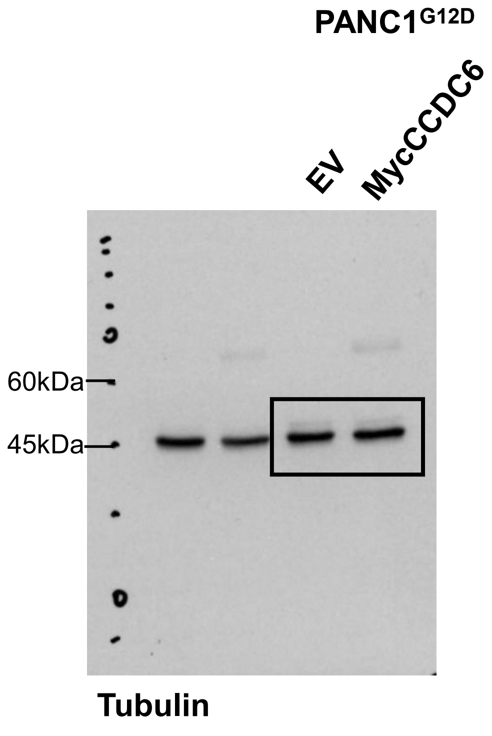
PANC1G12D
EV
MycCCDC6
60kDa
45kDa
Tubulin

Original Western Blot Relative to Figure 2A

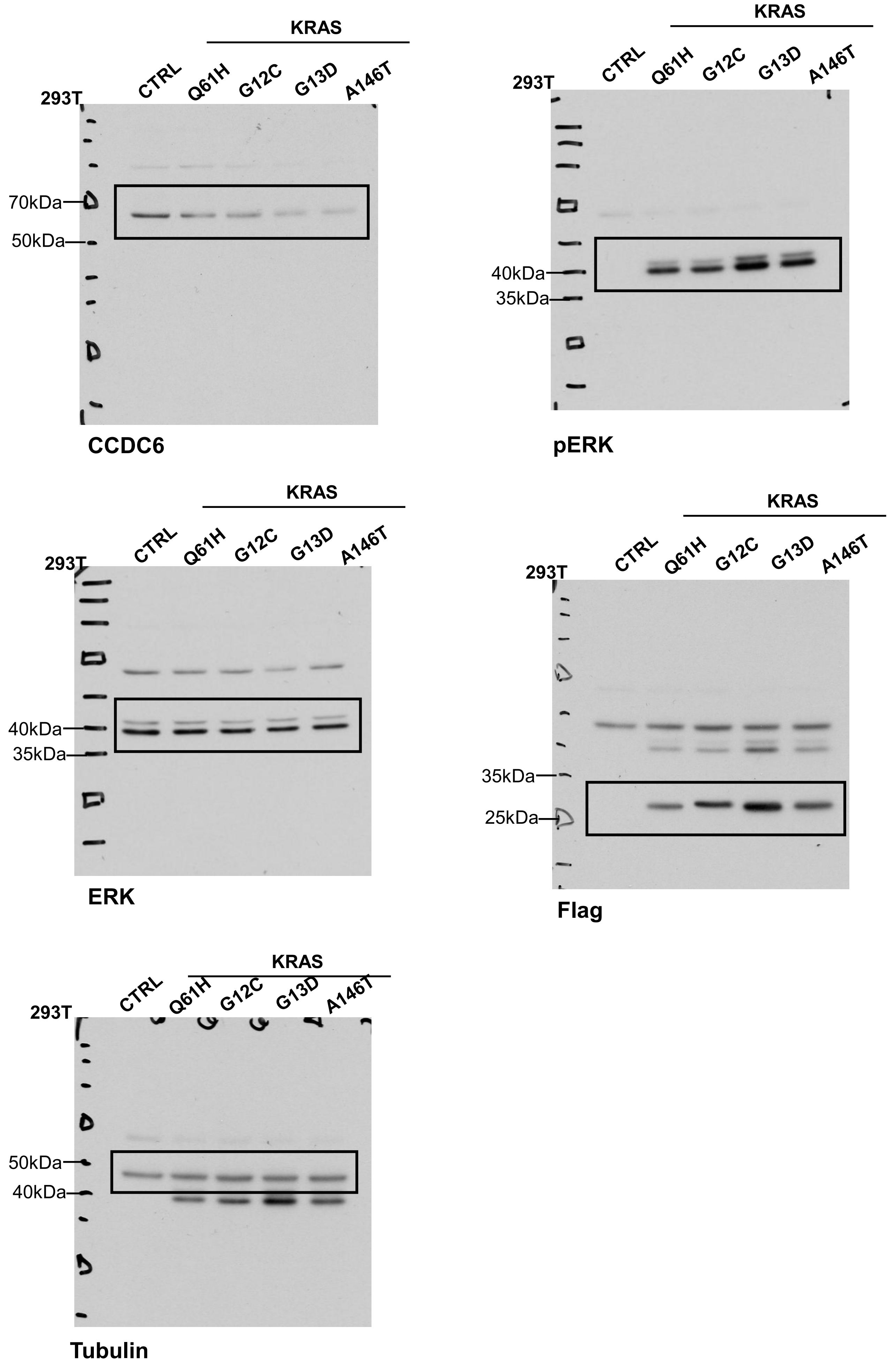

Original Western Blot Relative to Figure 2C

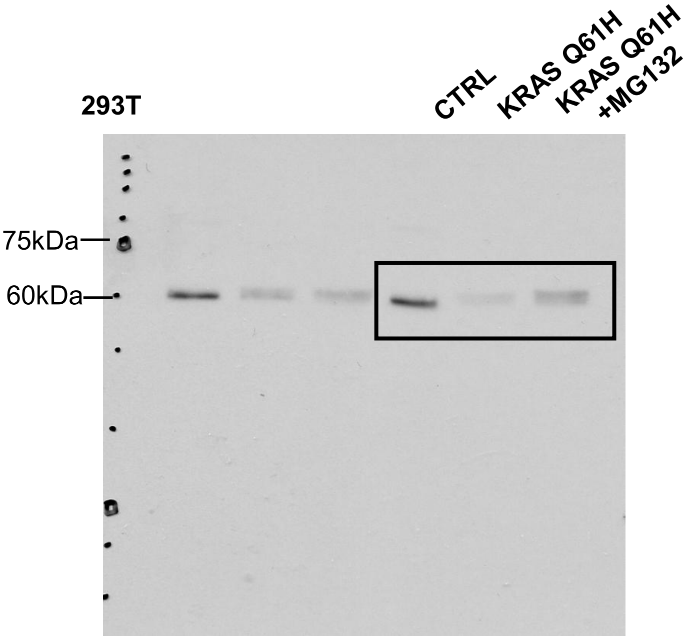


CCDC6

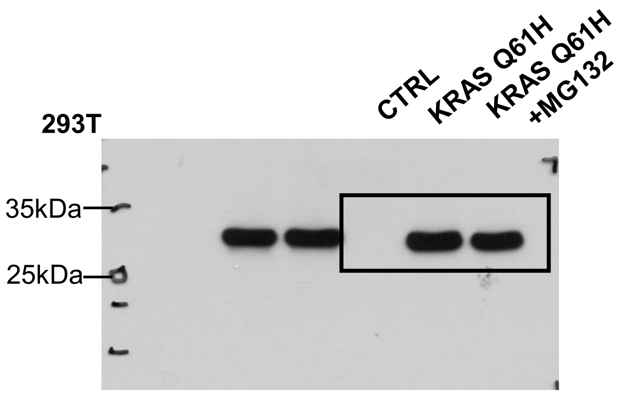


Flag

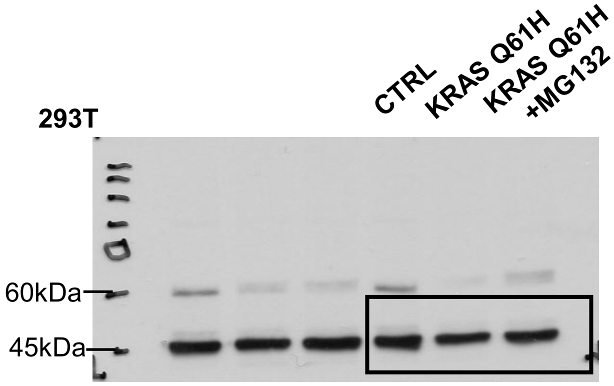


Tubulin

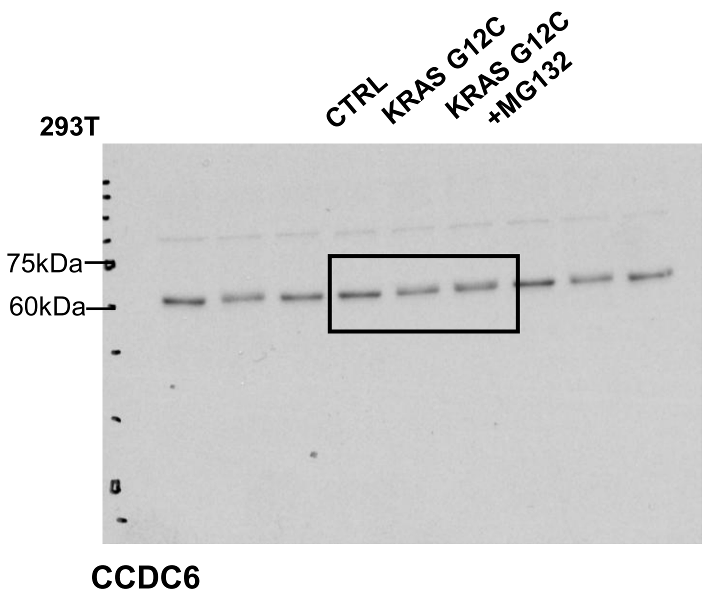
293T
CTRL
KRAS G12C
KRAS G12C +MG132
75kDa
60kDa
CCDC6

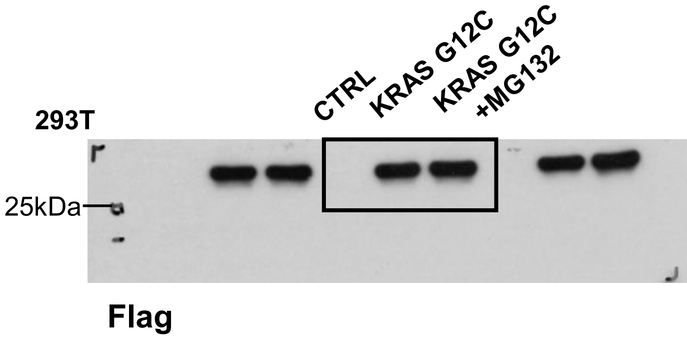
293T
CTRL
KRAS G12C
KRAS G12C +MG132
25kDa
Flag

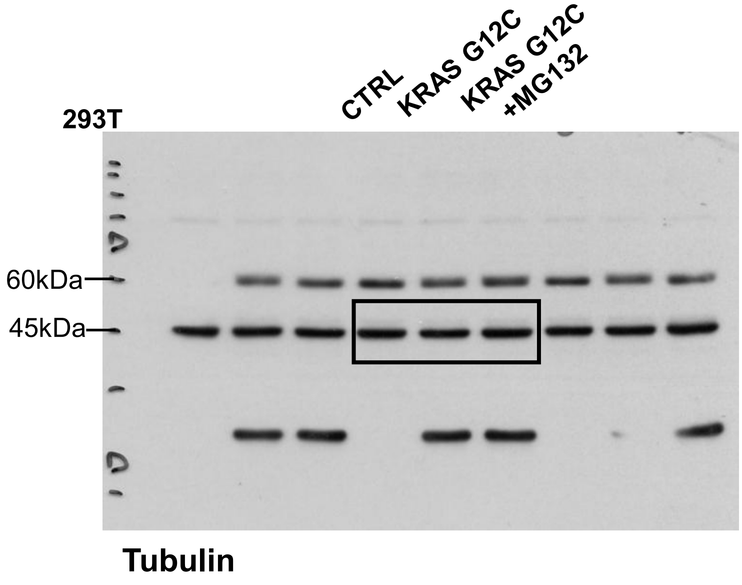
293T
CTRL
KRAS G12C
KRAS G12C +MG132
60kDa
45kDa
Tubulin

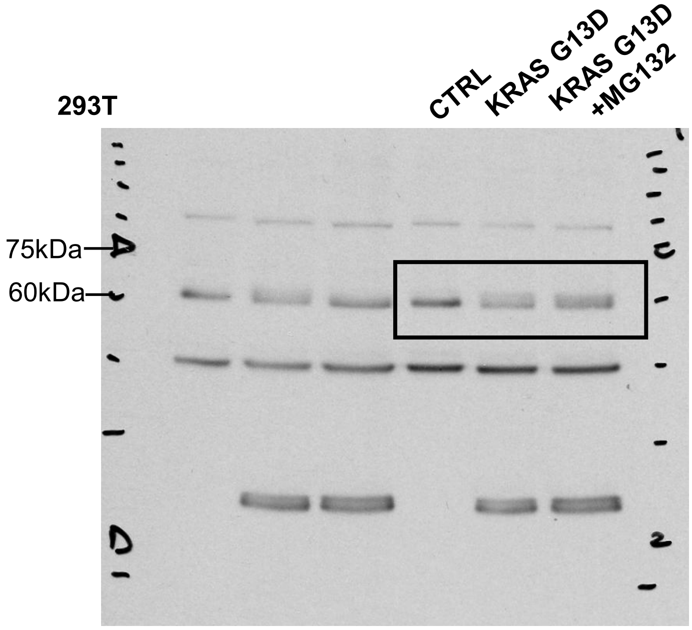


CCDC6

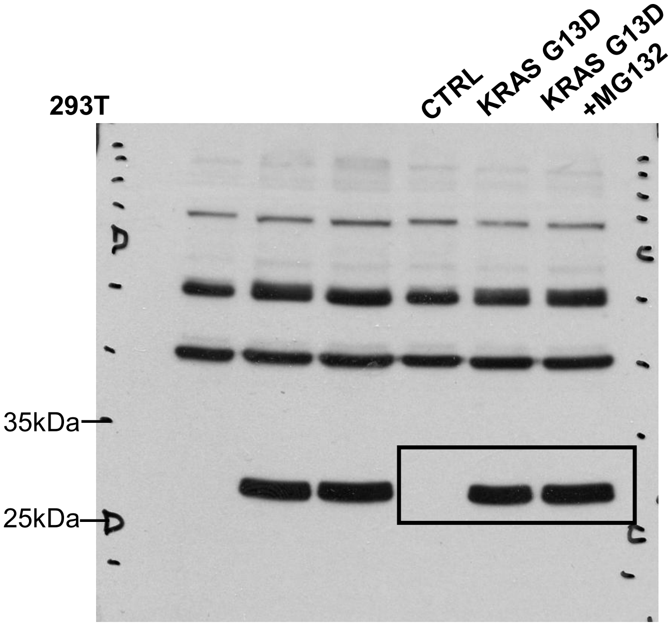


Flag

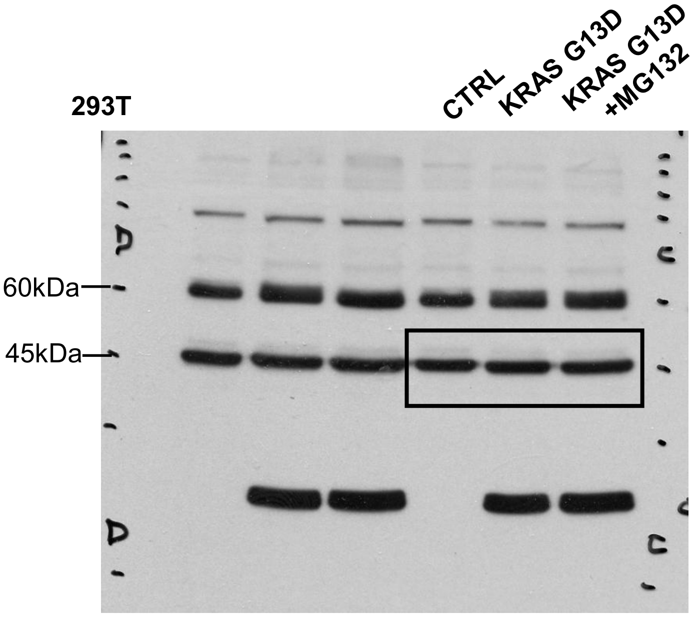


Tubulin

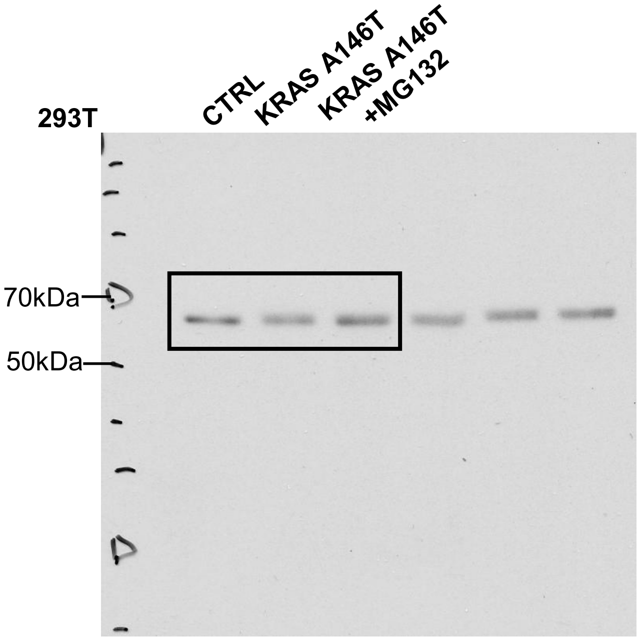


CCDC6

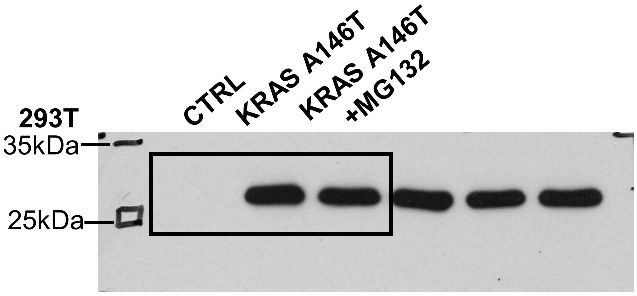


Flag

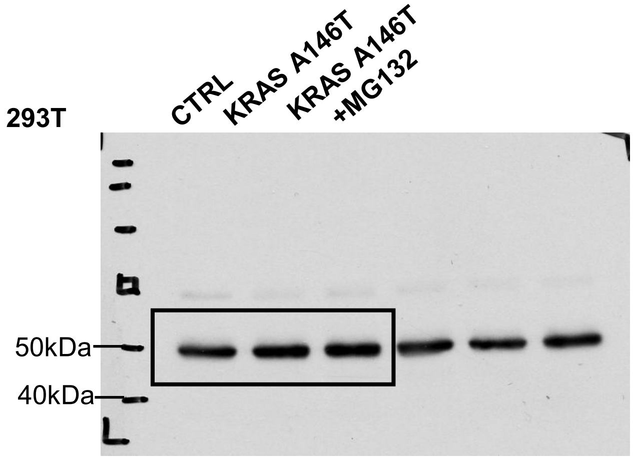


Tubulin

Original Western Blot Relative to Figure 2D

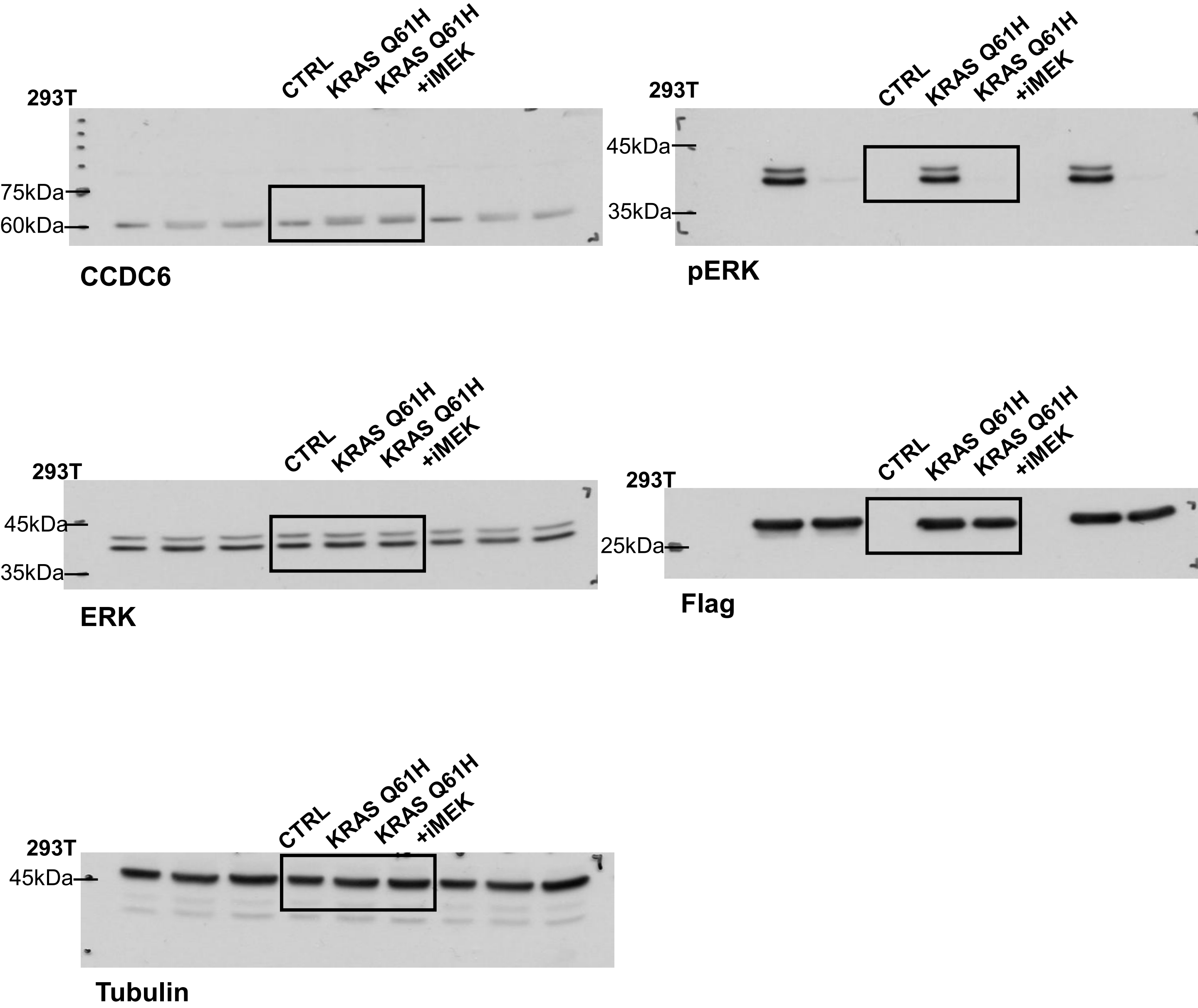

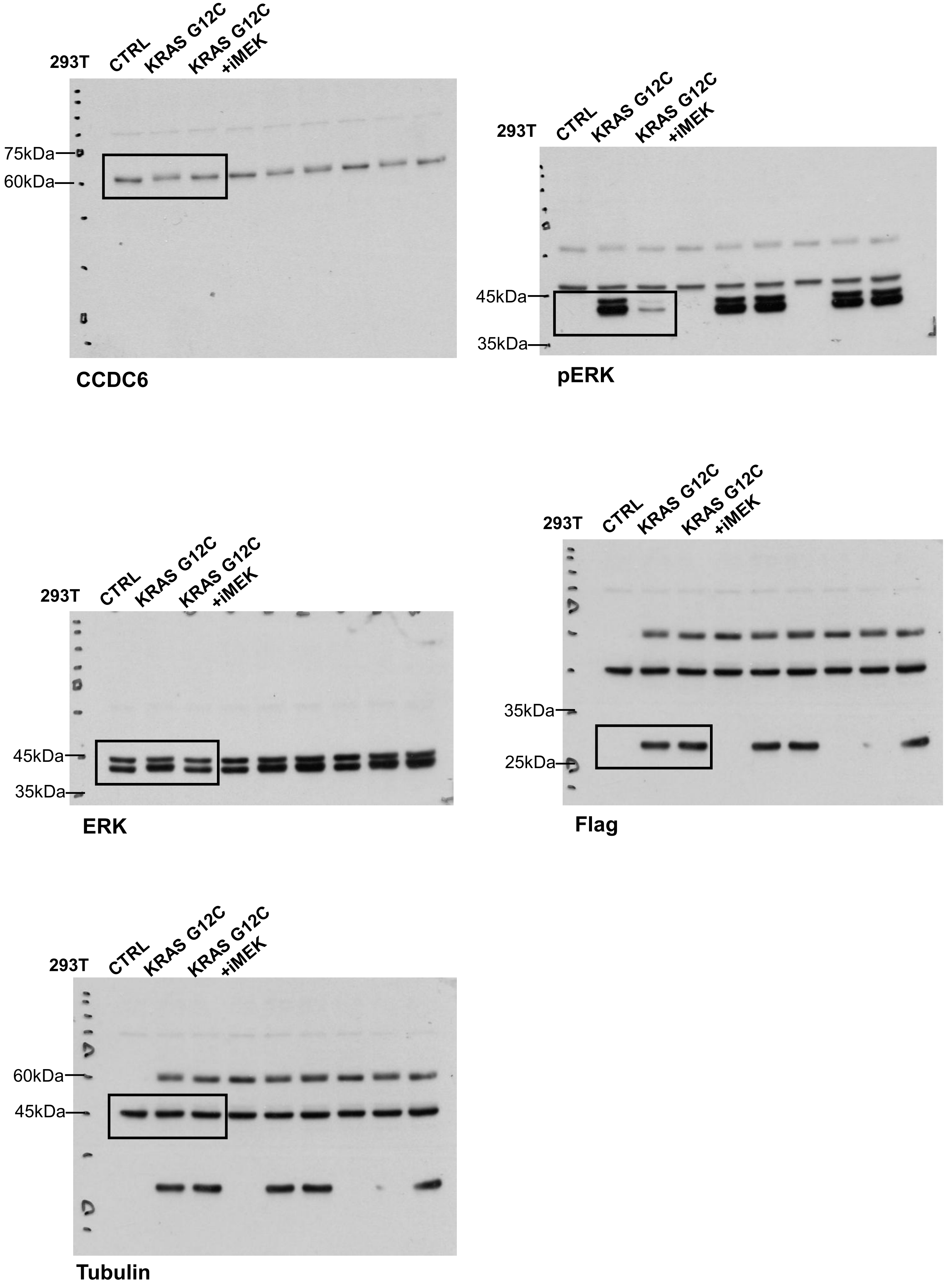
293T
CTRL
KRAS G12C
KRAS G12C +iMEK
75kDa
60kDa
CCDC6
293T
CTRL
KRAS G12C
KRAS G12C +iMEK
45kDa
35kDa
pERK
293T
CTRL
KRAS G12C
KRAS G12C +iMEK
45kDa
35kDa
ERK
293T
CTRL
KRAS G12C
KRAS G12C +iMEK
35kDa
25kDa
Flag
293T
CTRL
KRAS G12C
KRAS G12C +iMEK
60kDa
45kDa
Tubulin

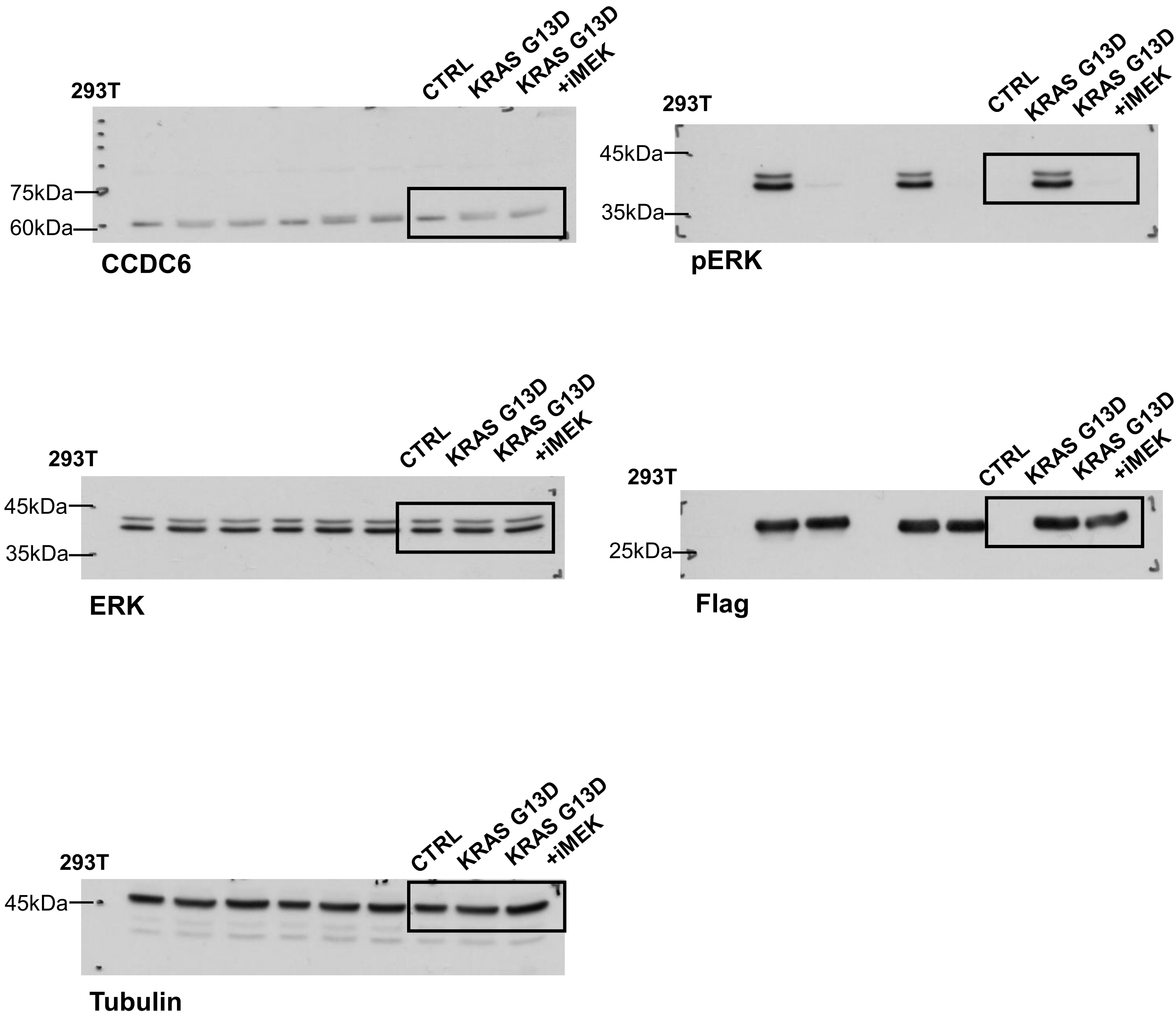
293T
CTRL
KRAS G13D
KRAS G13D +iMEK
75kDa
60kDa
CCDC6
293T
CTRL
KRAS G13D
KRAS G13D +iMEK
45kDa
35kDa
pERK
293T
CTRL
KRAS G13D
KRAS G13D +iMEK
45kDa
35kDa
ERK
293T
CTRL
KRAS G13D
KRAS G13D +iMEK
25kDa
Flag
293T
CTRL
KRAS G13D
KRAS G13D +iMEK
45kDa
Tubulin

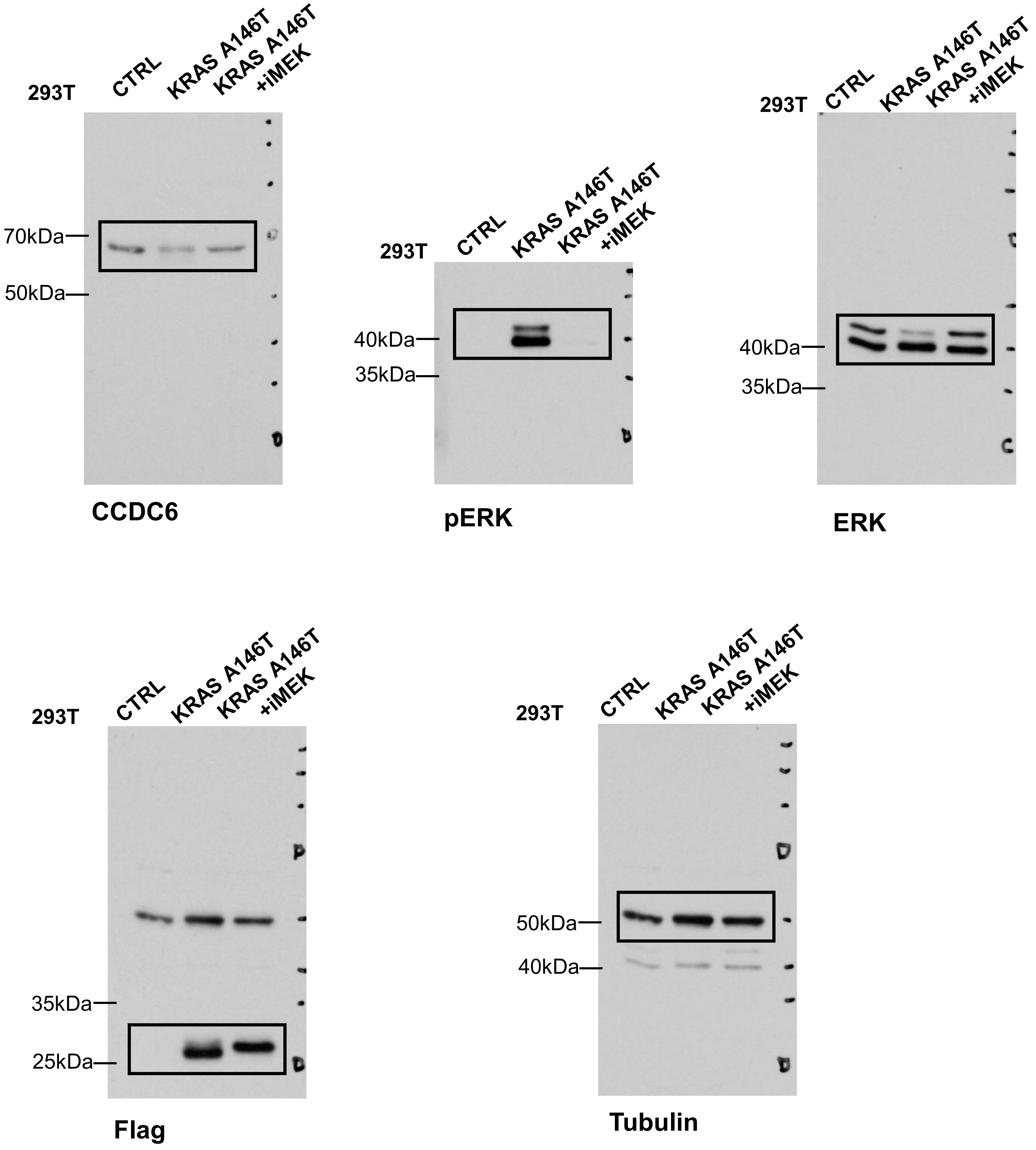
293T
CTRL
KRAS A146T
KRAS A146T +iMEK
70kDa
50kDa
CCDC6
293T
CTRL
KRAS A146T
KRAS A146T +iMEK
40kDa
35kDa
pERK
293T
CTRL
KRAS A146T
KRAS A146T +iMEK
40kDa
35kDa
ERK
293T
CTRL
KRAS A146T
KRAS A146T +iMEK
35kDa
25kDa
Flag
293T
CTRL
KRAS A146T
KRAS A146T +iMEK
50kDa
40kDa
Tubulin

Original Western Blot Relative to Figure 2E

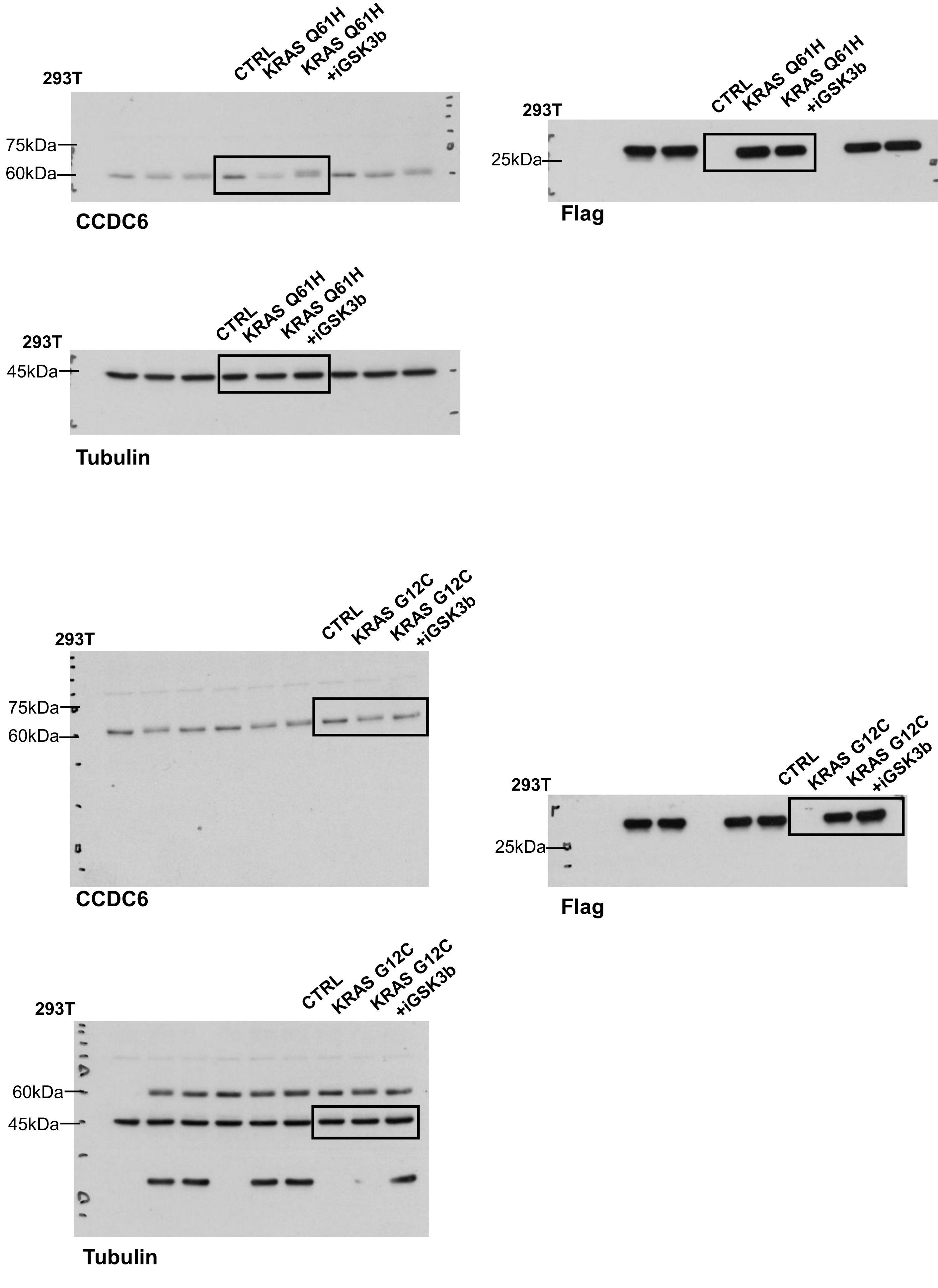

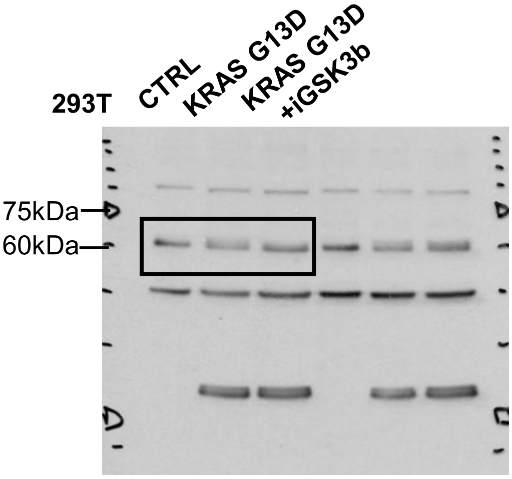


**CCDC6**

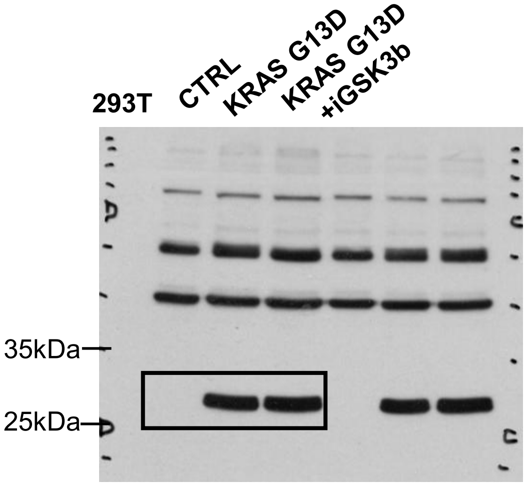


**Flag**

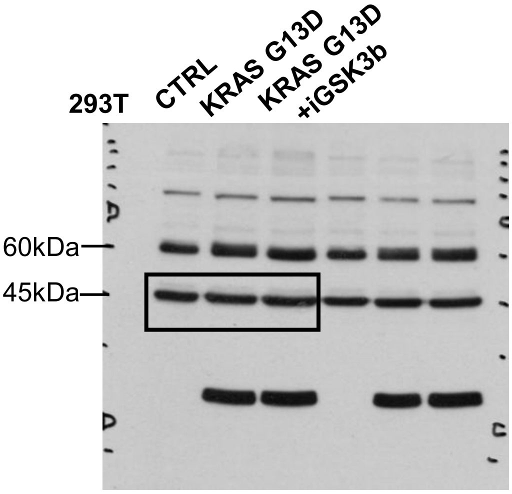


**Tubulin**

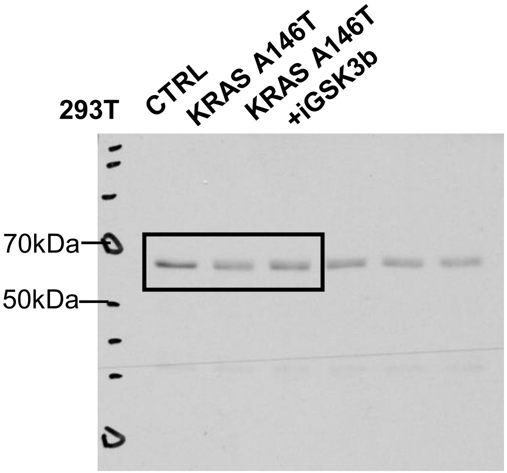


**CCDC6**

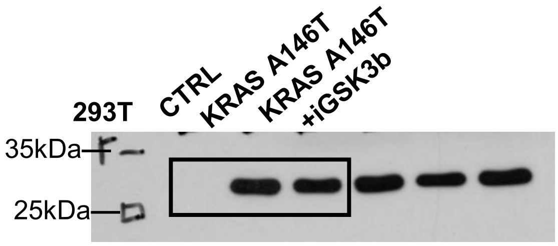


**Flag**

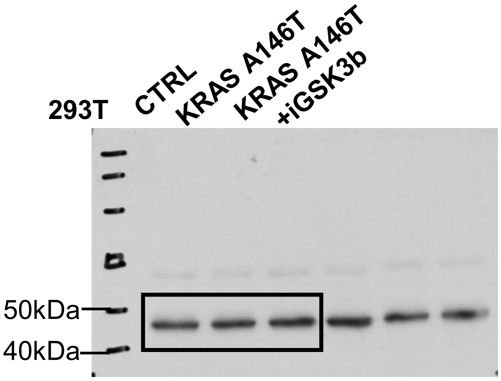


**Tubulin**

### Original Western Blot Relative to Figure 2F

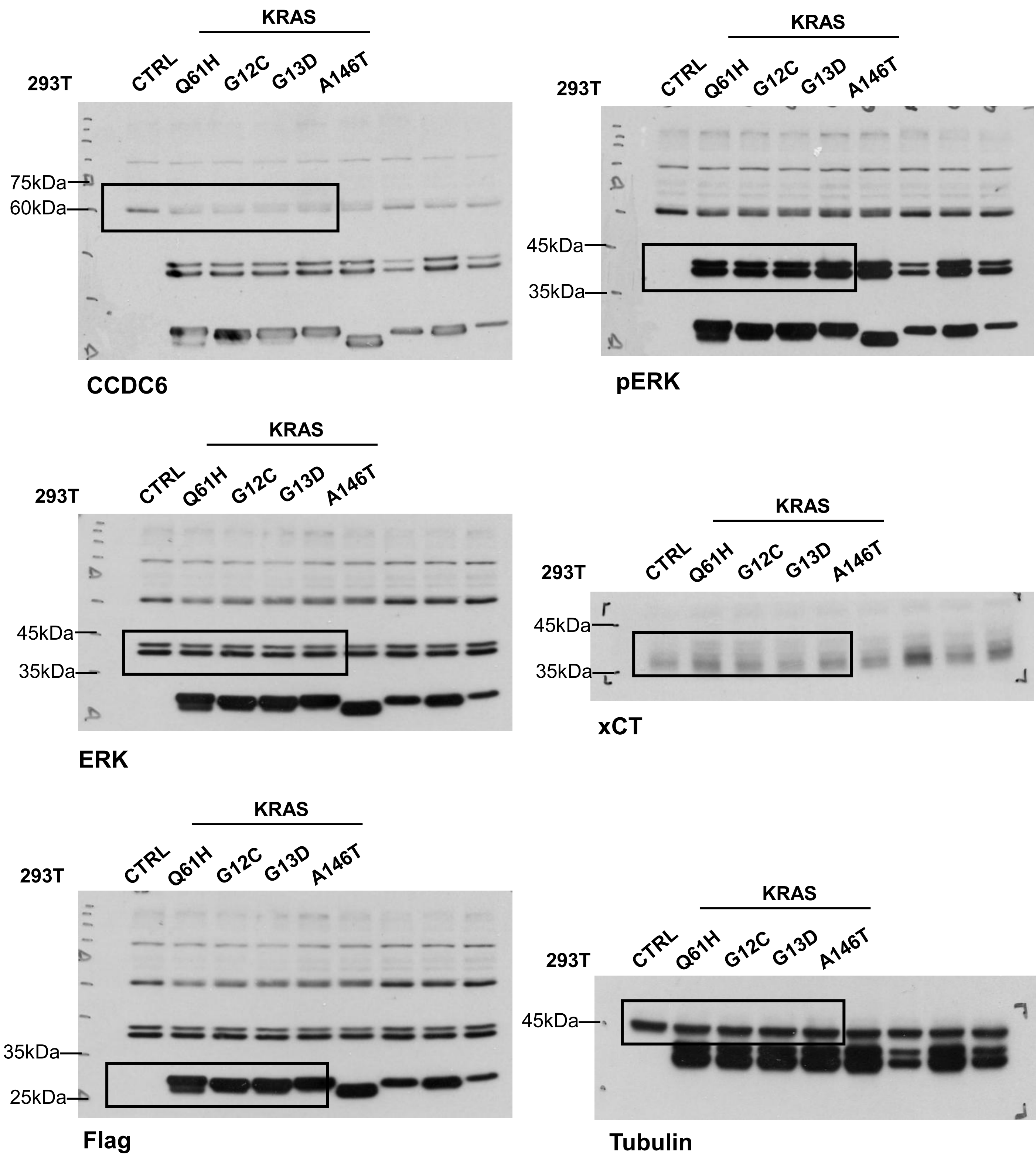

Original Western Blot Relative to Figure 2H

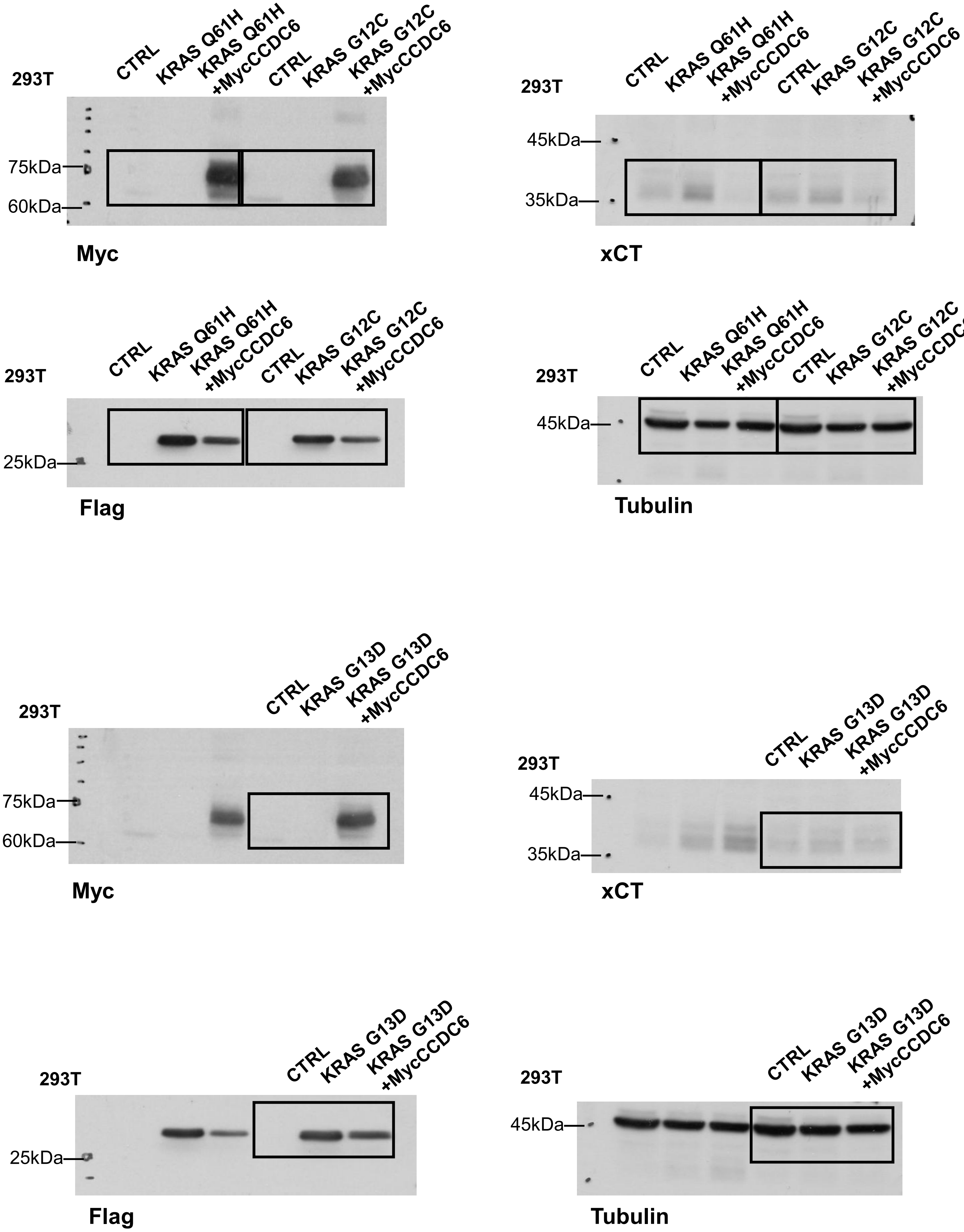

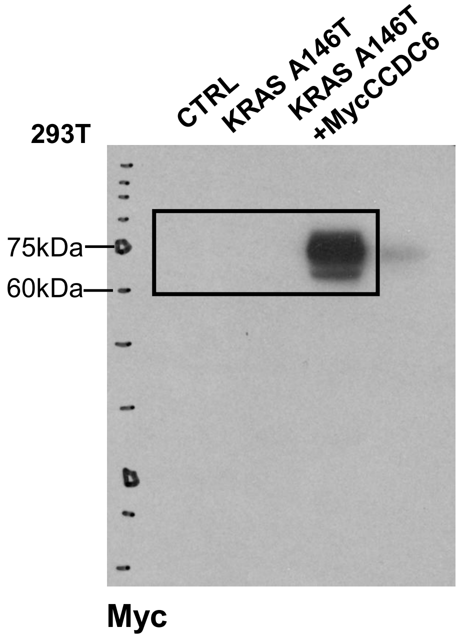

293T
CTRL
KRAS A146T
KRAS A146T +MycCCDC6
75kDa
60kDa
Myc


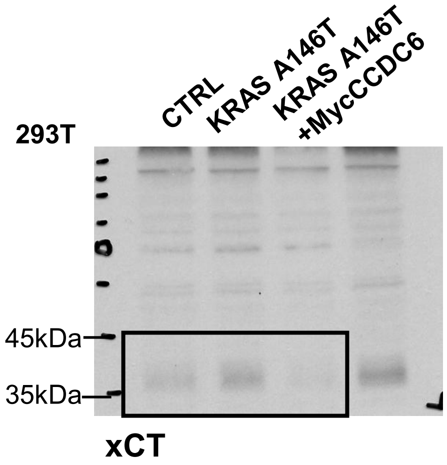

293T
CTRL
KRAS A146T
KRAS A146T +MycCCDC6
45kDa
35kDa
xCT


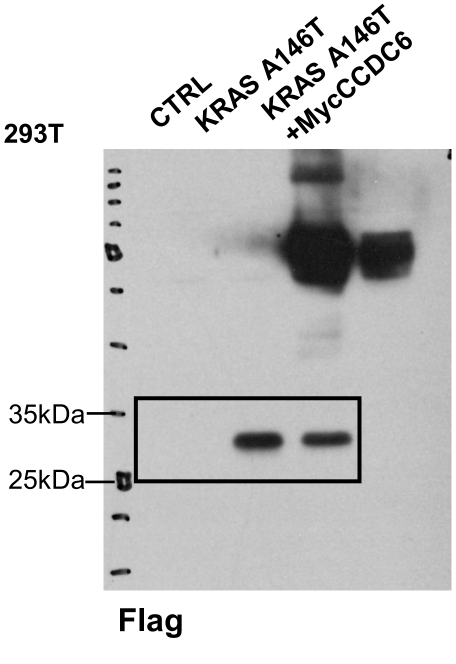

293T
CTRL
KRAS A146T
KRAS A146T +MycCCDC6
35kDa
25kDa
Flag


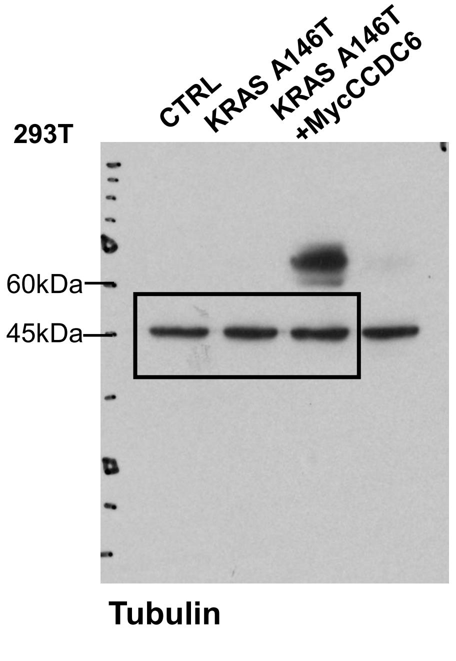

293T
CTRL
KRAS A146T
KRAS A146T +MycCCDC6
60kDa
45kDa
Tubulin

Original Western Blot Relative to Figure 3A

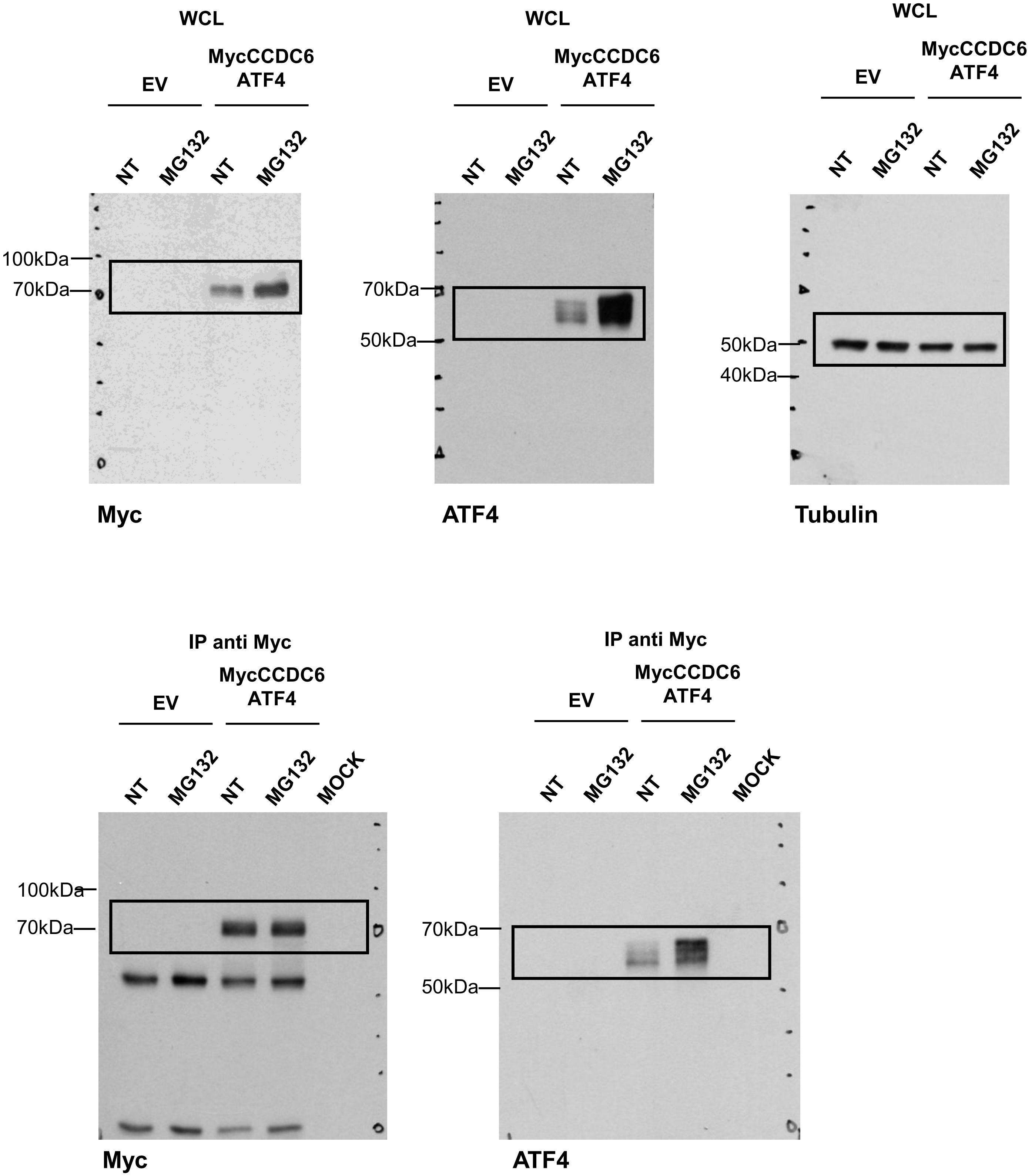

#### Original Western Blot Relative to Figure 3B

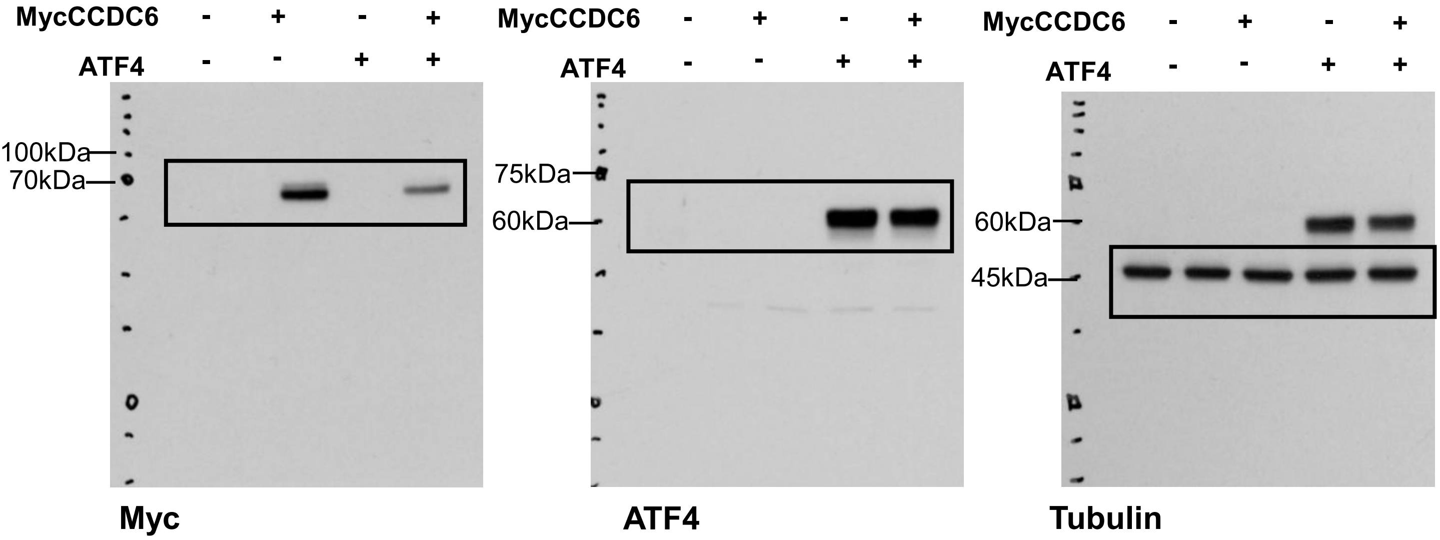


#### Original Western Blot Relative to Figure 3C

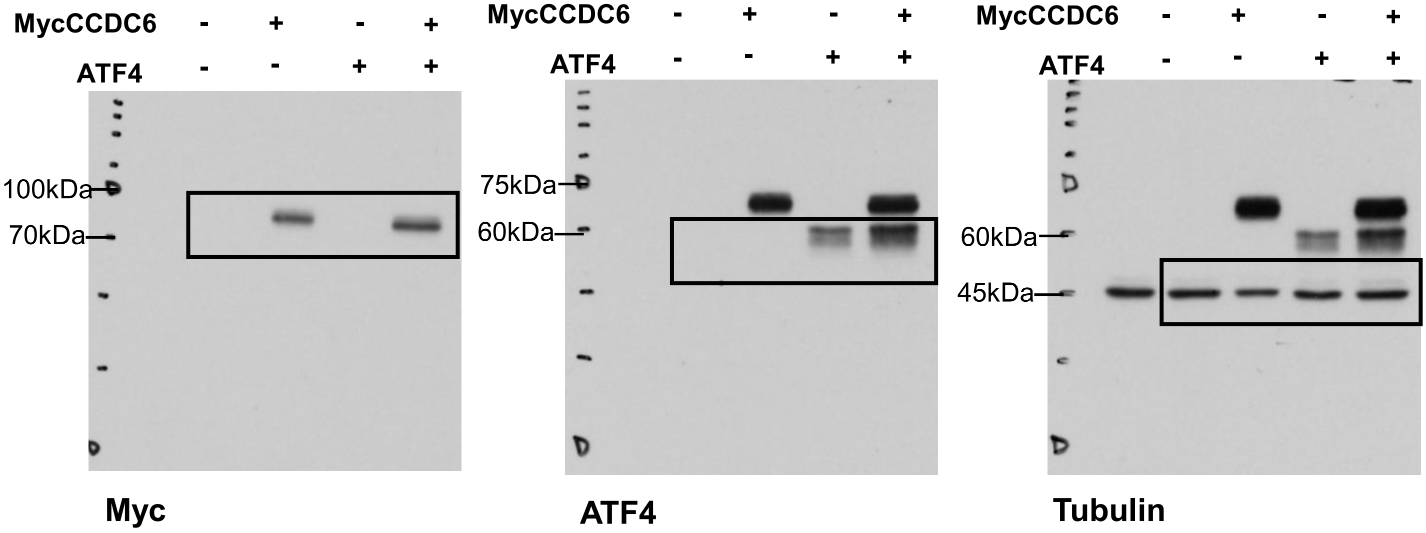

Original Western Blot Relative to Figure 5C

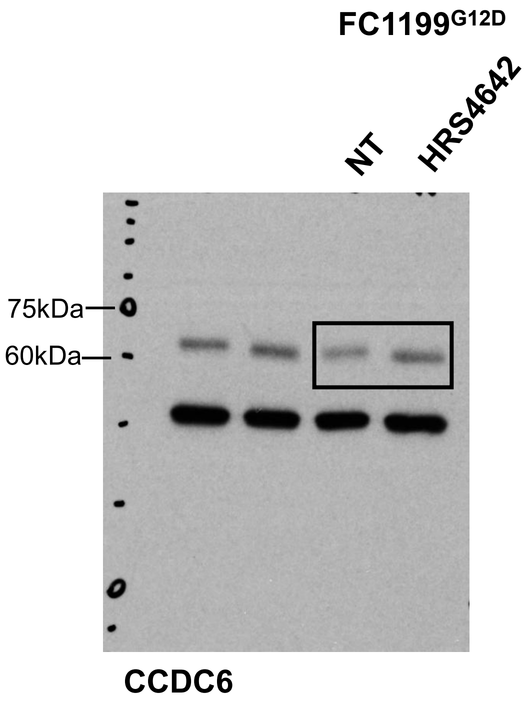


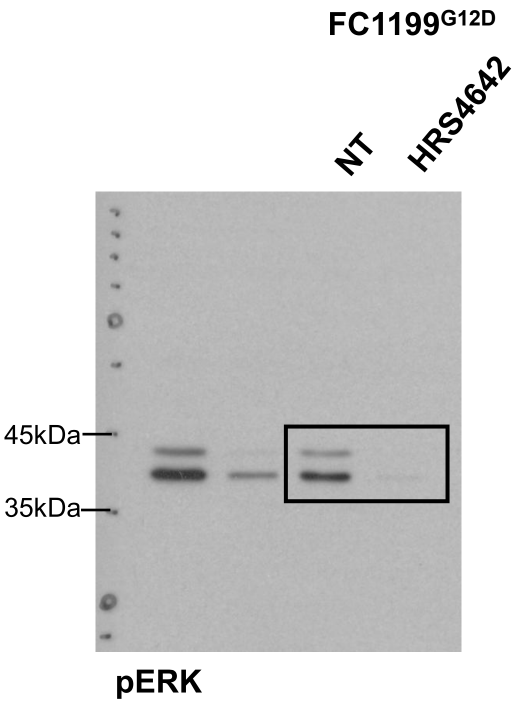


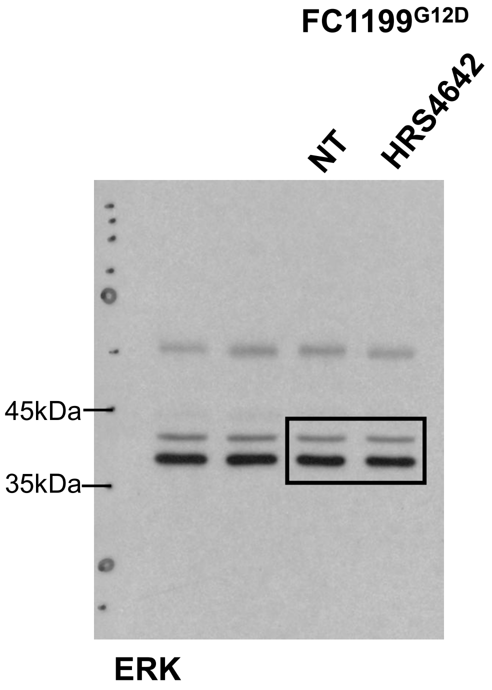


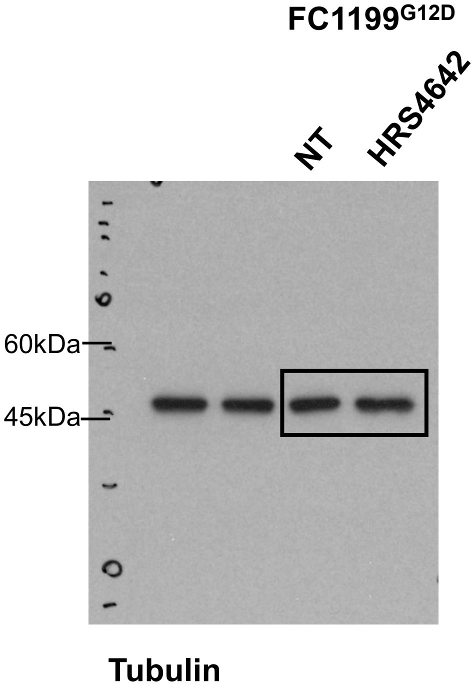

Original Western Blot Relative to Figure S1A

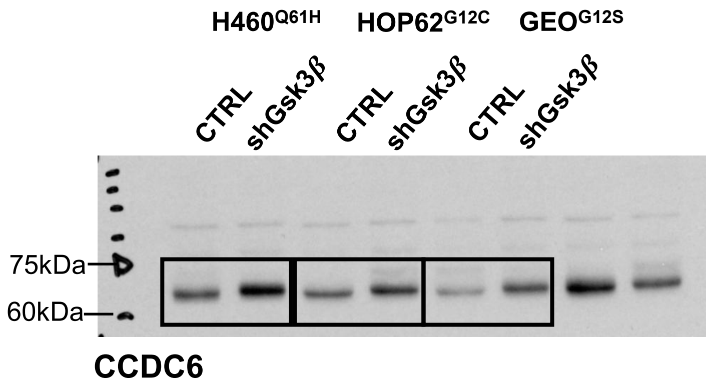


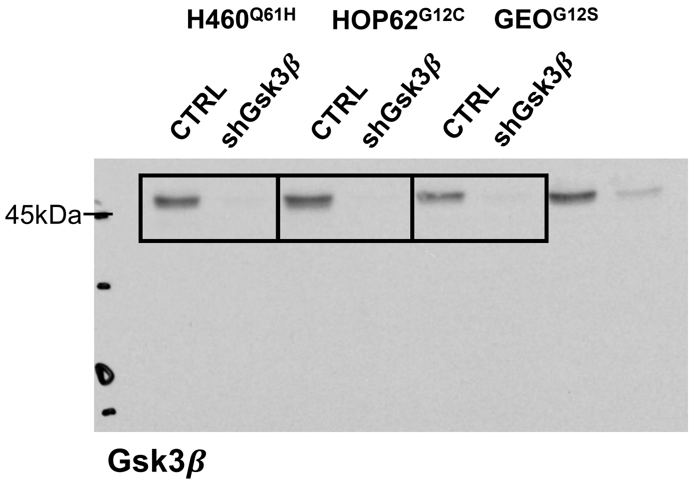


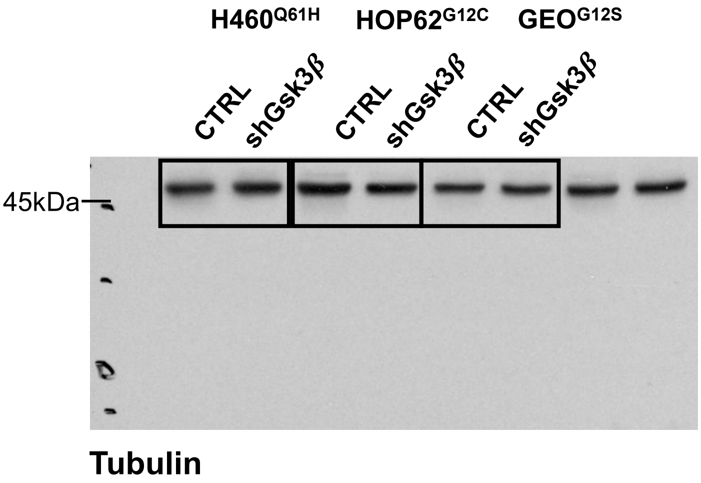

Original Western Blot Relative to Figure S2

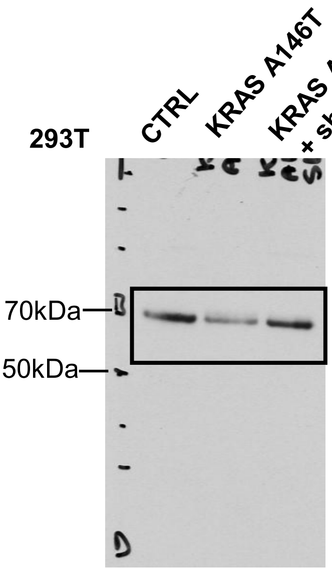


CCDC6

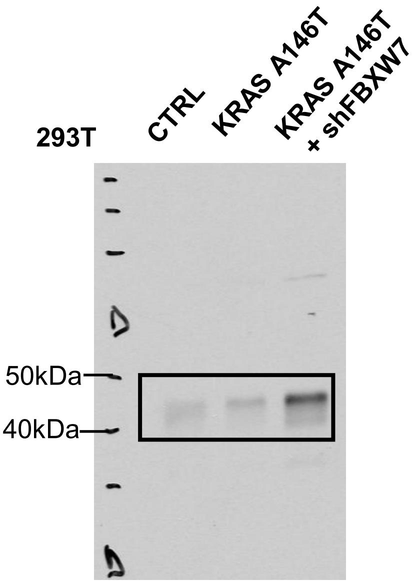


Cyclin E

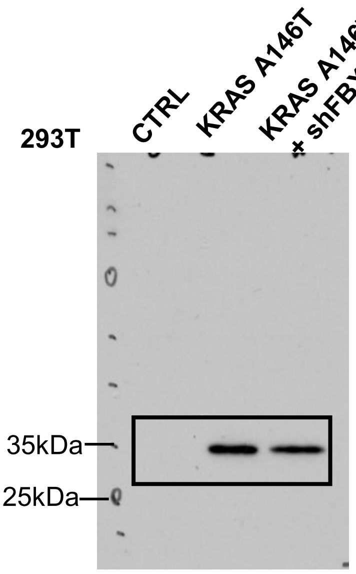


Flag

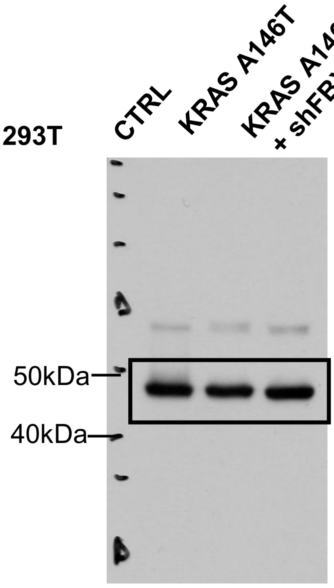


Tubulin
