## Supplementary material for "KRAS-Mediated CCDC6 Degradation Drives xCT Upregulation and Ferroptosis Evasion": Table_S1

**Table S1: Mutational profiling of primary human colorectal cancer (CRC) samples.**

| MUT/WT | PIK3CA | KIT | PDGFRA | BRAF | EGFR | KRAS | NRAS | CCDC6 HSCORE | Quartile |
|---|---|---|---|---|---|---|---|---|---|
| **WT** | WT | WT | WT | WT | WT | WT | WT | 0.6 | Q1 |
| **WT** | WT | WT | V824V | WT | WT | WT | WT | 0.9 | Q1 |
| **WT** | WT | WT | WT | V600E | WT | WT | WT | 1.0 | Q1 |
| **WT** | WT | WT | WT | V600E | WT | WT | WT | 0.3 | Q1 |
| **WT** | E542K | WT | WT | WT | WT | WT | WT | 0.5 | Q1 |
| **WT** | WT | WT | WT | WT | WT | WT | WT | 1.0 | Q1 |
| **WT** | WT | WT | WT | WT | WT | WT | WT | 1.2 | Q1 |
| **WT** | | | | | | WT | WT | 1.5 | Q2 |
| **WT** | WT | WT | WT | WT | WT | WT | WT | 1.5 | Q2 |
| **WT** | WT | WT | WT | WT | WT | WT | WT | 1.7 | Q2 |
| **WT** | WT | WT | WT | WT | WT | WT | WT | 1.7 | Q2 |
| **WT** | WT | WT | WT | WT | WT | WT | WT | 1.5 | Q2 |
| **WT** | WT | WT | WT | WT | WT | WT | WT | 1.6 | Q2 |
| **WT** | WT | WT | WT | V600E | WT | WT | WT | 1.7 | Q2 |
| **WT** | WT | WT | WT | WT | WT | WT | WT | 1.3 | Q2 |
| **WT** | WT | WT | WT | WT | WT | WT | WT | 1.3 | Q2 |
| **WT** | WT | WT | WT | WT | WT | WT | WT | 1.5 | Q2 |
| **WT** | WT | WT | WT | WT | WT | WT | WT | 2.2 | Q3 |
| **WT** | WT | WT | WT | WT | WT | WT | WT | 1.8 | Q3 |
| **WT** | WT | WT | WT | WT | WT | WT | WT | 2.0 | Q3 |
| **WT** | WT | WT | WT | WT | WT | WT | WT | 2.0 | Q3 |
| **WT** | WT | WT | WT | WT | WT | WT | WT | 1.8 | Q3 |
| **WT** | WT | WT | WT | WT | WT | WT | WT | 2.2 | Q3 |
| **WT** | WT | WT | D594G | WT | WT | WT | WT | 2.0 | Q3 |
| **WT** | WT | WT | WT | WT | WT | WT | WT | 2.0 | Q3 |
| **WT** | WT | WT | V824V | WT | WT | WT | WT | 1.8 | Q3 |
| **WT** | WT | M541L | WT | WT | WT | WT | WT | 2.0 | Q3 |
| **WT** | WT | M541L | WT | WT | WT | WT | WT | 2.0 | Q3 |
| **WT** | | | | | | WT | WT | 2.0 | Q3 |
| **WT** | WT | WT | V824V | WT | WT | WT | WT | 2.6 | Q4 |
| **WT** | WT | WT | WT | WT | WT | WT | WT | 2.8 | Q4 |
| **WT** | WT | WT | WT | WT | WT | WT | WT | 3.0 | Q4 |

| | | | | | | | | | |
|---|---|---|---|---|---|---|---|---|---|
| **WT** | WT | M541L | V824V | V600E | WT | WT | WT | 2.3 | Q4 |
| **WT** | Q546R | M541L | WT | WT | WT | WT | WT | 2.3 | Q4 |
| **WT** | WT | WT | WT | WT | WT | WT | WT | 2.4 | Q4 |
| **WT** | WT | WT | WT | WT | WT | WT | WT | 2.5 | Q4 |
| **WT** | WT | WT | WT | WT | WT | WT | WT | 2.55 | Q4 |
| **WT** | WT | WT | WT | WT | WT | WT | WT | 3.0 | Q4 |
| **WT** | WT | WT | WT | WT | WT | WT | WT | 3.0 | Q4 |
| **WT** | G1049R | WT | WT | WT | WT | WT | WT | 2.5 | Q4 |
| **WT** | WT | M541L | WT | V600E | WT | WT | WT | 3.0 | Q4 |
| **MUT** | WT | WT | WT | WT | WT | G13D | | 0.8 | Q1 |
| **MUT** | WT | M541L | WT | WT | WT | G13D | | 0.7 | Q1 |
| **MUT** | WT | M541L | WT | WT | WT | G13D | | 0.7 | Q1 |
| **MUT** | WT | WT | V824V | WT | WT | A146T | | 0 | Q1 |
| **MUT** | E545K | WT | WT | WT | WT | A146T | | 1.2 | Q1 |
| **MUT** | WT | WT | V824V | WT | WT | G12C | | 1.0 | Q1 |
| **MUT** | E542K | WT | V824V | WT | WT | G13D | | 0.5 | Q1 |
| **MUT** | WT | WT | V824V | WT | WT | G12D | | 0.6 | Q1 |
| **MUT** | WT | WT | WT | WT | WT | A146T | | 0.2 | Q1 |
| **MUT** | WT | WT | V828V | WT | WT | G12D | | 0.6 | Q1 |
| **MUT** | WT | WT | WT | WT | WT | G12D | | 1.0 | Q1 |
| **MUT** | WT | WT | WT | WT | WT | G12V | | 0 | Q1 |
| **MUT** | H1047L | WT | WT | WT | WT | G13D | | 1.2 | Q1 |
| **MUT** | E545K | M541L | WT | WT | WT | A146T | | 0.8 | Q1 |
| **MUT** | WT | WT | WT | WT | WT | G12D | | 1.2 | Q1 |
| **MUT** | WT | WT | WT | WT | WT | | Q61K | 1.2 | Q1 |
| **MUT** | WT | WT | WT | WT | WT | G13D | | 0.7 | Q1 |
| **MUT** | WT | WT | WT | WT | WT | G12S | | 0.1 | Q1 |
| **MUT** | WT | WT | WT | WT | WT | G13D | | 0.6 | Q1 |
| **MUT** | WT | WT | V824V | WT | WT | G12D | | 0.6 | Q1 |
| **MUT** | WT | WT | V824V | WT | WT | G13D | | 1.1 | Q1 |
| **MUT** | WT | WT | WT | WT | WT | | Q61K | 1.1 | Q1 |
| **MUT** | H1047R | | | | | G13D | | 0.7 | Q1 |
| **MUT** | / | / | / | / | / | G12D | | 1.1 | Q1 |
| **MUT** | WT | WT | WT | WT | WT | G13D | | 0 | Q1 |

| | | | | | | | | | |
|---|---|---|---|---|---|---|---|---|---|
| **MUT** | WT | WT | WT | WT | WT | G12C | | 1.1 | Q1 |
| **MUT** | WT | WT | WT | WT | WT | G12V | | 0.5 | Q1 |
| **MUT** | WT | WT | WT | WT | WT | G12A | | 1.6 | Q2 |
| **MUT** | WT | WT | V824V | WT | WT | L19F | | 1.4 | Q2 |
| **MUT** | WT | WT | V824V | WT | WT | G12S | | 1.3 | Q2 |
| **MUT** | WT | M541L | WT | WT | WT | Q61K | | 1.7 | Q2 |
| **MUT** | WT | WT | WT | WT | WT | | Q61K | 1.55 | Q2 |
| **MUT** | H1047R | WT | WT | WT | WT | G12V | | 1.6 | Q2 |
| **MUT** | E542K | | V824V | | | G12D | | 1.7 | Q2 |
| **MUT** | WT | M541L | WT | WT | WT | G12V | | 1.7 | Q2 |
| **MUT** | WT | WT | WT | WT | WT | G12D | | 1.3 | Q2 |
| **MUT** | WT | WT | WT | WT | WT | G12V | | 2.1 | Q3 |
| **MUT** | WT | WT | WT | WT | WT | G13D | | 2.0 | Q3 |
| **MUT** | WT | WT | WT | WT | WT | Q61H | | 2.0 | Q3 |
| **MUT** | WT | WT | V824V | WT | WT | G12D | | 2.2 | Q3 |
| **MUT** | WT | WT | WT | WT | WT | G12D | | 2.2 | Q3 |
| **MUT** | WT | WT | WT | WT | WT | G12D | | 2.0 | Q3 |
| **MUT** | WT | WT | WT | WT | WT | | Q61K | 2.0 | Q3 |
| **MUT** | / | / | / | / | / | G12D | | 2.2 | Q3 |
| **MUT** | / | / | / | / | / | Q61H | | 2.0 | Q3 |
| **MUT** | WT | WT | WT | WT | WT | Q61H | | 1.8 | Q3 |
| **MUT** | / | / | / | / | / | G13D | | 2.0 | Q3 |
| **MUT** | / | / | / | / | / | G12D | | 2.0 | Q3 |
| **MUT** | WT | WT | WT | WT | WT | G12V | | 2.0 | Q3 |
| **MUT** | WT | WT | WT | WT | WT | G12V | | 3.0 | Q4 |
| **MUT** | WT | WT | WT | G469R | WT | | G12V | 2.3 | Q4 |
| **MUT** | E545K | M541L | WT | WT | WT | G12D | | 3.0 | Q4 |
| **MUT** | WT | WT | WT | WT | WT | G13D | | 2.6 | Q4 |
| **MUT** | WT | WT | V824V | WT | WT | G12D | | 2.6 | Q4 |
| **MUT** | WT | WT | WT | WT | WT | G12V | | 2.9 | Q4 |
| **MUT** | WT | WT | WT | WT | WT | G12D | | 3.0 | Q4 |
| **MUT** | WT | WT | WT | V600E | WT | G12D | | 3.0 | Q4 |
| **MUT** | E542K | | V824V | | | G12D | | 3.0 | Q4 |
| **MUT** | WT | WT | WT | WT | WT | G12V | | 2.6 | Q4 |

| | | | | | | | | | |
|---|---|---|---|---|---|---|---|---|---|
| **MUT** | E545K | | | | | G12D | | 2.6 | Q4 |

**Table S1. Mutational profiling of primary human colorectal cancer (CRC) samples.**
A total of 101 primary CRC specimens were screened via next-generation sequencing (NGS) for RAS mutational status. Among these, 60 samples harbored RAS mutations, 55 KRAS and 5 NRAS (MUT), while 41 samples were confirmed as RAS wild type (WT). The table details the distribution of specific KRAS and NRAS isoforms, as well as additional co-occurring mutations identified across both WT and mutant cohorts.
